# Nim1-related kinases regulate septin organization and cytokinesis by modulating Hof1 at the cell division site

**DOI:** 10.1101/2025.02.07.634209

**Authors:** Bindu Bhojappa, Anubhav Dhar, VT Bagyashree, Jayanti Kumari, Freya Cardozo, Vaseef Rizvi, Deepthi Guturu, Saravanan Palani

## Abstract

The septin scaffold recruits and organizes actomyosin ring (AMR) components, thus, ensuring faithful cytokinesis. The septin-associated kinases - Elm1, Gin4, Hsl1, and Kcc4 are thought to stabilize and regulate the septin architecture at the bud neck, but the underlying mechanisms remain largely unknown. Here, we present a comprehensive, quantitative analysis of these four septin-associated kinases and reveal major roles for Elm1 and Gin4 in septin stability and architectural transitions during the cell cycle. We find that Elm1 and Gin4 play a previously overlooked role in AMR organization and constriction during cytokinesis. We report that the Gin4 kinase interacts directly with the AMR component and F-BAR protein Hof1 via its C-terminal membrane-binding kinase associated-1 (KA1) domain, and is likely involved in the proper organization and anchoring of Hof1 at the bud neck, representing an unappreciated mode of regulation during cytokinesis. We further show that Gin4 controls septin organization and AMR constriction in a kinase-independent manner, similar to Elm1. Using an extensive GFP-GBP-based tethering assay in *elm1*Δ and *gin4*Δ cells, we identify an important role for Hsl1 in maintaining septin organization and cell shape in coordination with Elm1, Gin4, and Kcc4, independent of its role in the morphogenetic checkpoint. Furthermore, our data indicate that Hsl1 acts downstream of Elm1, with its membrane-binding KA1 domain being critical for its function. Together, these findings reveal new insights into the modes by which the kinases Gin4 and Elm1 regulate cytokinesis, highlight a redundant role for Hsl1 in controlling septin organization and cytokinesis, and uncover the inherent redundancy and adaptability of the septin kinase network in *Saccharomyces cerevisiae*.

## INTRODUCTION

Septins, the fourth cytoskeletal element^1^, are an evolutionarily conserved family of GTP-binding proteins that assemble into filamentous structures on cell membranes^2,3^. Although septins belong to the P-loop GTPases^4^, recent studies have revealed that some septin proteins have lost the ability to hydrolyze bound GTP during evolution^5,6^. They play critical roles in various fundamental processes such as cytokinesis, ciliogenesis, spermatogenesis, and phagocytosis^7–10^. Septins polymerize into hetero-oligomeric complexes that further assemble into higher-order structures, such as filaments, rings, bundles, and gauzes^3,11–15^. These structures function as scaffolds that recruit and organize proteins required for cytokinesis and cell division across eukaryotes^16,17^. Septins act as a diffusion barrier at the cell division site in fungi and animal cells, contributing to the assembly, maturation, and constriction of the contractile AMR machinery during cytokinesis^18–22^. However, studies have suggested that redundant mechanisms can compensate for the septin diffusion barrier, rendering it dispensable for AMR constriction and cytokinesis, indicating that septins may not be strictly essential for AMR function in all contexts^15,17^. The budding yeast *S. cerevisiae* is an excellent model system for studying the relationship between septin organization and cytokinesis. In this organism, five mitotic septins Cdc3, Cdc12, Cdc10, Cdc11, and Shs1^7,23–26^ assemble into an octameric complex that subsequently polymerizes into septin filaments or rods capable of organizing diverse higher-order architectures at the plasma membrane^2,3,27^.

The terminal subunits of the octameric complex influence the architecture of the septin structure: Cdc11 promotes septin filament bundling, whereas Shs1 favours the formation of ring-like structures^28,29^. Septin filaments assemble at the mother-bud neck to form a stable "hourglass" structure that acts as a scaffold for proteins involved in cytokinesis, including components of the AMR complex^14,17,18^. At mitotic exit, this hourglass structure undergoes dynamic remodeling into a double-ring structure, marking the onset of cytokinesis^30–32^. This transition allows proper anchoring of the AMR to the plasma membrane and leads to efficient ring constriction and centripetal septum deposition between the dividing cells^18,33–36^. The mitotic exit network (MEN) is thought to play a crucial role in initiating this transition^30,31^. Despite its importance, the precise molecular mechanisms regulating the hourglass to double-ring (HDR) transition remain incompletely understood.

Several kinases associate with septin structures and regulate septin dynamics during different phases of the cell cycle. These septin-associated kinases localize to the septin hourglass at the beginning of the G1 phase and dissociate from the bud neck at the onset of cytokinesis, coincident with the start of the HDR transition^37–40^. In *S. cerevisiae*, four conserved septin-associated kinases - Elm1^38,39,41–43^, Gin4^38,40,44–50^, Hsl1^51–56^, and Kcc4^57,58^ are thought to exert distinct and overlapping regulatory effects on septin stability and organization^38,39,50,59,60^. Elm1 and Gin4 deletion significantly destabilizes the septin architecture^38,39,41,59^, while loss of Hsl1 and Kcc4 does not have any effect^59^. Several hypotheses have been proposed to describe the functions of these kinases during cytokinesis. Previous studies suggested that the LKB1-like kinase Elm1 and the Nim1-related kinases (Gin4, Hsl1, Kcc4) function in parallel pathways to regulate septin architecture at the bud neck^59,61^. More recently, it was shown that Elm1 acts as the most upstream kinase in this septin kinase network by regulating the activation and phosphorylation of the Nim1-related kinases^39,62^. In addition, Elm1 is known to regulate septin organization by binding to its substrate Bni5 independently of its kinase activity^39^. Gin4, on the other hand, is thought to regulate septin architecture by phosphorylating Shs1^49^. Hsl1 is best known for its role in negatively regulating Swe1 in the morphogenetic checkpoint pathway by forming a complex with Hsl7^52,55,59^. How Hsl1 and Kcc4 regulate septin organization remains unknown, and the mechanisms by which these kinases coordinate septin organization and cytokinesis remain incompletely understood.

Despite significant progress in understanding septin architecture and its regulation, the coordinated effects of septin-associated proteins on this process are only beginning to be understood, and several important questions remain open in the field of septin biology. In particular, it is unclear how septin-associated kinases contribute to the organization and remodeling of septin structures, and how these regulatory pathways are coordinated with AMR assembly and cytokinesis. Furthermore, whether these kinases influence cytokinesis through direct interactions with components of the cytokinetic machinery or indirectly through effects on septin organization remains largely unexplored. In this study, we investigate the roles of the septin-associated kinases Elm1, Gin4, Hsl1, and Kcc4 in regulating septin organization and cytokinesis in *S. cerevisiae*, with the goal of understanding how this kinase network coordinates septin dynamics with AMR function during cell division.

## RESULTS

### Dynamics of septin-associated kinases during the cell cycle and defects associated with their deletions

*S. cerevisiae* contains four major septin-regulatory kinases Elm1, Gin4, Hsl1, and Kcc4 that are known to localize to the mother-bud neck and contribute to septin stability and the morphogenesis checkpoint during the cell cycle^39,59^. Three of these kinases, commonly referred to as Nim1-related kinases^59^: Gin4, Hsl1, and Kcc4 contain a membrane-binding Kinase associated-1 (KA1) domain at the C-terminus **(Fig. 1A)**. The KA1 domain shows affinity for acidic phospholipids (e.g., phosphatidylserine) on the plasma membrane and is required for the bud neck targeting^47,48,51,57^. To characterize their temporal kinetics and localized accumulation at the bud neck septin scaffold, we performed time-lapse imaging of endogenously GFP-tagged septin kinases in cells co-expressing the core septin Cdc3 tagged with mCherry. All four kinases showed simultaneous, gradual recruitment to the septin hourglass during bud emergence and exhibited a sequential pattern of disappearance during the onset of cytokinesis, consistent with previous observations^38,39^ **(Fig. S1A-S1G, Movie-S1)**. Elm1 and Kcc4 exhibited lower fluorescence intensities at the bud neck prior to septin remodeling, suggesting a lower abundance at this site at the defined time period **(Fig. S1E-S1G)**, compared with the higher abundance of Gin4 and Hsl1, which exhibited increased fluorescence intensities prior to septin remodeling **(Fig. S1E-S1G)**. At the onset of cytokinesis, when the septin scaffold undergoes remodeling from the hourglass structure to a double-ring structure (HDR transition), all four kinases left the bud neck just before or during the HDR transition in a sequential manner **(Fig. S1E-S1G, Movie-S1)**. Interestingly, fluorescence intensities of Elm1 and Kcc4 began to decrease ∼4-6 minutes before the HDR transition **(Fig. S1E-S1G)**. In contrast, Gin4 and Hsl1 fluorescence signal intensities began to decrease just before or during the HDR transition, suggesting that they may serve as a trigger or key molecular cue for septin remodeling that subsequently crosstalks with downstream events of cytokinesis **(Fig. S1E-S1G)**. These observations suggest a defined temporal order of removal of these kinases from the bud neck during cytokinesis, which may directly rely on the interdependency of their localizations on one another. Overall, these data show that the four septin kinases are associated with the septin hourglass structure with similar patterns of recruitment but distinct removal kinetics, suggesting that they may play both overlapping and non-overlapping roles in septin hourglass stability and the HDR transition.

**Figure 1:**
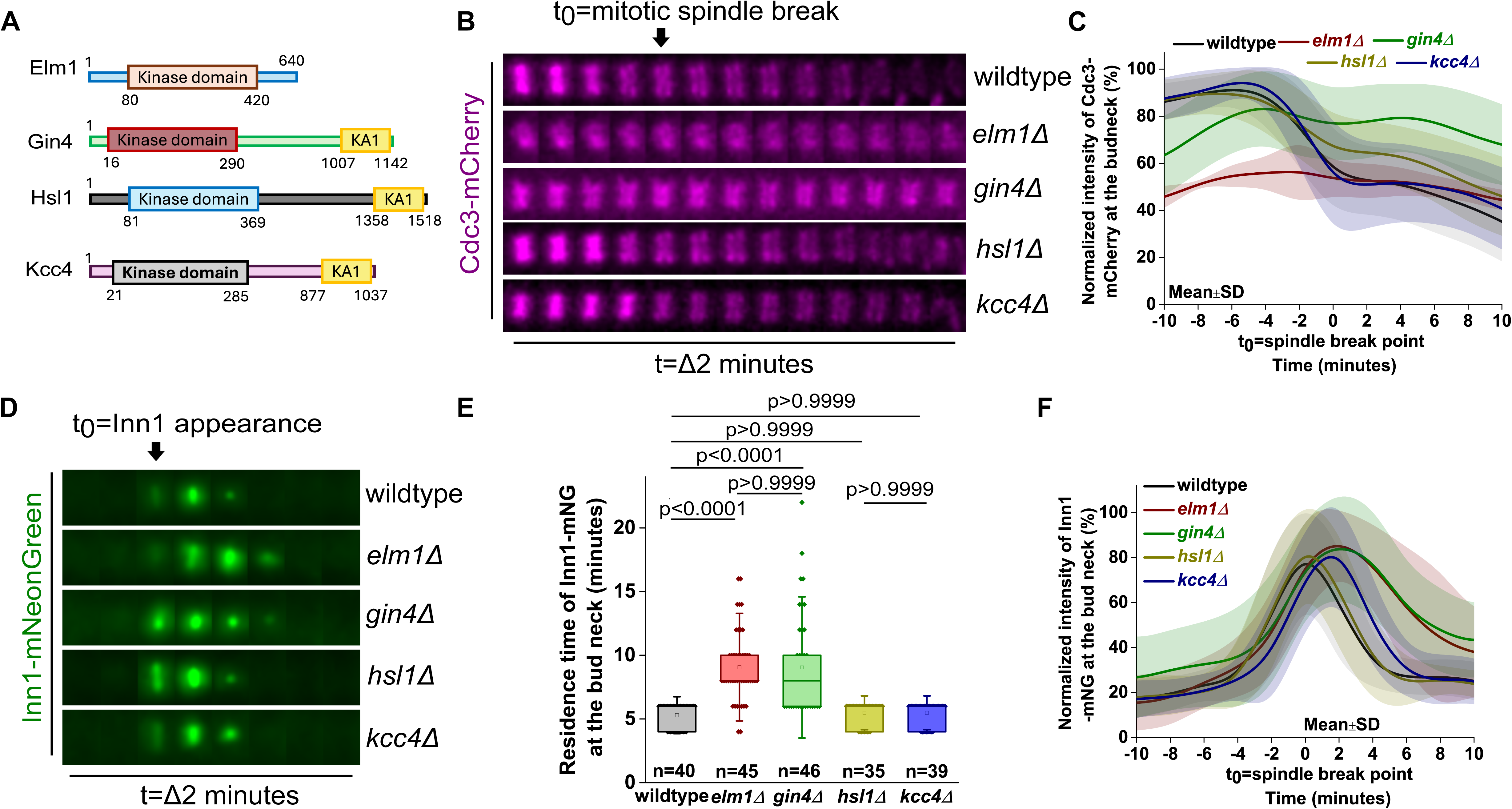
Septin organization and actomyosin ring (AMR) dynamics are defective in *elm1*Δ and *gin4*Δ cells. **(A)** Domain-level architecture of Elm1, Gin4, Kcc4 and Hsl1 kinases. The KA1 domain represents the membrane-binding domain. **(B)** Representative time-lapse images showing Cdc3-mCherry dynamics in septin-kinase deletion strains, captured at two-minute time intervals (t_0_=spindle breakpoint). **(C)** Normalized fluorescence intensity profile showing the temporal kinetics of Cdc3-mCherry in the indicated strains shown in (B) (wild-type: n=41, *elm1*Δ: n=29, *gin4*Δ: n=33, *hsl1*Δ: n=34, *kcc4*Δ: n=36 cells). **(D)** Representative time-lapse images of Inn1-mNG constriction captured at two-minute time intervals in septin-kinase deletion strains (t_0_= appearance of Inn1-mNG at the bud neck). **(E)** Quantification of the residence time of Inn1-mNG during the onset of cytokinesis in the indicated strains of (D), Kruskal-Wallis nonparametric test (****: p<0.0001, ns: p>0.05). **(F)** Normalized fluorescence intensity graph showing the kinetics of Inn1-mNG in the indicated strains of (D). A population of *elm1*Δ and *gin4*Δ cells exhibiting asymmetric constriction is included in the plot (wild-type: n=21, *elm1*Δ: n=13, *gin4*Δ: n=23, *hsl1*Δ: n=21 and *kcc4*Δ: n=18 cells).

### Septin-associated kinases Elm1 and Gin4 majorly regulate septin stability, HDR transition, and cellular morphology

Septin rearrangement during cytokinesis is believed to be partly governed by post-translational modifications (PTMs), including phosphorylation, ubiquitination, acetylation, and SUMOylation^37,45,63–68^. Previously, different studies have highlighted the role of kinase-mediated modifications in septin remodeling, but a comprehensive quantitative analysis for all the four septin-associated kinases under similar conditions is not present^39,49,50,69,70^. Thus, to understand the extent to which different septin kinases affect cell physiology and septin organization, we characterized cell morphology, cell growth, and septin organization in wild-type cells and in cells harbouring deletions of the septin-regulatory kinases - Elm1, Gin4, Hsl1, and Kcc4. We observed that cells exhibited elongated and clumped morphologies in the *elm1*Δ and *gin4*Δ backgrounds, respectively, at all three temperatures (23°C, 30°C, and 37°C) tested, consistent with previous observations^38,41,59,71^ **(Fig. S2A, S2B)**. Quantitative analysis revealed *elm1*Δ cells displayed a more severe defect, with nearly 100% of cells showing elongated morphology that failed to undergo isometric growth, whereas *gin4*Δ cells exhibited clumped morphology in over 73.33% of the cellular population **(Fig. S2A, S2B)**. While *hsl1*Δ cells largely showed round morphology similar to wild-type at 23°C and 30°C, a fraction of cells exhibited mild morphological defects at 37°C **(Fig. S2A, S2B)**. In contrast, no alterations in morphology were observed in *kcc4*Δ cells **(Fig. S2A, S2B)**. Growth assays of these kinase deletions showed a similar trend: *elm1*Δ and *gin4*Δ cells showed growth sensitivity at both 23°C and 30°C, which was heightened at 37°C for *elm1*Δ, while *hsl1*Δ exhibited growth sensitivity only at the higher temperature 37°C, in contrast to *kcc4*Δ cells, which grew identical to the wild-type strain **(Fig. S2C)**.

Next, we sought to systematically assess the effects of the absence of these kinases on septin architecture and dynamics. The core septin Cdc3 fused to mCherry was used to visualize the septin structures during the cell cycle^72^. Cdc3-mCherry localized to the emerging bud site in wild-type, *elm1*Δ, *gin4*Δ, *hsl1*Δ, and *kcc4*Δ cells but was subsequently mislocalized to the bud cortex, consistent with previous findings^39,41,50,61^ in ∼100% of *elm1*Δ cells and 80.48% of *gin4*Δ cells **(Fig. S2E, S2G, Movie-S2)**. In wild-type cells, septin formed an stable hourglass structure at the bud neck, which subsequently split into double rings coinciding with spindle breakage and mitotic exit **(Fig. S2D, S2F)**. The septin hourglass architecture appeared abnormal in *elm1*Δ and *gin4*Δ cells and exhibited a defective HDR transition **(Fig. 1B, 1C, Movie-S2)**. The *elm1*Δ cells showed a stronger defect in septin organization during bud emergence and the HDR transition compared with *gin4*Δ cells **(Fig. 1B, 1C, S2E, S2G)**. This suggests a more prominent role for Elm1 in the maintenance of septin filament architecture, consistent with previous literature reporting a role for Elm1 in stabilizing the daughter-side of the septin hourglass structure by regulating septin filament pairing^39^. Cells lacking the Hsl1 and Kcc4 kinases showed normal septin organization during bud emergence and HDR transition at the onset of cytokinesis, similar to wild-type cells **(Fig. 1B, 1C, S2E, S2G)**. However, approximately 7.14% of *hsl1*Δ cells displayed septin mislocalization to the bud cortex during bud emergence **(Fig. S2E, S2G)**, indicating that Hsl1 may also play a minor or supporting role in septin dynamics through a compensatory mode of regulation. These results therefore demonstrate a major requirement for Elm1 and Gin4 in proper septin arrangement and dynamics at the cell division site, while also hinting at a parallel minor role for Hsl1.

### Elm1 and Gin4 regulate AMR organization and constriction

Given that our data suggest major defects in septin regulation occur in the absence of Elm1 and Gin4, we asked whether these defects also translate into effects on AMR-mediated cytokinesis. We therefore investigated the roles of Elm1 and Gin4 in AMR organization at the cell division site and in constriction dynamics during cytokinesis. The AMR machinery generates the force required to drive plasma membrane ingression for successful cell separation^18,73^. In budding yeast, the AMR machinery consists primarily of actin filaments, tropomyosin, formins, Myo1, Hof1, Iqg1, Cyk3, and Inn1, along with many other regulatory proteins^18,21,74–76^. We used endogenously labelled Myo1-mNeonGreen (mNG) the sole non-muscle myosin-II in *S. cerevisiae* to determine AMR organization during bud emergence. Previous studies have shown that Myo1 recruits to the bud neck in a bi-phasic manner - via Bni5 prior to cytokinesis and via Iqg1 during cytokinesis^72,77^. Consistent with its interaction with septin-binding protein Bni5^72^, we observed that, similar to Cdc3-mCherry, Myo1-mNG was mislocalized to the bud cortex during bud emergence in 75.86% and 30% of *elm1*Δ and *gin4*Δ cells, respectively **(Fig. S3A, S3B)**. We then used Inn1 which arrives two-minutes after the start of AMR constriction and aids in plasma membrane ingression by coupling AMR constriction with primary septum (PS) formation as a marker to determine AMR constriction dynamics during cytokinesis^75,76^. In wild-type cells, Inn1-mNG arrived at the bud neck just prior to spindle breakage and constricted symmetrically as part of the AMR complex with a residence time of ∼5-6 minutes, and eventually disappeared from the bud neck **(Fig. 1D-1F)**. In *elm1*Δ and *gin4*Δ cells, Inn1-mNG was recruited normally to the cell division site but displayed an abnormal constriction pattern with an increased duration of constriction (∼8-10 minutes) **(Fig. 1D-1F, Movie-S3)**. By contrast, in *hsl1*Δ and *kcc4*Δ cells, Inn1-mNG was recruited normally and constricted similarly to the wild-type strain **(Fig. 1D-1F, Movie-S3)**. These results agree with the regulation of these kinases on septin dynamics, suggesting that Elm1 and Gin4 may directly or indirectly regulate AMR constriction and cytokinesis.

Inn1 interacts with Hof1, an F-BAR protein known to associate with the septin hourglass, transit to the AMR during the split-ring trigger^73,74,78–82^, and sense membrane curvature^74,76,82^. Inn1 also functions in parallel with Cyk3; Hof1, Inn1 and Cyk3 form a tripartite complex referred to as the Ingression Progression Complex (IPC)^76,83–85^. This complex then activates the primary component of the PS, Chitin Synthase II (Chs2)^76,83–85^. Thus, we monitored the constriction pattern and dynamics of Chs2-mNG and observed an increased residence time for Chs2 at the bud neck in both *elm1*Δ and *gin4*Δ cells **(Fig. S3C-S3E)**. Our observations suggest that loss of the septin kinases Elm1 and Gin4 may result in improper organization and constriction dynamics of the AMR, and possibly impaired IPC formation, resulting in abnormal cytokinesis.

To test the organization of the AMR/IPC at the bud neck, we assessed the organization of the F-BAR protein Hof1. Deletion of Hof1 has been shown to result in asymmetric AMR constriction and improper chitin deposition during PS synthesis^76^. We then analyzed the residence time of Hof1-mNG at the bud neck after the spindle breakpoint during cytokinesis. Hof1-mNG exhibited a residence time of 10-12 minutes after the mitotic spindle break in wild-type cells, which increased to >15 minutes in both *elm1*Δ and *gin4*Δ cells **(Fig. 2A, 2B, Movie-S4)**. By contrast, we did not observe any significant change in the recruitment and temporal kinetics of Hof1-mNG in *elm1*Δ and *gin4*Δ cells compared with wild-type during the mitotic spindle break **(Fig. 2C)**. Next, we analyzed the organization of Hof1-mNG at the cell division site. In wild-type cells, Hof1-mNG appeared as a continuous, uniform ring at the bud neck in 93.05% of the cellular population, whereas discontinuous or fragmented Hof1-mNG rings were observed in 82.69% and 71.07% of *elm1*Δ and *gin4*Δ cells, respectively **(Fig. 2D, 2E)**. These observations suggest that Elm1 and Gin4 regulate the proper spatial assembly of various AMR components to ensure robust cytokinesis. The effect on AMR dynamics observed in *elm1*Δ and *gin4*Δ cells may be an indirect consequence mediated through septins, because septin organization is significantly defective in these backgrounds. However, a previous study has shown that disruption of the septin scaffold does not affect cytokinesis, suggesting that Elm1 and Gin4 could regulate AMR organization and constriction independently of their effects on septin organization^17^. Together, these results reveal an unappreciated role for Elm1 and Gin4 in AMR assembly, organization, and constriction, opening avenues for future investigation.

**Figure 2:**
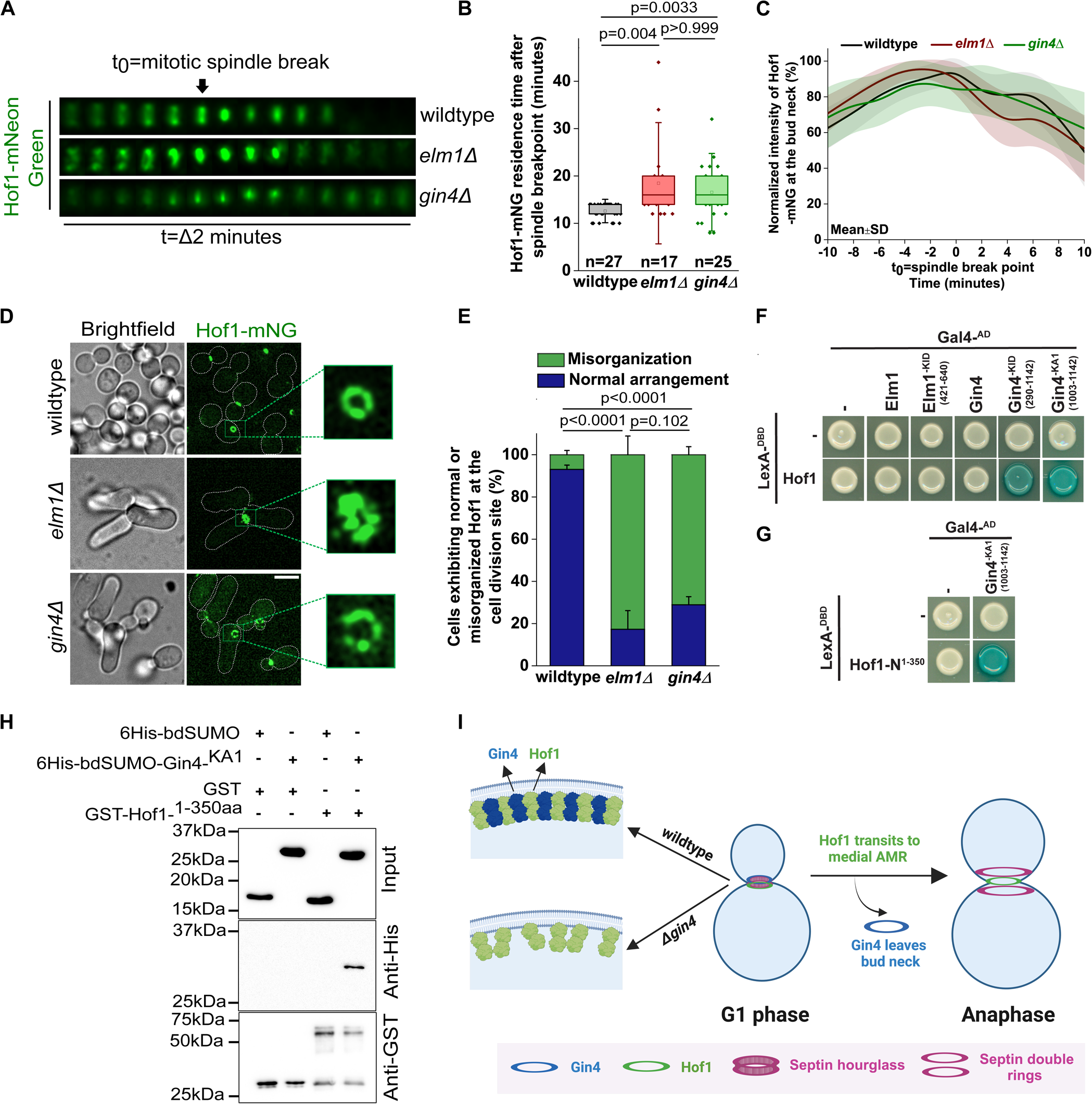
Gin4 regulates Hof1 organization and dynamics through interaction with the N-terminal membrane-binding F-BAR domain. **(A)** Representative time-lapse images of Hof1-mNG in wild-type, *elm1*Δ and *gin4*Δ cells imaged at two-minute time intervals (t_0_=spindle breakpoint). **(B)** Quantification of the residence time of Hof1-mNG after spindle breakpoint in the indicated strains shown in (A), Kruskal-Wallis nonparametric test (**: p<0.01, ns: p>0.05). **(C)** Normalized fluorescence intensity profile for the temporal kinetics of Hof1-mNG during cytokinesis in the indicated strains shown in (A), (wild-type: n=33, *elm1*Δ: n=21, *gin4*Δ: n=23 cells). **(D)** Representative images showing the organization of Hof1-mNG at the bud neck in the indicated strains shown in (A). Scale bar-5µm. **(E)** Bar graph showing the percentage of cells exhibiting misorganization of Hof1 at the cell division site in the indicated strains shown in (D), one-way ANOVA with Tukey’s multiple-comparison test (****: p<0.0001, ns: p>0.05). p-values correspond to the population of cells exhibiting disorganized Hof1-mNG at the bud neck. (N=3, wild-type: n=518, *elm1*Δ: n=225, *gin4*Δ: n=384 cells). **(F)** Representative images from the Yeast Two-Hybrid assay depicting the interaction of full-length Gin4, Gin4-KID and Gin4-KA1 with full-length Hof1. **(G)** Representative Yeast Two-Hybrid images showing the interaction between Gin4-KA1 (1003-1142aa) and N-terminal F-BAR containing domain of Hof1 (1-350aa). **(H)** Immunoblot showing the in vitro binding assay for the interaction between 6xHis-bdSUMO-Gin4^KA1^ and GST-Hof1^F-BAR^ fragments. GST-tagged Hof1 fragment is immobilized on glutathione resin, and bound 6xHis-bdSUMO-tagged Gin4 fragments were analyzed by SDS PAGE. Input (top) and pull-down fractions (middle) were probed with anti-His antibody, while GST-Hof1 fragments (bottom) were detected using anti-GST antibody, (+) present (–) absent. **(I)** Proposed model depicting the possible interplay between Gin4 and Hof1 at the plasma membrane interface.

### Gin4 directly interacts with the N-terminal F-BAR domain of Hof1 to regulate proper cytokinesis

To understand how the Elm1 and Gin4 kinases affect AMR dynamics, we performed a Yeast Two-Hybrid (Y2H) assay to identify novel interacting partners of these proteins. We found that two Gin4 fragments - Kinase Independent Domain (KID; 290-1142 aa) and Kinase associated-1 (KA1) (1003-1142 aa) but not the full-length protein, interacted with the full-length Hof1 protein **(Fig. 2F)**. Further, the N-terminal Hof1 fragment (1-350 aa), which contains its F-BAR domain, but not the other fragments, interacted with the Gin4-KA1 domain **(Fig. 2G, S3H)**. By contrast, neither full-length Elm1 nor its C-terminal fragment (421-640 aa) showed any interaction with Hof1 in our Y2H assay **(Fig. 2F)**. However, the Y2H assay for detecting protein-protein interactions has inherent limitations, including a bias toward high-affinity interactions and susceptibility to false-positive detection^86^. As a result, biologically relevant interactions may be missed, particularly when one or both interacting partners are membrane-associated, self-activating, require post-translational modifications, or interact indirectly through adaptor proteins^86^. Given these limitations of the Y2H approach, we performed an in-vitro binding assay between the Gin4^KA1^ domain and the N-terminal region of Hof1^N (1–350 aa)^ to independently validate the direct interaction between these proteins.

We purified 6His-bdSUMO-Gin4-KA1 (1003-1142 aa) and GST-Hof1-N-terminus (1-350 aa) fragments from *E. coli*. Strikingly, our results revealed that the N-terminal, membrane-binding, F-BAR-containing domain of Hof1 (1-350 aa) interacts strongly with the C-terminal membrane-binding KA1 domain of Gin4 (1003-1142 aa) **(Fig. 2H)**. The Hof1 N-terminal F-BAR fragment did not pull down the control 6His-bdSUMO fragment but specifically pulled down 6His-bdSUMO-Gin4^KA1^ under identical experimental conditions (**Fig. 2H**), confirming the specificity of the interaction between Hof1 F-BAR domain and Gin4-KA1.

Overall, these results reveal a direct physical link between the Gin4 and the AMR component Hof1, suggesting a novel and direct mode of regulation of cytokinesis by the septin kinase network in *S. cerevisiae*. The involvement of the membrane-binding interfaces of these proteins also suggests a possible coordinative mechanism in which Gin4 may regulate Hof1 to control its proper assembly and anchoring to the membrane prior to cytokinesis **(Fig. 2I)**. This is supported by the disorganized and irregular Hof1 ring at the cell division site observed in *gin4*Δ cells **(Fig. 2D, 2E)**. Moreover, *gin4*Δ cells also show asymmetric AMR constriction **(Fig. 2A)**, which is a known characteristic phenotype of *hof1*Δ cells^76^. Thus, our data, combined with these line of evidences, strongly suggest that Gin4 exerts direct control over AMR dynamics by physically interacting with Hof1.

To further understand the direct role of Gin4’s membrane-binding KA1 domain in cytokinesis and Hof1 dynamics, we generated an endogenously tagged Gin4-*ka1*Δ-GFP strain and compared its localization kinetics with those of Gin4-GFP during mitotic spindle breakdown. We observed a marked reduction in Gin4-*ka1*Δ-GFP localization at the bud neck during cytokinesis, suggesting that the KA1 domain contributes to the recruitment of Gin4 **(Fig. S4A-S4C)**. In addition, we analyzed the temporal kinetics of Hof1-mNG at the bud neck in both *gin4*Δ and Gin4-*ka1*Δ cells. Consistent with previous findings by Moravcevic et al. (2010), the Gin4-*ka1*Δ mutant phenocopied the *gin4*Δ deletion, exhibiting a clumped morphology^57^. In both *gin4*Δ and Gin4-*ka1*Δ cells, Hof1-mNG exhibited a comparable constriction pattern after mitotic spindle breakdown, extending beyond 20 minutes **(Fig. S4D, S4E)**. Additionally, Hof1-mNG exhibited similar localization kinetics during cytokinesis in *gin4*Δ and Gin4-*ka1*Δ cells **(Fig. S4D, S4F)**. Taken together, these results strongly suggest that the Gin4-KA1 domain is indispensable for Gin4 function and may directly regulate AMR maturation and constriction by influencing Hof1 dynamics at the cell division site.

### Gin4 exerts its control over septin organization and AMR components in a kinase-independent manner

Elm1 kinase activity is essential for its localization to the bud neck but is dispensable for its role in septin stability^39^. Bni5 is a myosin-recruiting protein that stabilizes and crosslinks septin filaments by binding to the terminal septin Cdc11^72,87–89^. Elm1 regulates the septin hourglass structure and stability through its binding partner Bni5, independent of its kinase activity^39^. We therefore asked whether Gin4 kinase activity is required for its functions in the regulation of septin organization and cytokinesis. To address this, we complemented *gin4*Δ cells with the kinase-dead *gin4-K48A* allele^38,44,90^ (henceforth referred to as Gin4^KD^), integrated into the *leu2* locus. Interestingly, Gin4^KD^-GFP localized to the bud neck, similar to endogenously tagged Gin4 and Gin4^FL^-GFP, and complemented the septin mislocalization defects during bud emergence in 78% of total *gin4*Δ cells **(Fig. 3A, 3B)**. Additionally, expression of Gin4^KD^-GFP in *gin4*Δ cells restored the residence time of Inn1-mNG at the bud neck to ∼5-7 minutes and the temporal kinetics during cytokinesis, similar to the wild-type strain **(Fig. 3C-3E)**. Furthermore, Gin4^KD^ restored Hof1 misorganization defects at the cell division site when expressed in *gin4*Δ cells in 77.14% of the cellular population **(Fig. S3F, S3G)**. Gin4 kinase is known to stabilize septins by phosphorylating its substrate, Shs1^49^. Hof1 has been shown to exhibit synthetic lethality with Gin4, possibly by regulating septins in a parallel pathway^74^. To test whether this synthetic lethality depends on the kinase activity of Gin4, we expressed Gin4^FL^ and Gin4^KD^ in cells carrying the *gin4*Δ *hof1*Δ double deletion **(Fig. S3I)**. Interestingly, Gin4^KD^ rescued the synthetic lethality, similar to Gin4^FL^, suggesting that the kinase activity of Gin4 is not required to restore viability in the absence of Hof1 **(Fig. S3I)**. These findings provide a strong evidence that Gin4 kinase activity is largely dispensable for its localization and for its functions in septin organization and cytokinesis, addressing an important aspect of how Gin4 may perform these functions in the absence of catalysis while raising several new questions.

**Figure 3:**
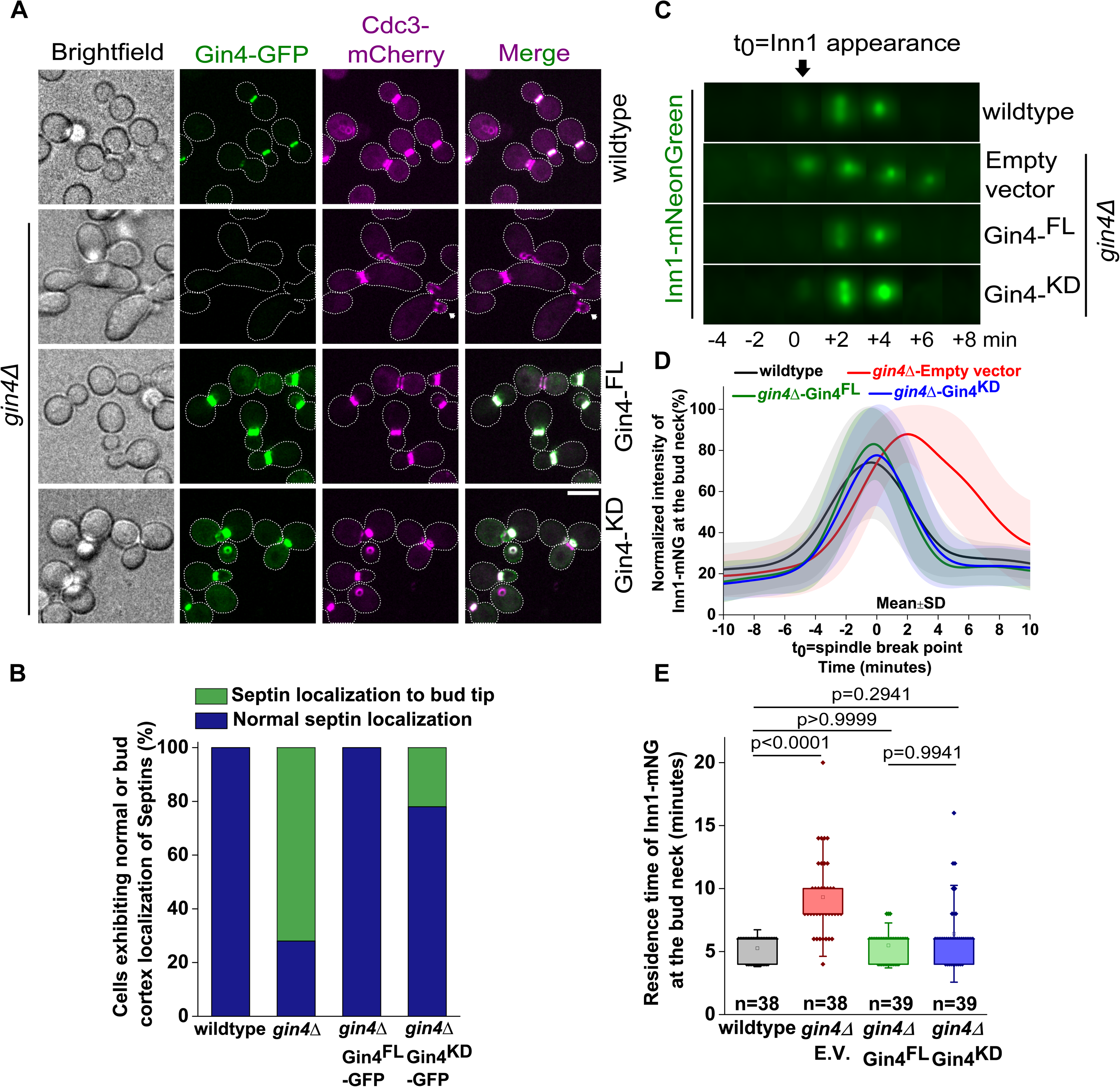
Gin4 controls septin organization and AMR dynamics independently of its kinase activity. **(A)** Representative images showing septin defects or mislocalization during bud emergence in *gin4*Δ cells and rescue of the mislocalization by Gin4^FL^-GFP and Gin4^KD^-GFP constructs expressed under pTEF promoter. White arrows indicate mislocalized Cdc3-mcherry in the represented strains. Scale bar-5µm. **(B)** Quantitation of septin mislocalization and its rescue in the indicated strains shown in (A) (wild-type: n=38, *gin4*Δ: n=32, *gin4*Δ-Gin4^FL^-GFP: n=32, *gin4*Δ-Gin4^KD^-GFP: n=36 cells). **(C)** Representative time-lapse images showing Inn1-mNG dynamics acquired at two-minute time intervals in *gin4*Δ cells expressing Gin4^FL^ and Gin4^KD^ constructs cloned under endogenous promoter (t_0_=appearance of Inn1-mNG at the bud neck). **(D)** Plot of normalized fluorescence intensity of Inn1-mNG in indicated strains shown in (C). A population of *gin4*Δ-Empty vector cells exhibiting asymmetric constriction is plotted in the graph (t_0_=spindle breakpoint) (wild-type: n=38, *gin4*Δ-Empty vector: n=21, *gin4*Δ*-* Gin4^FL^: n=39, *gin4*Δ*-* Gin4^KD^: n=31 cells). **(E)** Quantification of the residence time of Inn1-mNG during cytokinesis in indicated strains of (C), Kruskal-Wallis nonparametric test (****: p<0.0001, ns: p>0.05).

### Hsl1 kinase can partially compensate for Elm1 function upon forced tethering to the bud neck

Marquardt et al. (2024) showed that Gin4 and Elm1 exhibit reciprocal regulation during different cell cycle stages, and that their localization to the bud neck is interdependent^38^. The study also reported that Gin4 recruitment and localization to the bud neck is hampered as the cell cycle progresses in *elm1*Δ cells^38^. We independently corroborated these results and observed that Gin4 was mislocalized to the bud cortex during bud emergence **(Fig. S5A)** and exhibited defective localization to the bud neck during later cell cycle stages in *elm1*Δ cells **(Fig. S5B-S5D)**. Next, we asked whether restoring Gin4 localization to the bud neck could rescue the elongated morphology in cells lacking Elm1. To test this hypothesis, we performed an artificial tethering assay using the GFP-GBP strategy **(Fig. 4A)**. The GFP-Binding Protein (GBP) is a ∼13 kDa soluble protein derived from the heavy chain of camelid (*llama*) antibodies^91^. This single-domain antibody binds GFP-tagged proteins with very high affinity, in the picomolar range. GBP also binds other GFP variants, such as Yellow Fluorescent Protein (YFP), with a strict 1:1 stoichiometric ratio^92^. By contrast, GBP does not bind to Cyan Fluorescent Protein (CFP) or to derivatives of DsRed, including monomeric Red Fluorescent Protein (mRFP) and mOrange^92^. Owing to its high binding affinity and specificity, the GFP-GBP system was initially developed for the purification of GFP-tagged proteins^91,92^. More recently, this strategy has been adapted to artificially redirect proteins to non-canonical subcellular locations^38,91,92^. Based on this approach, we tagged the terminal septin Shs1^49,93^ and the bud neck-localizing septin-associated proteins (SAPs) the anillin-like protein - Bud4^94^, Bni5^39^, and Hsl1 with GBP at their C-termini. These GBP-tagged proteins were then used to redirect mislocalized Gin4-GFP to the bud neck, allowing us to assess whether forced relocalization of Gin4 could rescue the elongated morphology exhibited by *elm1*Δ cells.

**Figure 4:**
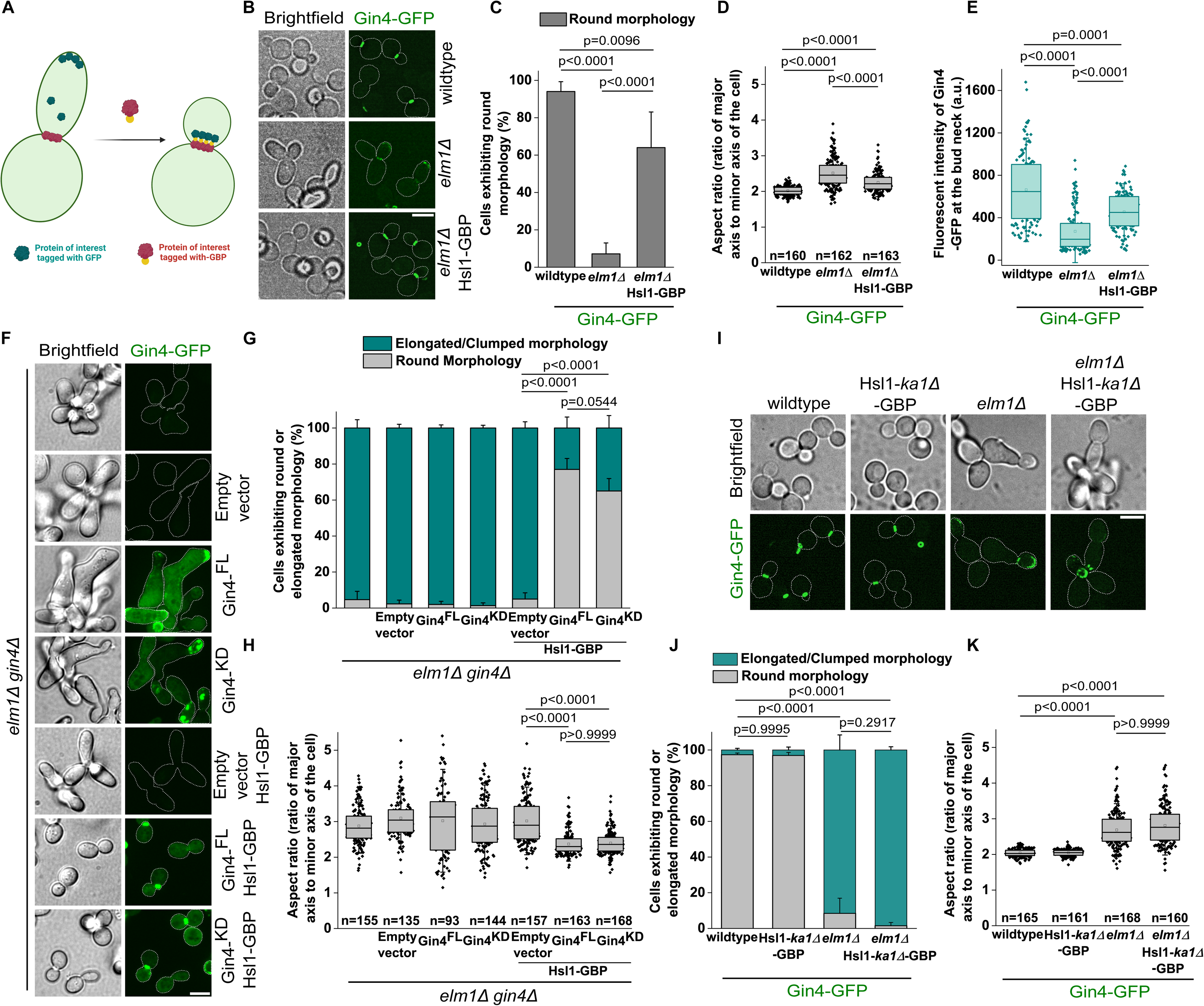
Artificial tethering of Gin4-GFP to the bud neck via Hsl1-GBP restores functionality in *elm1*Δ cells in a Gin4 kinase-independent manner. **(A)** Schematic representation of GFP-GBP artificial tethering strategy, in which the protein of interest is targeted the bud neck by C-terminal GFP tagging, while the binding partner is tagged with GBP. **(B)** Representative images showing artificial tethering of Gin4-GFP with Hsl1-GBP in *elm1*Δ cells. Scale bar-5µm. **(C)** Bar graph showing the percentage of cells exhibiting round morphology in the indicated strains shown in (B), one-way ANOVA with Tukey’s multiple-comparison test (**: p<0.01, ****: p<0.0001), (N=3, wild-type: n=407, *elm1*Δ: n=508, *elm1*Δ-Hsl1-GBP: n=400 cells). **(D)** Quantification of aspect ratios in the indicated strains shown in (B), Kruskal-Wallis nonparametric statistical test (****: p<0.0001), (N=3, n>150 cells/strain). **(E)** Graph depicting the raw fluorescence intensity of Gin4-GFP at the large bud in the indicated strains shown in (B), Kruskal-Wallis nonparametric statistical test (***: p<0.001, ****: p<0.0001) (N=3, wild-type: n=125, *elm1*Δ: n=125, *elm1*Δ-Hsl1-GBP: n=118 cells). **(F)** Representative images showing the localization of Gin4^FL^-GFP and Gin4^KD^-GFP constructs expressed under the pTEF promoter and their tethering to bud neck with Hsl1-GBP in *elm1*Δ *gin4*Δ double-deletion strains. Scale bar-5µm. **(G)** Stacked column graph showing the percentage of cells exhibiting round and elongated/clumped morphologies in the indicated strains shown in (F), one-way ANOVA with Tukey’s multiple-comparison test (****: p<0.0001, ns: p>0.05). p-values correspond to population of cells exhibiting round morphology (N=3, *elm1*Δ *gin4*Δ: n=339, *elm1*Δ *gin4*Δ-Empty vector: n=327, *elm1*Δ *gin4*Δ- Gin4^FL^: n=173, *elm1*Δ *gin4*Δ- Gin4^KD^: n=236, *elm1*Δ *gin4*Δ-Empty vector-Hsl1-GBP: n=356, *elm1*Δ *gin4*Δ-Gin4^FL^-Hsl1-GBP: n=365, *elm1*Δ *gin4*Δ-Gin4^KD^-Hsl1-GBP: n=459 cells). **(H)** Quantification of aspect ratios in the indicated strains shown in (F), Kruskal-Wallis nonparametric statistical test (****: p<0.0001, ns: p>0.05) (N=3, n>90 cells/strain). **(I)** Representative images showing artificial tethering of Gin4-GFP with Hsl1-*ka1*Δ-GBP in *elm1*Δ cells. Scale bar-5µm. **(J)** Stacked column graph showing the percentage of cells exhibiting round and elongated/clumped morphologies in the indicated strains shown in (I), one-way ANOVA with Tukey’s multiple comparison test (****: p<0.0001, ns: p>0.05) p-values correspond to the population of cells exhibiting round morphology (N=3, wild-type: n=674, Hsl1-*ka1*Δ-GBP: n=419, *elm1*Δ: n=572, *elm1*Δ Hsl1-*ka1*Δ-GBP: n=337 cells). **(K)** Quantification of aspect ratios in the indicated strains shown in (I), Kruskal-Wallis nonparametric statistical test (****: p<0.0001, ns: p>0.05), (N=3, n>150 cells/strain).

Forced tethering of Gin4-GFP to the bud neck via its related kinase Hsl1-GBP restored normal cellular morphology in 63.75% of the population in *elm1*Δ cells **(Fig. 4B, 4C, Table 1)**. To quantitatively assess this rescue, we also analyzed the aspect ratio, measured as the ratio of the major axis to the minor axis of the cell. As expected, the mean aspect ratio was higher in *elm1*Δ cells, than in wild-type cells, reflecting their elongated morphology **(Fig. 4D)**, and decreased in cells where Gin4-GFP was tethered to the bud neck with Hsl1-GBP, corroborating the rescue of cell morphology observed after restoring Gin4-GFP localization to the bud neck **(Fig. 4D)**. We measured fluorescence intensities of Gin4-GFP with and without tethering to the bud neck via Hsl1-GBP in large-budded *elm1*Δ cells, which validated that Gin4-GFP was indeed tethered to the bud neck **(Fig. 4E)**. In contrast to the rescue observed after tethering with Hsl1-GBP, artificial tethering of Gin4-GFP via other bud neck proteins Shs1, Bni5, or Bud4 did not rescue the elongated phenotype of *elm1*Δ cells, despite restoration of bud neck localization **(Fig. S5E-S5G)**. These results suggest that the observed rescue may be due to two underlying causes: (i) restoration of Gin4 protein at the bud neck, and (ii) tethering of Gin4 to Hsl1.

**Table 1:** Table summarizing the rescue of elongated or clumped morphology upon redirecting the Nim1-related kinases to the bud neck in *elm1*Δ cells and Elm1-GFP in *gin4*Δ cells using GFP-GBP artificial tethering strategy.

| Protein tagged with GFP | Protein tagged with GBP | Genetic background | Number of cells exhibiting round morphology | Number of cells exhibiting elongated/clumped morphology | Total number of cells | Percentage of cellular population exhibiting round morphology | Rescue of elongated or clumped morphology |
| --- | --- | --- | --- | --- | --- | --- | --- |
| Gin4 | Shs1 | *elm1* $\Delta$ | 24 | 464 | 488 | 4.91 | No |
| Gin4 | Bud4 | *elm1* $\Delta$ | 20 | 451 | 471 | 4.24 | No |
| **Gin4** | **Hsl1** | ***elm1* $\Delta$** | **255** | **145** | **400** | **63.75** | **Yes** |
| Gin4 | Hsl1-*ka1* $\Delta$ | *elm1* $\Delta$ | 6 | 331 | 337 | 1.78 | No |
| Gin4 | Bni5 | *elm1* $\Delta$ | 17 | 431 | 448 | 3.79 | No |
| **Gin4-Full length** | **Hsl1** | ***elm1* $\Delta$ *gin4* $\Delta$** | **282** | **83** | **365** | **77.26** | **Yes** |
| **Gin4-Kinase dead** | **Hsl1** | ***elm1* $\Delta$ *gin4* $\Delta$** | **301** | **158** | **459** | **65.57** | **Yes** |
| **Hsl1** | **Shs1** | ***elm1* $\Delta$** | **264** | **462** | **726** | **36.36** | **Yes** |
| Hsl1 | Bud4 | *elm1* $\Delta$ | 110 | 607 | 717 | 15.34 | No |
| Hsl1 | Bni5 | *elm1* $\Delta$ | 4 | 354 | 358 | 1.11 | No |
| **Hsl1** | **Gin4** | ***elm1* $\Delta$** | **521** | **233** | **754** | **69.09** | **Yes** |
| **Hsl1** | **Kcc4** | ***elm1* $\Delta$** | **498** | **364** | **862** | **57.77** | **Yes** |
| Kcc4 | Shs1 | *elm1* $\Delta$ | 21 | 566 | 587 | 3.57 | No |
| Kcc4 | Bud4 | *elm1* $\Delta$ | 60 | 603 | 663 | 9.04 | No |
| **Kcc4** | **Hsl1** | ***elm1* $\Delta$** | **516** | **192** | **708** | **72.88** | **Yes** |
| Kcc4 | Bni5 | *elm1* $\Delta$ | 6 | 563 | 569 | 1.05 | No |
| Kcc4 | Gin4 | *elm1* $\Delta$ | 119 | 364 | 483 | 24.63 | No |
| **Elm1** | **Bud4** | ***gin4* $\Delta$** | **616** | **270** | **886** | **69.52** | **Yes** |
| **Elm1** | **Hsl1** | ***gin4* $\Delta$** | **399** | **289** | **688** | **57.99** | **Yes** |
| Elm1 | Hsl1-*ka1* $\Delta$ | *gin4* $\Delta$ | 139 | 470 | 609 | 22.82 | No |
| **Elm1** | **Shs1** | ***gin4* $\Delta$** | **499** | **301** | **800** | **62.375** | **Yes** |

To test whether Gin4 kinase activity is required for the observed rescue in cellular morphology upon artificial tethering of Gin4-GFP to the bud neck with Hsl1-GBP. We tethered Gin4^FL^-GFP and Gin4^KD^-GFP, expressed under the pTEF promoter, via Hsl1-GBP in *gin4*Δ *elm1*Δ cells, and observed that both constructs restored cellular morphology and aspect ratio to wild-type levels in 77.26% and 65.57% of cells, respectively **(Fig. 4F-4H**, **Table 1)**. These results suggest that Gin4 tethering to the bud neck via Hsl1 may compensate for Elm1 function in a kinase-independent manner, further reinforcing the kinase-independent nature of Gin4 function. Because Hsl1 also contains a membrane-binding C-terminal KA1 domain^51,57^, we tested its requirement for the observed functional rescue in *elm1*Δ cells and found that artificial tethering of Gin4-GFP via Hsl1-*ka1*Δ-GBP did not rescue the cellular morphology of *elm1*Δ cells **(Fig. 4I-4K)**. Together, these observations suggest that (i) restoring Gin4 localization to the bud neck via Hsl1 is necessary to partially rescue the morphological defects and cell shape exhibited by *elm1*Δ cells (ii) Gin4 might form a complex with Hsl1, which becomes essential in the absence of Elm1, indicating that Elm1 might play a significant role in regulating the Gin4-Hsl1 complex and (iii) proper membrane tethering of Hsl1 via its KA1 domain may be essential for its function in *elm1*Δ cells.

### Septin and cytokinesis defects are rescued by artificial tethering of Gin4-GFP to the bud neck via Hsl1-GBP in *elm1Δ* cells

Rescue of cell shape alone does not necessarily imply restoration of underlying physiological processes such as septin organization, AMR dynamics, or cytokinesis. To address this, we analyzed septin dynamics using Cdc3-mCherry in *elm1*Δ cells with and without artificial tethering of Gin4-GFP to the bud neck via Hsl1-GBP. Specifically, we examined the initial recruitment of septins during bud emergence. Consistent with previous results^38,39^ and our observations, 100% of *elm1*Δ cells exhibited mislocalization of Cdc3-mCherry to the bud cortex during bud emergence **(Fig. 5A, 5B)**. By contrast, only 35.71% of cells showed septin mislocalization when Gin4-GFP was artificially tethered to the bud neck via Hsl1-GBP **(Fig. 5A, 5B)**. We also analyzed the localization kinetics of Gin4-GFP in these strains and found that nearly 100% of *elm1*Δ cells exhibited Gin4-GFP mislocalization to the bud cortex in the absence of tethering, whereas this was reduced to 42.85% upon tethering via Hsl1-GBP **(Fig. 5A, 5C)**. The extent of rescue in Cdc3-mCherry mislocalization (64.28%) and Gin4-GFP mislocalization (57.14%) in *elm1*Δ cells closely correlated with the fraction of cells that also exhibited rescue of cellular morphology upon tethering with Hsl1-GBP **(Fig. 4B-4C, 5A-5C)**.

**Figure 5:**
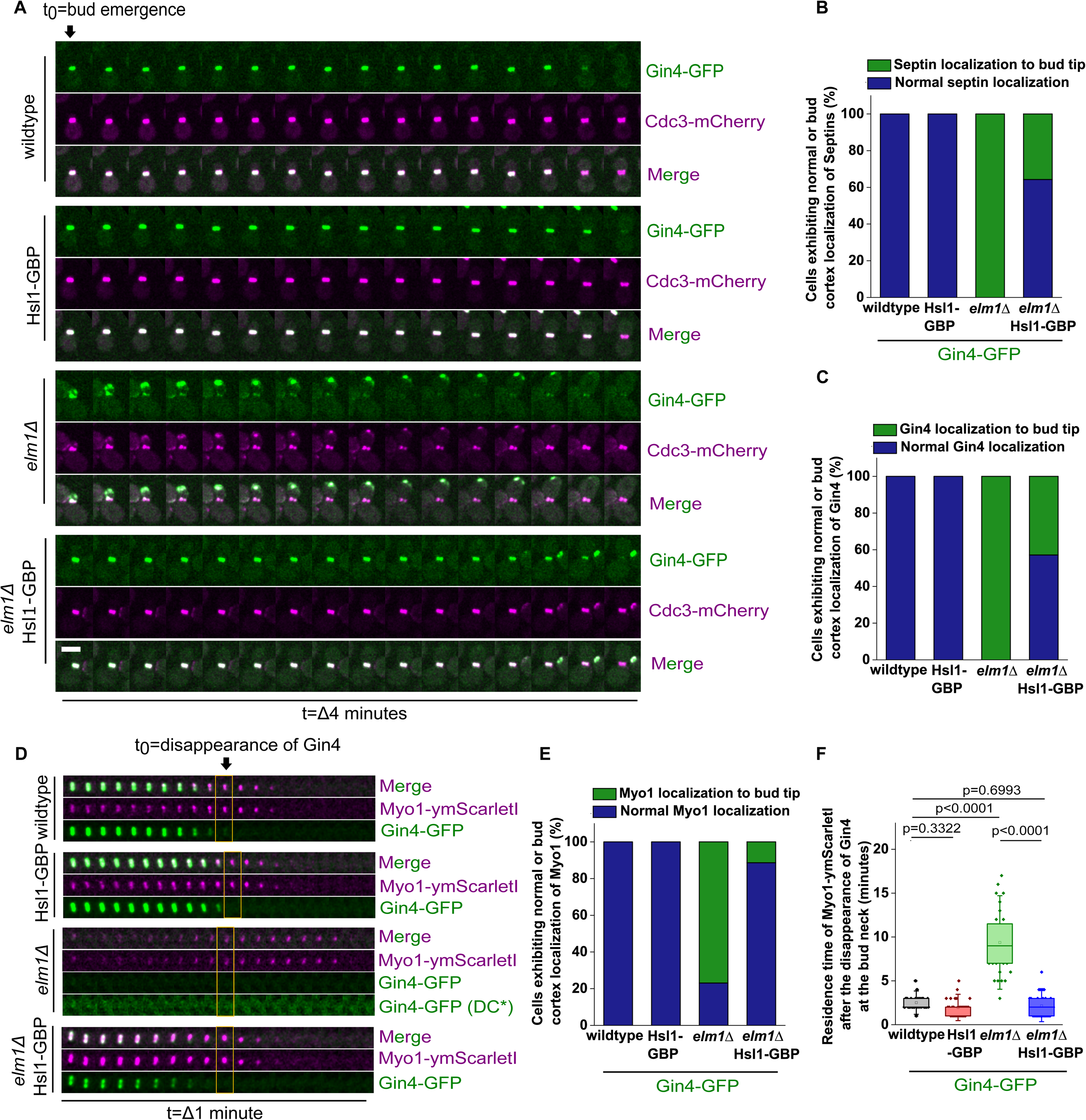
Redirecting Gin4 to the bud neck via Hsl1 in *elm1*Δ cells rescues septin organization and AMR dynamics. **(A)** Representative time-lapse montages showing the rescue of mislocalization of both Gin4-GFP and Cdc3-mCherry upon artificial tethering with Hsl1-GBP in *elm1*Δ cells. Scale bar-5µm. **(B)** Stacked column graph showing the percentage of cells exhibiting normal septin localization or septin mislocalization to bud cortex during the bud emergence in the indicated strains shown in (A) (wild-type: n=39, Hsl1-GBP: n=39, *elm1*Δ: n=39, *elm1*Δ-Hsl1-GBP: n=42 cells). **(C)** Quantification of Gin4-GFP localization rescue in the indicated strains shown in (A) (wild-type: n=39, Hsl1-GBP: n=39, *elm1*Δ: n=39, *elm1*Δ-Hsl1-GBP: n=42 cells). **(D)** Representative montages of the bud neck showing the constriction kinetics of Myo1-ymScarletI in *elm1*Δ strains in which Gin4-GFP is artificially tethered to bud neck via Hsl1-GBP**. (E)** Quantitative analysis for normal localization and mislocalization of Myo1-ymScarletI in the indicated strains shown in (D) (wild-type: n=41, Hsl1-GBP: n=37, *elm1*Δ: n=39, *elm1*Δ-Hsl1-GBP: n=44 cells). **(F)** Graph showing the constriction dynamics of Myo1-ymScarletI in the indicated strains shown in (D). Disappearance of Gin4 from the bud neck is used to normalize the initial time point for Myo1 constriction. Kruskal-Wallis nonparametric statistical test (****: p<0.0001, ns: p>0.05) (wild-type: n=39, Hsl1-GBP: n=30, *elm1*Δ: n=32, *elm1*Δ-Hsl1-GBP: n=39 cells).

To determine whether AMR defects were similarly rescued, we analyzed the dynamics of Myo1 C-terminally tagged with ymScarletI during cytokinesis. In *elm1*Δ cells, 76.92% of cells displayed Myo1 mislocalization to the bud cortex during its recruitment, consistent with previous reports^39^. Upon artificial tethering of Gin4-GFP to the bud neck via Hsl1-GBP, the proportion of cells exhibiting Myo1 mislocalization was reduced to 11.36% **(Fig. 5D, 5E)**. Additionally, we quantified the residence time of Myo1 at the bud neck by defining the disappearance of Gin4 as t_0_, corresponding to the onset of constriction, and measuring the time intervals until complete loss of the Myo1 signal from the bud neck. In wild-type cells, Myo1 constriction was completed within approximately 4 minutes, whereas in *elm1*Δ cells this process was prolonged to nearly 10 minutes **(Fig. 5D, 5F)**. The residence time was rescued to ∼4 minutes upon tethering Gin4-GFP to the bud neck with Hsl1-GBP in *elm1*Δ cells **(Fig. 5D, 5F)**.

Taken together, these results indicate that (i) rescue of cellular morphology correlates with restoration of both Gin4 and septin localization dynamics as the cell cycle progresses, suggesting a direct mode of regulation of Gin4 and Hsl1 downstream of Elm1 and (ii) restoration of AMR dynamics in *elm1*Δ cells upon artificial tethering of Gin4 to the bud neck via Hsl1 suggests an unconventional role for Hsl1 in the absence of Elm1.

### Rescue of septin localization in *elm1Δ* cells upon redirecting Gin4-GFP to the bud neck via Hsl1-GBP is largely independent of the morphogenetic checkpoint

Swe1 is a checkpoint kinase that regulates the morphogenetic signaling pathway and plays a key role in controlling cell shape by regulating isometric cell growth^41,59,61^. Previous studies have shown that Elm1, Gin4, Hsl1, and Kcc4 negatively regulate Swe1 through inhibitory phosphorylation, thereby promoting isometric growth and maintaining a round cell morphology^41,59^. It has also been reported that deletion of Swe1 in combination with septin kinase deletions (*elm1*Δ *swe1*Δ, *gin4*Δ *swe1*Δ, or *hsl1*Δ *swe1*Δ) rescues the elongated and clumped morphologies associated with the corresponding individual kinase deletions^41,61^. However, this rescue is specific to cell shape and does not restore septin localization or cytokinesis defects^41,61^. Based on these observations, we asked whether the rescue of cellular morphology and aspect ratio observed in *elm1*Δ cells upon artificial redirection of Gin4-GFP to the bud neck via Hsl1-GBP is mediated through negative regulation of Swe1 during mitotic spindle breakdown, or instead represents a direct consequence of restoring Gin4 and Hsl1 localization.

To address this, we deleted Swe1 in *elm1*Δ cells in which Gin4-GFP was either untethered or artificially tethered to the bud neck using Hsl1-GBP. Consistent with Bouquin et al., (2000)^61^, deletion of Swe1 alone rescued the elongated morphology in 60.66% of the *elm1*Δ population when Gin4-GFP was not tethered to the bud neck **(Fig. S6A-S6C)**. Strikingly, deletion of Swe1 did not alter the proportion of round cells in *elm1*Δ cells in which Gin4-GFP was redirected to the bud neck via Hsl1-GBP **(Fig. S6A-S6C)**.

We further analyzed septin organization using Cdc3-mCherry in *elm1*Δ and *elm1*Δ *swe1*Δ cells, with and without artificial tethering of Gin4-GFP to the bud neck via Hsl1-GBP. Although cell shape was rescued in both *elm1*Δ *swe1*Δ control cells and *elm1*Δ *swe1*Δ cells with Gin4-GFP tethered via Hsl1-GBP, only 23.63% of cells in the *elm1*Δ *swe1*Δ control strain exhibited rescue of septin mislocalization during bud emergence **(Fig. 6A, 6B)**. This partial rescue by Swe1 deletion also correlated with restoration of mislocalized Gin4 during bud emergence in *elm1*Δ cells **(Fig. 6C, 6D)**. Notably, more than 70% of cells exhibited rescue of both Gin4 and Cdc3 mislocalization in *elm1*Δ *swe1*Δ cells when Gin4-GFP was artificially tethered to the bud neck via Hsl1-GBP, in contrast to the rescue exhibited by only ∼23% of cells in which Gin4-GFP was not tethered **(Fig. 6A-6D)**.

**Figure 6:**
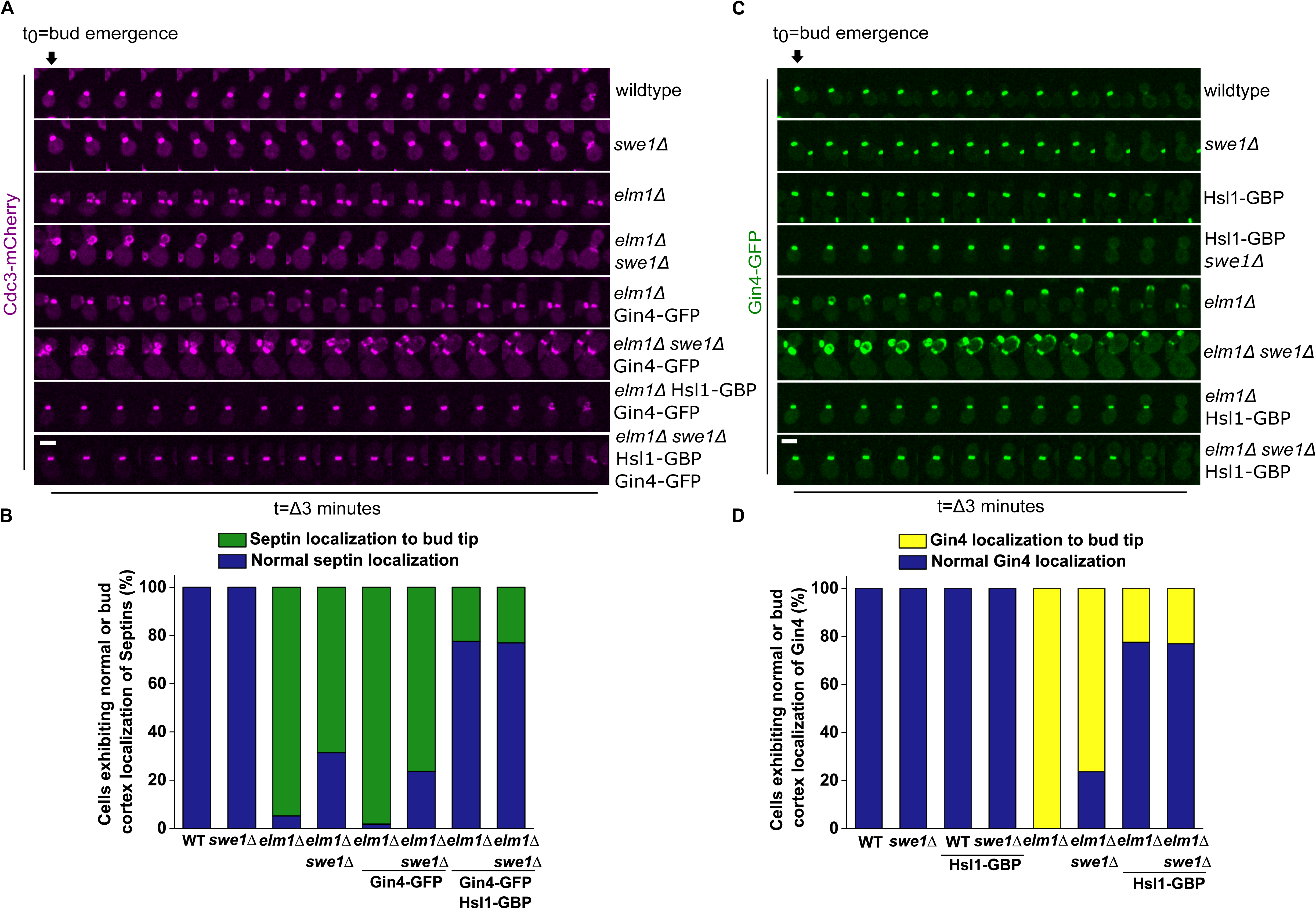
Gin4 tethering to the bud neck via Hsl1 bypasses the requirement for Elm1 independently of the morphogenetic checkpoint kinase Swe1. **(A)** Time-lapse montages representing the dynamics of Cdc3-mCherry in both *elm1*Δ and *elm1*Δ *swe1*Δ strains upon artificial tethering of Gin4-GFP to bud neck with Hsl1-GBP (t_0_ = bud emergence). Scale bar-5µm. **(B)** Stacked column graph showing the percentage of cells exhibiting normal localization and mislocalization of Cdc3-mCherry in the indicated strains shown in (A) (wild-type: n=55, *swe1*Δ: n=49, *elm1*Δ: n=58, *elm1*Δ *swe1*Δ: n=51, *elm1*Δ-Gin4-GFP: n=55, *elm1*Δ *swe1*Δ-Gin4-GFP: n=55, *elm1*Δ-Gin4-GFP-Hsl1-GBP: n=58, *elm1*Δ *swe1*Δ-Gin4-GFP-Hsl1-GBP: n=52 cells). **(C)** Representative time-series montages showing the localization of Gin4-GFP in both *elm1*Δ and *elm1*Δ *swe1*Δ strains upon artificial tethering of Gin4-GFP to the bud neck via Hsl1-GBP (t_0_ = bud emergence). Scale bar-5µm. **(D)** Quantification graphs depicting the normal localization and mislocalization of Gin4-GFP at the bud cortex during bud emergence in the indicated strains shown in (C) (wild-type: n=57, *swe1*Δ: n=52, Hsl1-GBP: n=54, Hsl1-GBP *swe1*Δ: n=51, *elm1*Δ-Gin4-GFP: n=55, *elm1*Δ *swe1*Δ-Gin4-GFP: n=55, *elm1*Δ-Gin4-GFP-Hsl1-GBP: n=58, *elm1*Δ *swe1*Δ-Gin4-GFP-Hsl1-GBP: n=52 cells).

Collectively, these findings suggest that (i) the rescue of elongated cell morphology to normal round cell morphology in *elm1*Δ cells upon artificial tethering of Gin4-GFP to the bud neck via Hsl1-GBP may depend on Hsl1-mediated regulation of the morphogenesis checkpoint; however, (ii) the observed rescue in septin and Gin4 localization clearly does not depend on the morphogenesis checkpoint and reflects direct regulation of septin organization by Gin4 and Hsl1.

Intrigued by the rescue in cellular morphology observed only after tethering Gin4-GFP to Hsl1-GBP and not to other bud neck proteins, we extended our assay to its related kinases and tethered Hsl1-GFP and Kcc4-GFP via various bud-neck proteins in *elm1*Δ cells **(Fig. 7A-7C and S7A-S7C)**. Our results revealed that Hsl1-GFP restored normal cell morphology in 36.36% of cells in *elm1*Δ when tethered with Shs1-GBP, while tethering with Gin4-GBP or Kcc4-GBP resulted in significantly higher rescue of 69.09% and 57.77%, respectively, suggesting possible coordination among these kinases **(Fig. 7A-7C**, **Table 1)**. Strikingly, like Gin4-GFP, Kcc4-GFP tethering also restored cell morphology similar to wild-type in 72.88% of the population, only when tethered to the bud neck with Hsl1-GBP **(Fig. S7A-S7C, Table 1)**. However, Kcc4-GFP tethering via different bud neck proteins did not rescue the phenotype, although a minor rescue in 24.63% of cells was observed upon tethering with Gin4-GBP, suggesting possible crosstalk between Kcc4 and Gin4 **(Fig. S7A-S7C, Table 1)**.

**Figure 7:**
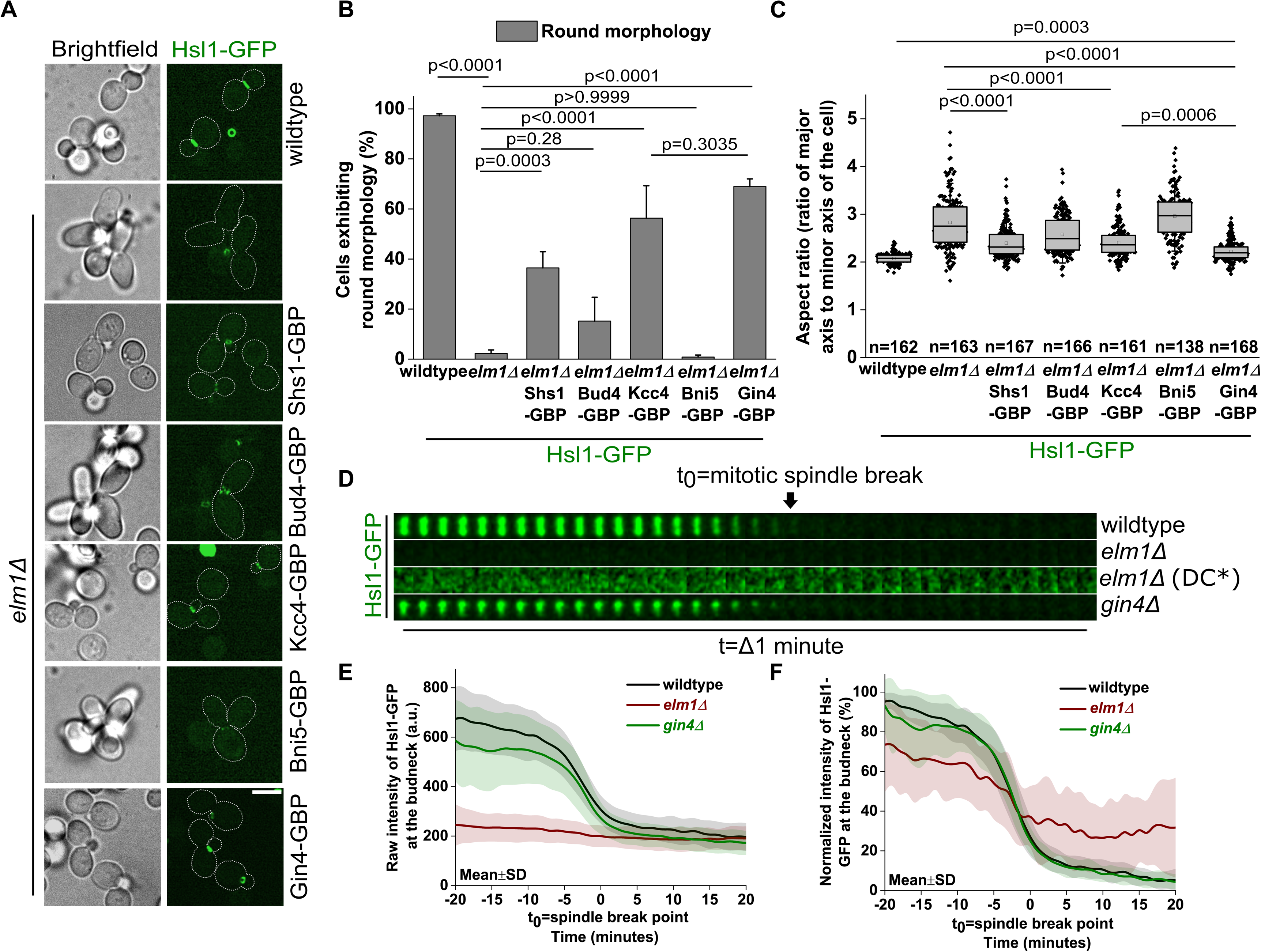
Restoring Hsl1-GFP localization to the bud neck via septins or its related kinases can bypass the requirement of Elm1. **(A)** Representative images showing artificial tethering of Hsl1-GFP to the bud neck via Shs1-GBP, Bud4-GBP, Kcc4-GBP, Bni5-GBP and Gin4-GBP in *elm1*Δ cells. Scale bar-5µm. **(B)** Bar graph showing the percentage of cells exhibiting round morphology in the indicated strains shown in (A), one-way ANOVA with Tukey’s multiple comparison test (****: p<0.0001, ns: p>0.05) (N=3, wild-type: n=724, *elm1*Δ: n=458, *elm1*Δ-Shs1-GBP: n=726, *elm1*Δ-Bud4-GBP: n=717, *elm1*Δ-Kcc4-GBP: n=862, *elm1*Δ-Bni5-GBP: n=358 and *elm1*Δ-Gin4-GBP: n=754 cells). **(C)** Quantification of aspect ratios in the indicated strains shown in (A), Kruskal-Wallis nonparametric statistical test (***: p<0.001, ****: p<0.0001) (N=3, n>135 cells/strain). **(D)** Representative montages of the bud neck depicting the localization of Hsl1-GFP in *elm1*Δ and *gin4*Δ cells during cytokinesis. DC*=Differential contrast. **(E)** Plot of raw fluorescence intensity of Hsl1-GFP in the indicated strains shown in (D) (t_0_=spindle breakpoint) (wild-type: n=29, *elm1*Δ: n=24, *gin4*Δ: n=38 cells). **(F)** Plot of normalized fluorescence intensity of Hsl1-GFP in the indicated strains shown in (D) (t_0_=spindle breakpoint).

To support these results, we analyzed the localization of Hsl1 and Kcc4 in *elm1*Δ and *gin4*Δ cells. As reported previously^39^, Kcc4 exhibited mislocalization to the bud cortex during the initial phases of the cell cycle and defective localization at the onset of cytokinesis **(Fig. S8B-S8E)**. Recruitment and localization of Hsl1 to the bud neck were also severely impaired in *elm1*Δ cells **(Fig. 7D-7F, S8A)**, suggesting a role for Elm1 in maintaining Hsl1 localization at the bud neck. By contrast, recruitment and localization of Hsl1 and Kcc4 were similar to wild-type in *gin4*Δ cells **(Fig. 7D-7F, S8A-S8E)**. These data suggest that loss of Hsl1 at the bud neck may contribute significantly to the elongated phenotype observed in *elm1*Δ cells, and, consistent with this idea, restoration of Hsl1 localization rescues the observed morphological defects in our experiments. The degree of phenotypic rescue appears to depend on the presence of Hsl1 at the bud neck, and higher rescue percentages are observed when Hsl1 is tethered with another kinase (Gin4 or Kcc4), suggesting coordinated effects among these kinases and that Hsl1 may act as a downstream effector in the septin kinase network.

Our results reveal that Hsl1 regulates normal cell morphology, possibly through downstream signaling events triggered by the presence of Elm1 in wild-type cells, in coordination with either Gin4 or Kcc4. Thus, Elm1 may act upstream to control the localization and activation of the Nim1-related kinases (Hsl1, Gin4, and Kcc4) at the septin scaffold in a precise, spatiotemporal manner, within a network with multiple feedback regulatory links. Overall, the ability to restore normal function by simply rewiring the septin kinase network demonstrates the inherent redundancy and robustness of this network in *S. cerevisiae*.

### Hsl1 and its membrane-binding ability are required for the rescue of *gin4*Δ cells by artificially tethered Elm1

Elm1 does not localize to the bud neck in the absence of Gin4 and can rescue the elongated/clumped phenotype exhibited by *gin4*Δ cells upon artificial tethering to the bud neck septin scaffold^38^. We corroborated these findings independently and found that Elm1 is not recruited to the bud neck in *gin4*Δ cells **(Fig. S9A-S9D)**. We found that Elm1-GFP tethering via Shs1-GBP, Bud4-GBP, or Hsl1-GBP could restore normal cell morphology and aspect ratio in *gin4*Δ cells **(Fig. S9E-S9G)**. We therefore asked whether the presence of Hsl1 is necessary for the observed rescue upon Elm1 tethering to the bud neck in *gin4*Δ cells. To test this, we artificially tethered Elm1-GFP to the bud neck via Shs1-GBP in *gin4*Δ and *gin4*Δ *hsl1*Δ cells. While 62.37% of *gin4*Δ cells showed restoration of cell morphology and aspect ratio, no rescue was observed in the *gin4*Δ *hsl1*Δ double deletion strain upon tethering Elm1 to the bud neck, indicating that Hsl1 is required downstream of Elm1 tethering in *gin4*Δ cells **(Fig. 8A, 8C, 8D**, **Table 1)**. Additionally, we observed rescue of septin misorganization upon tethering Elm1-GFP to the bud neck with Shs1-GBP in *gin4*Δ cells in 82.92% of the cellular population, which was lost in *gin4*Δ *hsl1*Δ cells in >75% of the population **(Fig. 8A, 8E)**. Redirected Elm1-GFP also showed mislocalization to the bud cortex during bud emergence in *gin4*Δ *hsl1*Δ cells **(Fig. 8F)**. A loss of rescue was observed when Elm1-GFP was tethered to the bud neck in *gin4*Δ cells via Hsl1-*ka1*Δ-GBP, suggesting that Hsl1 acts downstream of Elm1 and that its membrane-binding KA1 domain is required for its function **(Fig. S9H-S9J)**. These data support the idea that Hsl1 acts downstream of Elm1 and Gin4 in the septin kinase network to control septin stability.

**Figure 8:**
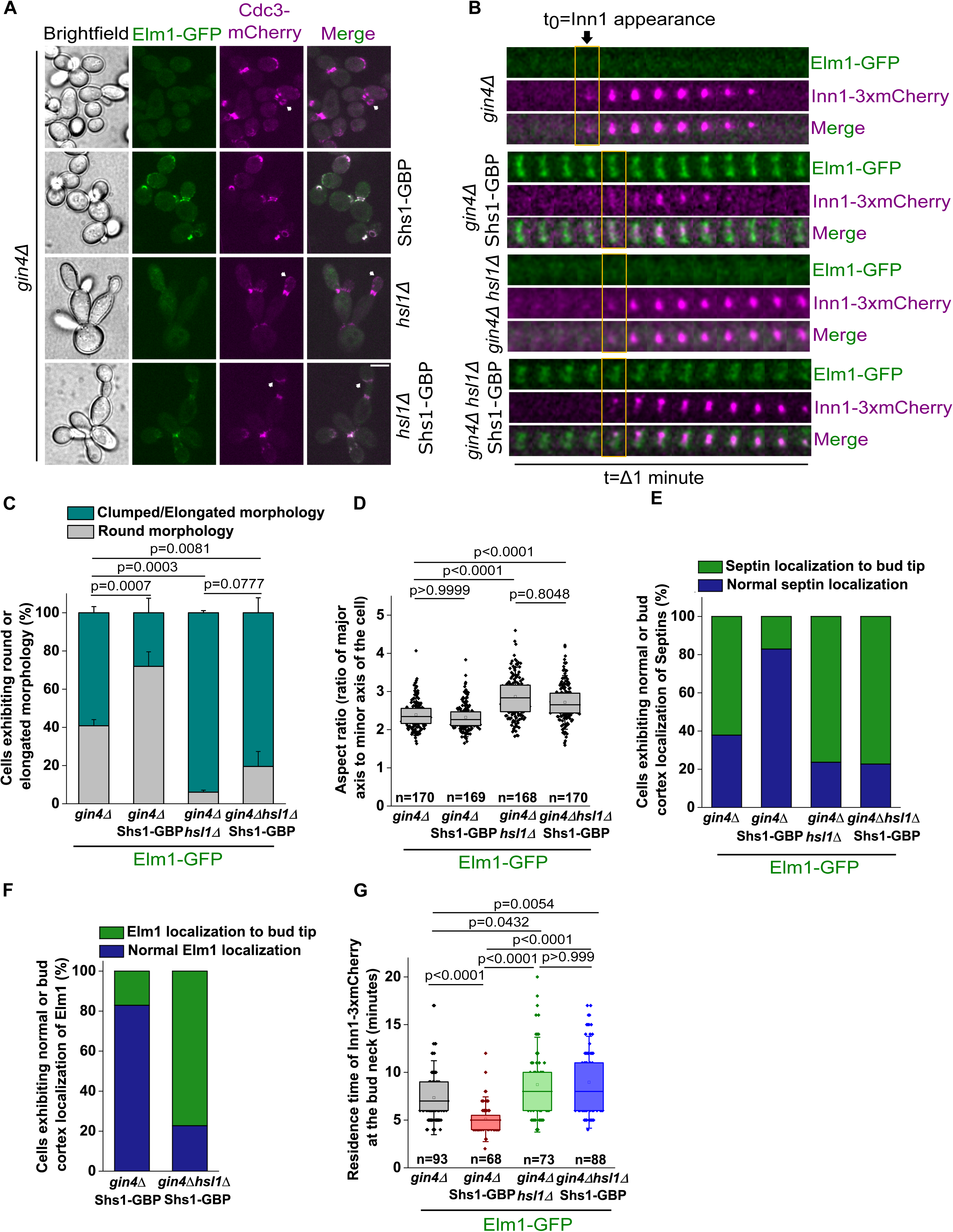
A non-canonical role for Hsl1-Kinase in regulating septin organization and AMR dynamics. **(A)** Representative images showing Cdc3-mCherry mislocalization during bud emergence in *gin4*Δ and *gin4*Δ *hsl1*Δ cells in which Elm1-GFP is artificially tethered to bud neck via Shs1-GBP. White arrows indicate mislocalized Cdc3-mCherry in the represented strains. Scale bar-5µm. **(B)** Representative time-lapse montages of the bud neck showing the constriction profile of Inn1-3xmCherry in *gin4*Δ and *gin4*Δ *hsl1*Δ cells in which Elm1-GFP is artificially tethered to bud neck via Shs1-GBP. **(C)** Bar graph showing the percentage of cells exhibiting round and elongated morphologies in the indicated strains shown in (A), one-way ANOVA with Tukey’s multiple-comparison test (**: p<0.01, ***: p<0.001, ns: p>0.05). p-values correspond to population of cells exhibiting round morphology, (N=3, *gin4*Δ: n=689, *gin4*Δ-Shs1-GBP: n=639, *gin4*Δ *hsl1*Δ: n=492, *gin4*Δ *hsl1*Δ-Shs1-GBP: n=590 cells). **(D)** Quantification of aspect ratios in the indicated strains shown in (A), Kruskal-Wallis nonparametric statistical test (****: p<0.0001, ns: p>0.05), (N=3, n>165 cells/strain). **(E)** Stacked column graph representing the percentage of cells exhibiting normal septin localization and septin mislocalization in the indicated strains of (A) (*gin4*Δ: n=44, *gin4*Δ-Shs1-GBP: n=41, *gin4*Δ *hsl1*Δ: n=38, *gin4*Δ *hsl1*Δ-Shs1-GBP: n=44 cells). **(F)** Quantification of normal localization and mislocalization of Elm1-GFP in the indicated strains shown in (A) (*gin4*Δ-Shs1-GBP: n=37, *gin4*Δ *hsl1*Δ-Shs1-GBP: n=44 cells) **(G)** Quantification of the residence time of Inn1-3xmcherry during cytokinesis in the indicated strains shown in (B), Kruskal-Wallis nonparametric statistical test (*: p<0.05, **: p<0.01, ****: p<0.0001, ns: p>0.05) (N=2, n>60 cells/strain).

Next, we asked whether Elm1-GFP tethering to the bud neck in *gin4*Δ cells could rescue other observed cellular defects related to AMR dynamics during cytokinesis. To test this, we first assessed the recruitment and localization of the Myo1 during bud emergence. We observed that Myo1-3xmCherry exhibited mislocalization to the bud tip in 12.82% *gin4*Δ cells during bud emergence **(Fig. S10A, S10B)**. However, Myo1-3xmCherry localization was restored to the bud neck upon artificial tethering of Elm1-GFP in *gin4*Δ cells. The cellular population with mislocalized Myo1-3xmCherry increased further upon deletion of Hsl1 in *gin4*Δ cells, even when Elm1-GFP was artificially tethered to the bud neck with Shs1-GBP, reaching up to 87.17% of the cellular population **(Fig. S10A, S10B)**. We also assessed the residence time of Inn1-3xmCherry in these cells to measure constriction dynamics of the AMR^75,76^. We found that the Inn1-3xmCherry residence time was similar to that of wild-type cells when Elm1-GFP was tethered via Shs1-GBP in *gin4*Δ cells, but was significantly increased when Elm1-GFP was tethered via Shs1-GBP in *gin4*Δ *hsl1*Δ cells **(Fig. 8B, 8G)**. These results suggest that (i) Elm1 tethering to the bud neck restores normal Myo1 localization and AMR constriction dynamics in the absence of Gin4, (ii) Hsl1 is required for Elm1 to exert its influence over AMR organization and dynamics after bud neck tethering in *gin4*Δ cells, and (iii) the membrane-binding KA1 domain of Hsl1 is required for this activity downstream of Gin4.

Taken together, our results suggest that Hsl1 functions both downstream of and in coordination with Elm1 and Gin4 to maintain septin stability and ensure proper cytokinesis **(Fig. 9)**. In wild-type cells, Hsl1 likely acts downstream of Elm1 and Gin4 and localizes normally to the bud neck, where it may help coordinate septin architecture with AMR constriction during cytokinesis **(Fig. 9)**. In the absence of Elm1, localization of both Gin4 and Hsl1 becomes defective during later cell cycle stages, disrupting the potential crosstalk between Gin4 and Hsl1. This disruption may lead to septin mislocalization and disorganization of the AMR machinery **(Fig. 9)**. Notably, restoring Hsl1 localization using a GFP-GBP tethering strategy strengthens this link, suggesting that Elm1 plays an important role in recruiting or stabilizing Hsl1 at the bud neck. These observations indicate that Hsl1 acts downstream of Elm1 in this pathway, with partially overlapping functions. Furthermore, the requirement for Hsl1 in the rescue observed upon Elm1 tethering in *gin4*Δ cells supports the idea that Hsl1 mediates downstream effects of Elm1 in regulating septin stability and cytokinesis. Therefore, we propose that Hsl1, through a compensatory pathway, facilitates functional crosstalk between the septin kinase network and septin/AMR components, which is essential for proper septin organization and symmetric AMR constriction, ultimately ensuring faithful cytokinesis.

**Figure 9:**
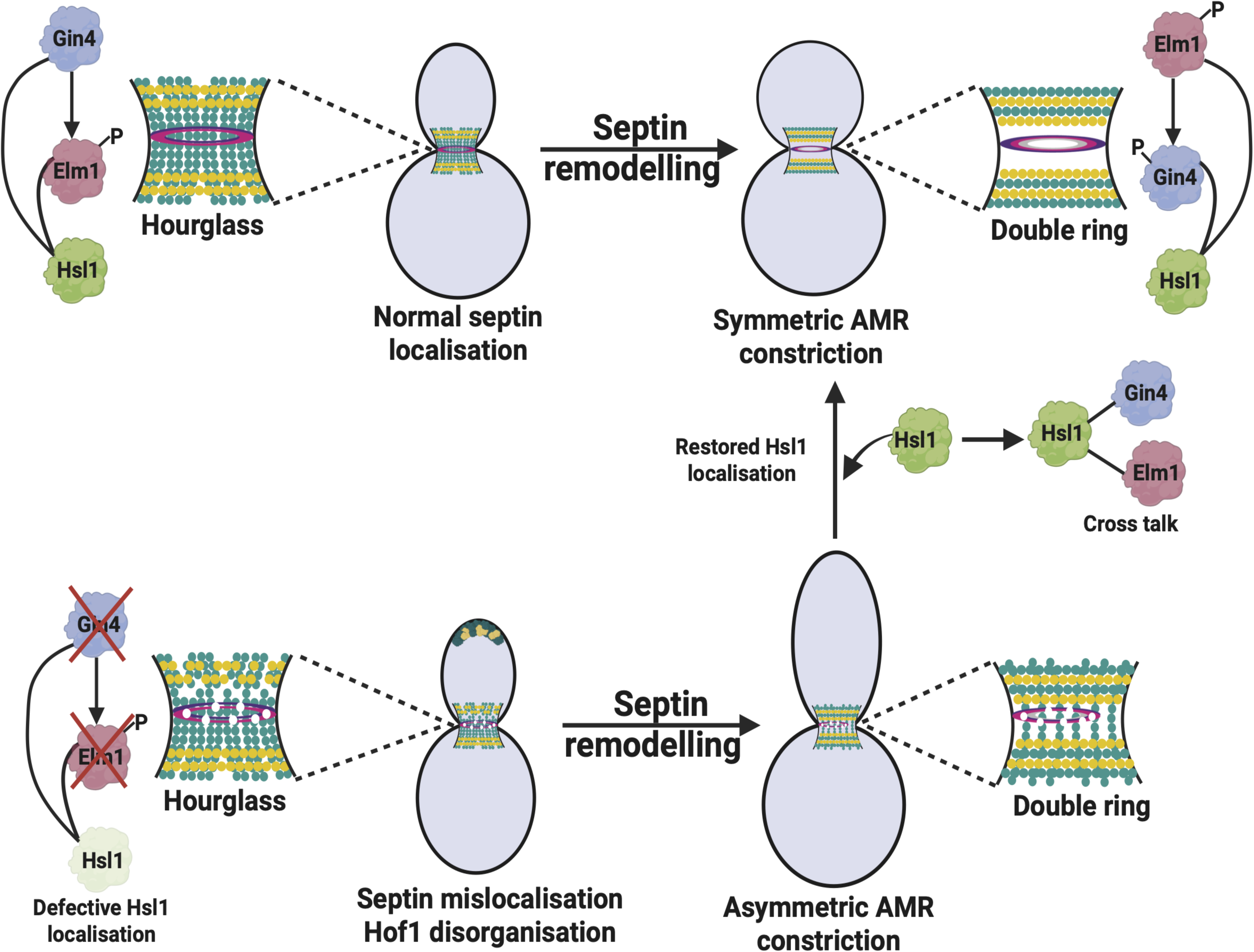
Representative model for the role of Hsl1 kinase in septin organization and AMR constriction downstream of Gin4 and Elm1.

## DISCUSSION

How septin architecture remodeling crosstalk with the AMR machinery and cytokinesis remains a dynamic area of investigation, as these processes are spatiotemporally controlled by septin-interacting proteins and PTMs^37,46,63–68^. Yet the regulatory mechanisms governing septin remodeling and AMR constriction during the cell cycle in yeast and higher eukaryotes remain incompletely understood. In *S. cerevisiae*, four septin-associated kinases Elm1, Gin4, Hsl1, and Kcc4 have been reported to primarily regulate septin-dependent processes^38,39,59,61^. However, the precise mechanisms by which these kinases influence septin architecture and coordinate cytokinesis remain poorly understood. In this study, we identify critical roles for Elm1 and Gin4 in maintaining proper septin organization and AMR dynamics during cytokinesis. Our results further reveal that Gin4 exerts its functions in septin stability and AMR dynamics in a kinase-independent manner, challenging the prior view of kinase-dependent functions. We also identify a direct physical interaction between Gin4 and the AMR protein Hof1, providing new insights into how septin regulatory pathways directly crosstalk with the cytokinetic machinery.

Septin remodeling marks the onset of AMR constriction and cytokinesis^17,18,30,82^, but the molecular mechanisms coordinating these processes remain largely unclear. We observed that septins are severely mislocalized to the bud cortex in *elm1*Δ and *gin4*Δ cells, supporting previous reports that these kinases are required for proper septin organization at the bud neck^38,39,59^. Consistent with these defects, the dynamics of the AMR complex proteins Inn1 and Hof1 were significantly perturbed, leading to delayed constriction in the absence of Elm1 and Gin4. These observations suggest that septin-associated kinases regulate intermediate signaling events that coordinate septin architecture with the assembly and stability of the cytokinetic machinery, and they highlight an underappreciated mode of regulation of AMR organization and dynamics **(Fig. 1D-1F, 2A-2E)**.

A key question arising from these observations is whether the crosstalk between the septin kinase network and AMR dynamics is direct and septin-independent, or indirect and septin-dependent. Previous studies have shown that AMR constriction can proceed in certain septin mutants that disrupt septin organization, suggesting that septin kinases may also exert direct regulatory effects on the cytokinetic machinery^17^, further supporting a direct contribution of Elm1 and Gin4 to the dysregulated cytokinetic events observed in this study. Supporting this idea, we found that Gin4 interacts with the F-BAR protein Hof1, as demonstrated by yeast two-hybrid and in-vitro binding assays **(Fig. 2F-2H)**. Notably, the membrane-binding domains of Gin4 and Hof1 were sufficient for this interaction, suggesting that Gin4 activation at the membrane^48,57^ may directly influence Hof1 organization and function during AMR maturation **(Fig. 2H)**. These findings suggest a potential mechanism for anchoring Hof1 to the plasma membrane prior to cytokinesis **(Fig. 2I)**. Interestingly, previous work has shown that *gin4*Δ and *hof1*Δ are synthetic lethal and that Hof1 exhibits mislocalization in *gin4*Δ cells^74^. These results also point to a coordinative mechanism in which Gin4 directly crosstalks with an AMR component, Hof1, to regulate AMR organization and constriction. To our knowledge, this represents the first evidence physically linking the septin kinase network to AMR proteins and highlights a novel regulatory input on AMR assembly in *S. cerevisiae*. Future work is required to delineate the molecular mechanisms underlying this regulation.

Interestingly, our results further demonstrate that the kinase activity of Gin4 is not required for its role in septin organization, AMR constriction **(Fig. 3A-3E)**, or cell viability in the absence of Hof1 **(Fig. S3I)**. This highlights an important aspect of Gin4 function in cells and is similar to the kinase-independent function reported for Elm1^39^. These observations raise an important question regarding the functional significance of kinase activity within the septin kinase network. Our results, together with previous studies^38,59^, establish that built-in functional redundancies exist among the septin kinases (Elm1, Gin4, Hsl1 and Kcc4), and that their recruitment to the bud neck is also interdependent. Kinase activities may play an important role in regulating the signaling that coordinates the precise arrival and spatiotemporal activation of these kinases at the bud neck. These results also raise important questions about post-translational regulatory mechanisms of septin filament organization, as a number of studies have shown that these kinases directly phosphorylate septins^49^. Thus, the role of post-translational regulation of septins will be better understood with future work in this direction.

Although these kinases exhibit distinct functions, they also display mutual interdependence^38,62^. Recent studies have proposed a new perspective regarding this crosstalk, revealing reciprocal regulation between Gin4 and Elm1 through direct binding and phosphorylation at different stages of the cell cycle^38^. The existence of redundancy in the septin kinase network explains the longstanding differences in the phenotypes observed in *elm1*Δ and *gin4*Δ cells. The *elm1*Δ cells show more pronounced abnormalities in cell shape and septin organization than *gin4*Δ cells, even though Elm1 is known to be absent from the bud neck in *gin4*Δ cells^38,41,59^. Our analysis of the temporal localization dynamics of Hsl1 and Kcc4 suggests a potential explanation for this discrepancy. Specifically, the recruitment of Hsl1 and Kcc4 to the bud neck was strongly perturbed in *elm1*Δ cells but remained largely intact in *gin4*Δ cells **(Fig. 7D-7F, S8A-S8E)**. These findings suggest that the localization of Hsl1 and Kcc4 to the bud neck is crucial for their normal function and may explain the less prominent phenotype observed following loss of Gin4, emphasizing the redundant roles of Hsl1 and Kcc4 in septin organization.

Consistent with this interpretation, our GFP-GBP tethering experiments demonstrated that restoring localization of Hsl1 in *elm1*Δ cells, together with Gin4 and Kcc4, can bypass the requirement for Elm1 during the cell cycle, independent of its function in the morphogenetic checkpoint pathway **(Fig. 4B-4E, 6A-6D, 7A-7C and S7A-S7C)**. These findings highlight a previously underappreciated level of functional redundancy among the Nim1-related kinases. Our results also suggest that Hsl1 can play a non-canonical role in regulating septin organization and AMR dynamics downstream of Elm1 and Gin4 **(Fig. 5A-5F, 8A-8G, S10A, S10B)**. We therefore speculate that synergy exists among Elm1, Gin4, and Hsl1 to regulate downstream signaling pathways that control septin organization and cytokinesis. Investigating the requirement for Hsl1 kinase activity in the absence of Elm1 or Gin4 may provide further insight into the molecular mechanisms underlying Hsl1 regulation. Additionally, elucidating the phosphorylation-dependent modifications involved, along with identifying the specific regulatory residues in future studies, will help clarify the molecular mechanisms by which these kinases coordinate with one another to ensure the precise execution of cytokinesis.

Strikingly, deletion of the membrane-binding KA1 domain of Hsl1 abolished its ability to rescue the elongated defect exhibited in the absence of Elm1 **(Fig. 4I-4K and S9H-S9J)**, emphasizing the importance of membrane association for the proper function of these kinases^57^. Membrane binding may facilitate kinase activation or enable interactions with septin filaments and other regulatory proteins at the bud neck. However, the precise mechanisms by which membrane association contributes to Hsl1 function remain unclear and will require further investigation. The underlying molecular mechanisms and actual cause-and-effect relationships by which Hsl1 and its related kinases regulate septin organization at the filament-level remain to be understood. Approaches such as platinum-replica electron microscopy (PREM) and in-vitro reconstitution assays may prove instrumental in dissecting how these kinases influence septin filament organization and higher-order assembly in future studies^27,39,82^. Such studies will provide critical insights into the modes of regulation of septins through the coordinated action and signaling inputs from multi-septin kinase modules across eukaryotes.

In summary, our study provides a comprehensive quantitative analysis of septin-regulatory kinases in controlling septin organization and cytokinesis. We identify Elm1 and Gin4 as key regulators of septin architecture and uncover previously uncharacterized roles for these kinases in AMR organization and constriction. These findings pave the way for future investigations into the intricate molecular mechanisms and cross-regulatory interactions within the septin-kinase network and with AMR proteins during the cell cycle in *S. cerevisiae* and other related organisms, where septins and their associated proteins play key roles in processes such as cell division.

## Supporting information

Bhojappa et al_Supplementary figures_legend

## Acknowledgements

We thank the DST-FIST microscope facility of the Department of Biochemistry, Indian Institute of Science. We also thank the Bioimaging facility, Division of Biological Sciences. We thank Prof. Erfei Bi (University of Pennsylvania) and Prof. Gislene Periera (Centre for Organismal Studies (COS) Heidelberg) for the plasmids. We are grateful for the help of Ms. Spoorthi Hirematt in image acquisition at the DST-FIST microscope facility. We thank Prof. P. N. Rangarajan, Prof. Franz Meitinger, and Prof. Ramanujam Srinivasan for their valuable suggestions on the manuscript. Schematic diagrams were created in BioRender. B.B. acknowledges the DST-INSPIRE Fellowship. A.D. and J.K. acknowledge the IISc Ph.D. fellowship from IISc. F.C. acknowledges the ICMR fellowship.

## Author contributions

Conceptualization: S.P.; Funding acquisition: S.P.; Supervision: S.P.; Project administration: S.P.; Methodology: S.P., B.B.; Investigation: B.B., A.D., B.V.T., J.K., F.C., D.G.; Formal analysis: B.B., A.D., V.R.; Resources: S.P., B.B., B.V.T., J.K.; Data curation: B.B.; Visualization: B.B.; J.K.; F.C. B.V.T; Writing- original draft: S.P., A.D., B.B.; Writing-review and editing: All authors

## Materials and Methods

### Strain construction

All yeast strains in this study were constructed using the *S. cerevisiae* wild-type strains ESM356 and YPH499, derived from the S288C genetic background. The strain number and the genotype are listed in supplementary table S1. C-terminal epitope tagging and endogenous gene deletions were carried out based on the PCR-based integration strategy^95,96^. All strains were cultured at 23°C with shaking at 180 rpm. Synthetic complete (S.C.) media lacking tryptophan, uracil, leucine, and histidine were used to select auxotrophic recombinants. Antibiotic selections for endogenous gene manipulation were done on YPD agar plates supplemented with 200 µg/ml G418 (Sisco Research Laboratories Pvt. Ltd.; catalogue no. 58327), 100 µg/ml hygromycin-B (Sisco Research Laboratories Pvt. Ltd.; catalogue no. 67317), and 100 µg/ml nourseothricin (Jena Biosciences; catalogue no. AB-102). To visualize septin dynamics, Cdc3-mCherry, YIP211-cdc3-mCherry^72,97^, and YIP204-cdc3-mCherry^77^ plasmids were cut with BglII (catalogue no. R0144S; New England Biolabs) and integrated into the CDC3 locus. Cell cycle dynamics were visualized using a yeast expression vector with mRuby2-Tub1 under the His3 promoter (pHIS3p:mRuby2-Tub1+3"UTR::Ura3)^98^ and pAFS125-GFP-TUB1^99^, which were cut with ApaI (Catalogue no. R0114S; New England Biolabs) and StuI (catalogue no. R0187S; New England Biolabs), respectively, and integrated into the ura3 loci. Yeast transformation for epitope tagging and endogenous gene deletions was performed using a lithium acetate (LiOAc)-based protocol as described^100^.

### Oligonucleotides and Plasmids

All oligonucleotides used for endogenous gene tagging and deletion in this study are listed in supplementary table S2. The plasmids used and generated for this study are listed in supplementary table S3. For the generation of clones used in **Fig. 3A** and **3B,** Gin4^FL^ was amplified from the endogenous locus and integrated into pRS305 under the pTEF promoter. The ligation reaction was performed using NEB Hifi-Builder (New England Biolabs; cat. no.: E2621L). Catalytically dead Gin4^KD^(K48A) was constructed based on the site-directed mutagenesis strategy using the oligos (P855, P856, P857, and P1024). The kinase-dead mutant was confirmed correct by sequencing. To construct Gin4^FL^ and Gin4^KD^ used in **Fig. 3C-3E, S3F, S3G, and S3I**, the Gin4 gene fragment was amplified from the genomic loci with the endogenous promoter using PCR and was ligated into a pRS305 integration vector cut with BamHI-HF (Catalogue no. R3136S; New England Biolabs).

### Live-cell imaging

Cells were inoculated in S.C. media and incubated overnight at 23°C. The secondary culture was diluted to log phase (optical density-0.4 to 0.6) from the overnight saturated culture. 6% Concanavalin A type 6 (Sigma-Aldrich; catalogue no. C2010) was coated onto a confocal dish (ibidi GmbH-catalogue no. 81218-200; Cellvis-catalogue no. D35C4-20-1.5-N) and was incubated at 23°C. The secondary culture cells were pelleted at 3200 rpm for 3 minutes. The cells were washed in S.C. media to remove non-adherent cells and subjected to live-cell imaging.

The laser point scanning confocal microscope Olympus FV-3000, equipped with high-sensitivity GaAsP photomultiplier tube (PMT) detectors and solid-state lasers (488 nm and 561 nm), was used to acquire time-lapse imaging for **Fig. S1A-S1D, S2A**. Images were acquired at a one-minute time interval with 0.5 µm step size and seven z-slices for 2-3 hours. The raw images were three-dimensionally (3D) deconvolved in Olympus CellSens Dimension (3.1) software. Maximum-intensity projection images were used for representation.

The Olympus IX83 widefield microscope equipped with a CoolLED PE-4000 LED light source and Prime-BSI ScMOS camera was used for acquiring images shown in **Fig. 1B, 1D, 2A, 3C, and S3C**. All images were acquired with a 100x oil-immersion objective with 1.45 N.A., 1 µm step-size, and three z-slices at a 2-minute time interval for 60-150 minutes. The raw images were processed for 3D deconvolution using Olympus CellSens Dimension (3.1) software. Maximum-intensity projection images were used for representation.

The images for Hof1 mis-organization and artificial tethering experiments in **Fig. 2D, S3A, S3F, 4B, 4I, 7A, S5E, S7A, S9E, and S9H** were acquired using the Oxford Andor Dragonfly 502 spinning disc confocal system equipped with a fully motorized Dmi8 inverted setup. The raw images were captured with a 100x oil immersion objective with 0.5 µm step-size, 11 z-slices, in an Andor Sona scMOS camera, and deconvolved using Andor Fusion software. The 3D-deconvolved and max-projected images were used for representation.

The Olympus IX-83 Laser scanning spinning disc Yokogawa-W1 microscope was used to capture **Fig. 3A, 4F, S6A, 8A, and S10A** images. Raw images were acquired with a 100x oil-immersion objective with 1.45 N.A., using 1 µm step-size and five z-slices. The time-lapse imaging for **Fig. S1E, S2D, S4A, S4D, 5A, 5D, S5A, S5B, 6A, 6C, 7D, 8B, S8A-S8C, S9A, and S9B** was acquired at a one-minute time interval with 60x objective with 1.42 N.A. The images and time-lapse montages were 3D-deconvolved in Olympus CellSens Dimension (3.1) software. Maximum***-***intensity projection images were used for representation.

### Protein accumulation kinetics quantification and analysis

The live-cell imaging data were opened in Fiji (ImageJ2) image analysis software^101^ using the protocol established^102^. In brief, a region of interest (ROI) was drawn around the bud neck of the sum-intensity projected image. The extracted fluorescent intensity values were corrected using background values. All the individual fluorescent intensity values obtained in an excel sheet at each time point were normalized using the minimum and maximum values according to the formula below:

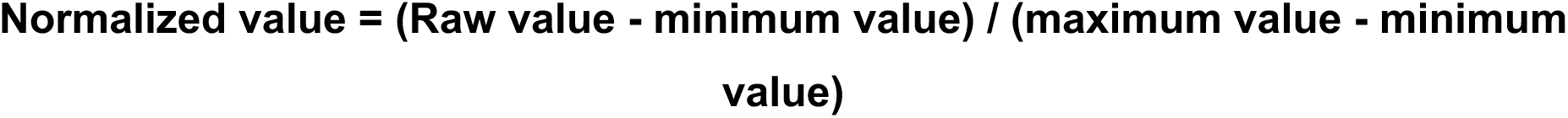

The data points ranging between the -20 and +20 time points were separately extracted for the ‘n’ number of cells indicated with each graph, respectively. The mean and standard deviation were calculated. A line plot was generated using Origin-Pro (2015, Sr2, 69.2.272; Origin Lab, USA) image analysis software to represent the temporal accumulation kinetics of the protein of interest. The small bud is considered as t_0_ for quantifications involving bud emergence. The complete drop in Cdc3-mCherry intensity is considered as the HDR transition in **Figure S1E.** The spindle breakpoint, where the spindle splits during the cytokinesis, is considered as t_0_ or the reference time-point for measuring Hof1, Inn1 and Chs2 temporal dynamics during cytokinesis.

Quantitative analysis of septin and Hof1 misorganization was performed manually by counting the number of cells exhibiting septin mislocalization to the bud cortex during bud emergence and the defective/discontinuous Hof1 ring at the bud neck. The residence time of AMR components at the bud neck during cytokinesis was calculated from the time-lapse movies as the time interval between the first appearance of the signal and its complete disappearance from the bud neck.

### Quantitative analysis of cellular morphology

Brightfield images were acquired using a 100x oil-immersion objective. Cells containing a large bud were selected for analysis. Cell boundaries were manually drawn by selecting a polygonal ROI in Fiji. The marked ROIs were analyzed to obtain measurements of cell area, major axis length, minor axis length, and aspect ratio. The aspect ratio, defined as the ratio of the cell’s major axis to the minor axis, was used as a quantitative parameter to distinguish between round and elliptical cell shapes.

### Yeast Two-Hybrid Assay

Genes or gene fragments of interest were amplified and cloned into NotI restriction enzyme (NEB; Catalogue No. R3189S)-cut pMM5S and pMM6S plasmids consisting of the DNA binding domain of LexA (Lex^DBD^) and activation domain of Gal4 (Gal4^AD^), respectively. The clones were transformed into the yeast strains SGY37 (MATa) and YPH500 (MATalpha). The transformants were selected on S.C. media lacking histidine and leucine, respectively. Mating was performed on YPD plates and was incubated at 30°C for 2 days, followed by replica plating on SC-His-Leu double selection plates. The plates were incubated for 2 days at 30°C, then overlayed with freshly prepared X-Gal mix (500 mM sodium phosphate buffer (pH 7.0), 10% SDS, 1 M KCl, 1 M MgCl^2^, 0.04% X-gal, and 0.4% low-melting agarose) for detection of β-galactosidase activity^103,104^. Interaction of LexA^DBD^-Protein of interest A with Gal4^AD^-Protein of interest B resulted in expression of the lacZ gene encoding β-galactosidase, which converted X-gal to a product with blue color^105^. The plates were scanned and documented after 14-16 hours of incubation with overlay mix.

### Protein expression, purification, and in vitro binding assay

The N-terminal fragment of Hof1 containing the F-BAR domain (1-350 aa) and KA1 fragment of Gin4 (1003-1142 aa) were PCR-amplified from *S. cerevisiae* genomic DNA and cloned into pGEX-5X1 cut with EcoRI restriction enzyme (Catalogue No. R3101S) and pET28a-6His-bdSUMO vector cut with NdeI restriction enzyme (Catalogue No. R0111S), respectively, using NEB Hifi Builder. Both plasmids were transformed into BL21-DE3 E. coli competent cells and were selected on Luria Bertani (LB) plates containing Kanamycin (50 mg/ml) for pGEX-5X1-Hof1^N (1–350)^ and Ampicillin (100 mg/ml) for pET28a-6His-bdSUMO-Gin4^KA1^. LB broth with Kanamycin or Ampicillin was used to grow primary cultures overnight at 37°C with constant shaking at 200 rpm. The primary culture was diluted to 0.1 O.D. in 500 ml LB broth with Kanamycin or Ampicillin. Protein expression was induced with 0.5 mM isopropyl β-D-1-thiogalactopyranoside (IPTG) once the O.D. reached log phase (0.4-0.5). The culture was incubated at 30°C with shaking at 200 rpm for 3-4 hrs. The cells were pelleted at 4°C, 7000 rpm, for 20 min. Media were aspirated and the pellets were washed with cold phosphate-buffered saline (PBS) containing 1 mM phenylmethylsulfonyl fluoride (PMSF) and were stored at -80°C.

### Purification of 6His-bdSUMO-Gin4^KA1^

Pellets were kept on ice for 10 min. and were subjected to lysis with the lysis buffer (50 mM Tris pH 8, 200 mM NaCl, 10% glycerol). The lysed pellets were clarified at 4°C and 15,000xg. The clarified lysate was mixed with Ni-NTA agarose resin (cat. no. 786-940, G-Biosciences), washed with wash buffer (50 mM Tris pH 8, 200 mM NaCl, 10 mM imidazole), and was incubated at 4°C for 2 hrs. The lysate with Ni-NTA resins was then transferred to poly-prep chromatography columns (cat. no. 731-1550, BIO-RAD laboratories Inc.), and was subjected to gradient elution with increasing concentrations of imidazole. The eluted protein fractions were buffer-exchanged with storage buffer (50 mM Tris-HCl pH 7.4, 2000 mM NaCl, 1 mM DTT, and 10% glycerol) using PD Midi Trap G-10 Sephadex columns (GE Healthcare, cat. No. GE28-9180-11). The purified proteins were flash-frozen in liquid nitrogen and were stored at -80°C. The protein concentration was estimated using a spectrophotometer at 280 nm and was analyzed, along with known concentrations of Bovine Serum Albumin (BSA) standards, on Sodium Dodecyl-Sulfate PolyAcrylamide Gel electrophoresis (SDS-PAGE).

### Purification of GST-Hof1^N (1-350)^

Pellets were kept on ice for 10 min. and were subjected to lysis with the lysis buffer (PBS, 0.5 mM EDTA, 1 mM DTT, 1 mg/ml lysozyme, and complete mini-EDTA-free protease inhibitor cocktail tablets). The clarified lysate was incubated with 1% Triton-X-100 for 20 min. on ice and centrifuged at 4°C, 22,000xg, for 30 min. The lysate was then transferred to a tube containing glutathione sepharose-4B resin (cat. no. GE17-0756-01, GE), washed with wash buffer (5x) (PBS, 0.5 mM EDTA, and 1 mM DTT), and was incubated at 4°C for 2-3 hrs. The lysate with glutathione Sepharose resins was then transferred to poly-prep chromatography columns after incubation, washed with wash buffer, and was stored at 4°C.

### In vitro binding assay

GST or GST-Hof1^N (1–350)^ (∼3 µg) bound to glutathione beads were mixed with ∼1 µg 6His-bdSUMO-Gin4^KA1^ and incubated in binding buffer (PBS, 0.2% NP-40) for 1 hour at 4°C. The beads were washed 3-5 times with wash buffer (PBS, 1% NP-40) and were analyzed by western blot using Anti-His (H-3) mouse monoclonal IgG1 (Santa Cruz/#sc-8036/Lot. No. C0421), Anti-GST (B-14) mouse monoclonal IgG1 (Santa Cruz/#sc-138/Lot. No. K1020), and Anti-mouse IgG-HRP (Cell Signalling/#7076S/Lot. No. 36).

### Spot assay

The yeast strains were grown in liquid YPD broth overnight at 23°C. The cultures were diluted to 1.0 O.D. and were serially diluted in the order 10^0^, 10^-1^, 10^-2^, 10^-3^, and 10^-4^. These dilutions were spotted on three YPD plates and incubated at 23°C, 30°C, and 37°C. Growth was monitored, and plates were scanned after 48 hrs of incubation.

The strains used for rescue of *gin4*Δ *hof1*Δ synthetic lethality **(Fig. S3I)** were grown overnight in S.C. media lacking Uracil and Leucine. The culture was diluted to an O.D. of 1.0 and was serially diluted in the order 10^0^, 10^-1^, 10^-2^, 10^-3^, and 10^-4^. These dilutions were spotted on three SC-Ura-Leu plates and were incubated at 23°C for two days. Subsequently, the colonies were patched onto 5-FOA plates (1 mg/ml) to select for Ura***-***cells.

### Statistical analysis

All experiments were performed independently three times. Statistical significance for bar graphs and stacked column plots was determined using one-way ANOVA with Tukey’s multiple-comparison test, whereas residence-time plots were analyzed using the Kruskal-Wallis nonparametric test. All statistical analyses were performed using in-built functions of GraphPad Prism (v.6.04). A p-value of p<0.05 was considered statistically significant. (*: p<0.05, **: p<0.01, ***: p<0.001, ****: p<0.0001, ns: p>0.05).

## Supplementary movie legends

**Movie-S1:** Representative movies showing the recruitment of Elm1, Gin4, Hsl1, and Kcc4 during bud emergence and cytokinesis.

**Movie-S2:** Representative movies showing the localization and dynamics of Cdc3-mCherry in wild-type, *elm1*Δ, *gin4*Δ, *kcc4*Δ, and *hsl1*Δ backgrounds.

**Movie-S3:** Representative movies showing Inn1-mNG constriction in wild-type, *elm1*Δ, *gin4*Δ, *kcc4*Δ, and *hsl1*Δ backgrounds.

**Movie-S4:** Representative movies showing the localization and constriction of Hof1-mNG in wild-type, *elm1*Δ, and *gin4*Δ cells during the cell cycle.

---

**Supplemental Table 1:** Yeast strains used in this study.

| Strain number | Genotype | Source |
| --- | --- | --- |
| YSP002 | *MATa ura3-52 leu2Δ1 trp1Δ63 his3Δ200 (ESM356)* | 106 |
| YSP120 | *ESM356-Elm1-GFP-Hph Cdc3-mCherry-Trp* | This study |
| YSP121 | *ESM356-Gin4-GFP-Hph Cdc3-mCherry-Trp* | This study |
| YSP122 | *ESM356-Kcc4-GFP-Hph Cdc3-mCherry-Trp* | This study |
| YSP123 | *ESM356-Hsl1-GFP-Hph Cdc3-mCherry-Trp* | This study |
| YSP524 | *ESM356-Cdc3-mCherry-Trp, Tub1-GFP-ura* | This study |
| YSP520 | *ESM356-Cdc3-mCherry-Trp, Tub1-GFP-Ura, Δelm1::His3mX6* | This study |
| YSP521 | *ESM356-Cdc3-mCherry-Trp, Tub1-GFP-Ura, Δgin4::His3mX6* | This study |
| YSP1104 | *ESM356-Cdc3-mCherry-Trp, Tub1-GFP-Ura, Δkcc4::His3mX6* | This study |
| YSP518 | *ESM356-Δhsl1-His3mX6, Cdc3-mCherry-Trp, Tub1-GFP-Ura* | This study |
| YSP058 | *ESM356-Δelm1::His3MX6* | This study |
| YSP062 | *ESM356-Δgin4::His3MX6* | This study |
| YSP050 | *ESM356-Δhsl1::His3MX6* | This study |
| YSP115 | *ESM356-Δkcc4::KanMX6* | This study |
| YSP950 | *ESM356-Inn1-mNG-KanMX6, pHis3-mRuby2-Tub1-Ura3* | This study |
| YSP1017 | *ESM356-Inn1-mNG-KanMX6, pHis3-mRuby2-Tub1-Ura3, Δelm1::3HA-Trp* | This study |
| YSP1004 | *ESM356-Inn1-mNG-KanMX6, pHis3-mRuby2-Tub1-Ura3, Δgin4::3HA-Trp* | This study |
| YSP1014 | *ESM356-Inn1-mNG-KanMX6, pHis3-mRuby2-Tub1-Ura3, Δhsl1::3HA-Trp* | This study |
| YSP1037 | *ESM356-Inn1-mNG-Kanmx6, pHis3-mRuby2-Tub1-Ura3, Δkcc4::3HA-Trp* | This study |
| YSP948 | *ESM356-Hof1-mNG-KanMX6, pHis3-mRuby2-Tub1-Ura3* | This study |
| YSP1024 | *ESM356-Hof1-mNG-KanMX6, pHis3-mRuby2-Tub1-Ura3, Δelm1::3HA-Trp* | This study |
| YSP1095 | *ESM356-Hof1-mNG-Kanmx6, pHis3-mRuby2-Tub1-Ura3, Δgin4::3HA-Trp* | This study |
| YSP947 | *ESM356-Chs2-mNG-KanMX6, pHis3-mRuby2-Tub1-Ura3* | This study |
| YSP1019 | *ESM356-Chs2-mNG-KanMX6, pHis3-mRuby2-Tub1-Ura3, Δelm1::3HA-Trp* | This study |
| YSP1002 | *ESM356-Chs2-mNG-KanMX6, pHis3-mRuby2-Tub1-Ura3, Δgin4::3HA-Trp* | This study |
| YSP003 | *MATα ura3-52 lys2-801amber ade2-101ochre trp1Δ63 his3Δ200 leu2Δ1 (YPH500)* | 100 |
| YSP004 | *MATa leu2 ADE2 ura3-52::URA3-lexA-op-LacZ his3 trp (SGY37)* | 100 |
| YSP997 | *ESM356-Cdc3-mCherry-Ura, Δgin4::3HA-Trp* | This study |
| YSP1581 | *ESM356-Cdc3-mCherry-Ura Δgin4::3HA-Trp pRS305-(Vector alone) cut with Kas I and integrated into Leu locus* | This study |
| YSP1582 | *ESM356-Cdc3-mCherry-Ura Δgin4::3HA-Trp pRS305-pTEF-Gin4(FL)-GFP-tCyc1 cut with Kas I and integrated into Leu locus* | This study |
| YSP1583 | *ESM356-Cdc3-mCherry-Ura Δgin4::3HA-Trp pRS305-pTEF-Gin4(K48A)-GFP-tCyc1 cut with Kas I and integrated into Leu locus* | This study |
| YSP1236 | *ESM356-Inn1-mNG-KanMX6, pHis3-mRuby2-Tub1-Ura3, Δgin4::3HA-Trp, pRS305-(Empty vector)-cut with Kas I and integrated into leu locus* | This study |
| YSP1237 | *ESM356-Inn1-mNG-KanMX6, pHis3-mRuby2-Tub1-Ura3, Δgin4::3HA-Trp, pRS305-pGin4-Gin4(FL)-tGin4-cut with Kas I and integrated into leu locus* | This study |
| YSP1238 | *ESM356-Inn1-mNG-KanMX6, pHis3-mRuby2-Tub1-Ura3, Δgin4::3HA-Trp, pRS305-pGin4-Gin4(Kinase dead)-tGin4-cut with Kas I and integrated into leu locus* | This study |
| YSP1312 | *ESM356-Hof1-mNG-Kanmx6, pHis3-mRuby2-Tub1-Ura3, Δgin4::3HA-Trp, pRS305-(Empty vector)-cut with Kas1 and integrated into leu locus* | This study |
| YSP1313 | *ESM356-Hof1-mNG-Kanmx6, pHis3-mRuby2-Tub1-Ura3, Δgin4::3HA-Trp, pRS305-pGin4-Gin4(FL)-tGin4-cut with Kas1 and integrated into leu locus* | This study |
| YSP1314 | *ESM356-Hof1-mNG-Kanmx6, pHis3-mRuby2-Tub1-Ura3, Δgin4::3HA-Trp, pRS305-pGin4-Gin4(Kinase dead)-tGin4-cut with Kas1 and integrated into leu locus* | This study |
| YSP001 | *MATa ura3-52 lys2-801amber ade2-101ochre trp1Δ63 his3Δ200 leu2Δ1 (YPH499)* | 100 |
| YSP1086 | YPH499-Δhof1::TRP1 pRS316-HOF1<br>Δgin4::HIS3MX6 (MFY1474) | <sup>74</sup> |
| YSP1357 | YPH499 Δhof1::TRP1 pRS316-HOF1<br>Δgin4::HIS3MX6, pRS305-Empty Vector-cut with<br>Kas1 and integrated into Leu locus | This study |
| YSP1358 | YPH499 Δhof1::TRP1 pRS316-HOF1<br>Δgin4::HIS3MX6, pRS305-pGin4-Gin4(FL)-tGin4-cut<br>with Kas1 and integrated into Leu locus | This study |
| YSP1359 | YPH499 Δhof1::TRP1 pRS316-HOF1<br>Δgin4::HIS3MX6, pRS305-pGin4-Gin4(K48A)-tGin4-cut with Kas1 and integrated into Leu locus | This study |
| YSP034 | ESM356-Gin4-GFP-Hph | This study |
| YSP1345 | ESM356-GFP-Gin4-GFP-Hph Δelm1::3HA-Trp | This study |
| YSP1419 | ESM356-Gin4-GFP-Hph, Δelm1::3HA-Trp, Hsl1-GBP-KanMX6 | This study |
| YSP1651 | ESM356-Δelm1::3HA-Trp Δgin4::6HA NatNT2 | This study |
| YSP1665 | ESM356-Δelm1::3HA-Trp Δgin4::6HA NatNT2<br>pRS305 cut with Kas I and integrated into Leu locus | This study |
| YSP1666 | ESM356-Δelm1::3HA-Trp Δgin4::6HA NatNT2<br>pRS305-pTEF-Gin4(FL)-GFP-tCyc1 cut with Kas I<br>and integrated into Leu locus | This study |
| YSP1667 | ESM356-Δelm1::3HA-Trp Δgin4::6HA NatNT2<br>pRS305-pTEF-Gin4(K48A)-GFP-tCyc1 cut with Kas<br>I and integrated into Leu locus | This study |
| YSP1658 | ESM356-Δelm1::3HA-Trp Δgin4::6HA NatNT2<br>pRS305 cut with Kas I and integrated into Leu locus<br>Hsl1-GBP-KanMX6 | This study |
| YSP1652 | ESM356-Δelm1::3HA-Trp Δgin4::6HA NatNT2<br>pRS305-pTEF-Gin4(FL)-GFP-tCyc1 cut with Kas I<br>and integrated into Leu locus Hsl1-GBP-KanMX6 | This study |
| YSP1653 | ESM356-Δelm1::3HA-Trp Δgin4::6HA NatNT2<br>pRS305-pTEF-Gin4(K48A)-GFP-tCyc1 cut with Kas<br>I and integrated into Leu locus Hsl1-GBP-KanMX6 | This study |
| YSP1737 | ESM356-Gin4-GFP-Hph, Hsl1-Δka1-GBP-KanMX6 | This study |
| YSP1738 | ESM356-Gin4-GFP-Hph Δelm1:3HA-Trp Hsl1-Δka1-GBP-KanMX6 | This study |
| YSP1590 | ESM356-Gin4-GFP-Hph mRuby-Tub1-Ura | This study |
| YSP1594 | ESM356-Gin4-GFP-Hph Δelm1::3HA-Trp, *pHis3-*<br>*mRuby2-Tub1-Ura3* | This study |
| YSP1411 | *ESM356-Gin4-GFP-Hph, Δelm1::3HA-Trp Shs1-GBP-KanMX6* | This study |
| YSP1415 | *ESM356-Gin4-GFP-Hph, Δelm1::3HA-Trp, Bud4-GBP-KanMX6* | This study |
| YSP1423 | *ESM356-Gin4-GFP-Hph, Δelm1::3HA-Trp, Bni5-GBP-KanMX6* | This study |
| YSP119 | *ESM356-Hsl1-GFP-Hph* | This study |
| YSP1485 | *ESM356-Hsl1-GFP-Hph Δelm1::3HA-Trp* | This study |
| YSP1605 | *ESM356-Hsl1-GFP-Hph Δelm1::3HA-Trp Shs1-GBP-KanMX6* | This study |
| YSP1606 | *ESM356-Hsl1-GFP-Hph Δelm1::3HA-Trp Bud4-GBP-KanMX6* | This study |
| YSP1740 | *ESM356-Hsl1-GFP-Hph, Δelm1::3HA-Trp Kcc4-GBP-KanMX6* | This study |
| YSP1745 | *ESM356-Hsl1-GFP-Hph, Δelm1::3HA-Trp Bni5-GBP-KanMX6* | This study |
| YSP1560 | *ESM356-Hsl1-GFP-Hph, Δelm1::3HA-Trp Gin4-GBP-KanMX6* | This study |
| YSP036 | *ESM356-Kcc4-GFP-Hph* | This study |
| YSP1482 | *ESM356-Kcc4-GFP-Hph Δelm1::3HA-Trp* | This study |
| YSP1734 | *ESM356-Kcc4-GFP-Hph, Δelm1::3HA-Trp Hsl1-GBP-KanMX6* | This study |
| YSP1730 | *ESM356-Kcc4-GFP-Hph, Δelm1::3HA-Trp Shs1-GBP-KanMX6* | This study |
| YSP1732 | *ESM356-Kcc4-GFP-Hph, Δelm1::3HA-Trp Bud4-GBP-KanMX6* | This study |
| YSP1747 | *ESM356-Kcc4-GFP-Hph, Δelm1::3HA-Trp Bni5-GBP-KanMX6* | This study |
| YSP1518 | *ESM356-Kcc4-GFP-Hph Δelm1::3HA-Trp Gin4-GBP-KanMX6* | This study |
| YSP1906 | *ESM356-Kcc4-GFP-Hph pHis3-mRuby2-Tub1-Ura3* | This study |
| YSP1908 | *ESM356-Hsl1-GFP-Hph pHis3-mRuby2-Tub1-Ura3* | This study |
| YSP1910 | *ESM356-Kcc4-GFP-Hph Δelm1::3HA-Trp pHis3-mRuby2-Tub1-Ura3* | This study |
| YSP1912 | *ESM356-Kcc4-GFP-Hph Δgin4::3HA-Trp pHis3-mRuby2-Tub1-Ura3* | This study |
| YSP1914 | *ESM356-Hsl1-GFP-Hph Δelm1::3HA-Trp pHis3-mRuby2-Tub1-Ura3* | This study |
| YSP1916 | *ESM356-Hsl1-GFP-Hph* Δ*gin4::3HA-Trp* *pHis3-mRuby2-Tub1-Ura3* | This study |
| YSP1588 | *ESM356-Elm1-GFP-Hph* *pHis3-mRuby2-Tub1-Ura3* | This study |
| YSP1592 | *ESM356-Elm1-GFP-Hph* Δ*gin4::3HA-Trp* *pHis3-mRuby2-Tub1-Ura3* | This study |
| YSP032 | *ESM356-Elm1-GFP-Hph* | This study |
| YSP1346 | *ESM356-Elm1-GFP-Hph* Δ*gin4::3HA-Trp* | This study |
| YSP1410 | *ESM356-Elm1-GFP-Hph*, Δ*gin4::3HA-Trp* *Shs1-GBP-KanMX6* | This study |
| YSP1414 | *ESM356-Elm1-GFP-Hph*, Δ*gin4::3HA-Trp*, *Bud4-GBP-KanMX6* | This study |
| YSP1418 | *ESM356-Elm1-GFP-Hph*, Δ*gin4::3HA-Trp*, *Hsl1-GBP-KanMX6* | This study |
| YSP1422 | *ESM356-Elm1-GFP-Hph*, Δ*gin4::3HA-Trp*, *Bni5-GBP-KanMX6* | This study |
| YSP1735 | *ESM356-Elm1-GFP-Hph*, *Hsl1-Δka1-GBP-KanMX6* | This study |
| YSP1736 | *ESM356-Elm1-GFP-Hph* Δ*gin4::3HA-Trp* *Hsl1-Δka1-GBP-KanMX6* | This study |
| YSP1591 | *ESM356-Elm1-GFP-Hph* Δ*gin4::3HA-Trp* *Cdc3-mCherry-Ura* | This study |
| YSP1453 | *ESM356-Elm1-GFP-Hph* Δ*gin4::3HA-Trp*, *Shs1-GBP-KanMX6*, *Cdc3-mCherry-Ura* | This study |
| YSP1715 | *ESM356-Elm1-GFP-Hph* Δ*gin4::3HA-trp* Δ*hsl1::natNT1* *Cdc3-mCherry-Ura* | This study |
| YSP1717 | *ESM356-Elm1-GFP-Hph* Δ*gin4::3HA-trp* *Shs1-GBP-KanMX6* Δ*hsl1::natNT1* *Cdc3-mCherry-Ura* | This study |
| YSP1757 | *ESM356-Elm1-GFP-Hph* Δ*gin4::3HA-trp* *Inn1-3xmCherry-His3MX6* | This study |
| YSP1759 | *ESM356-Elm1-GFP-Hph*, Δ*gin4::3HA-Trp* *Shs1-GBP-KanMX6* *Inn1-3xmCherry-His3MX6* | This study |
| YSP1755 | *ESM356-Elm1-GFP-Hph* Δ*gin4::3HA-trp* Δ*hsl1::natNT1* *Inn1-3xmCherry-His3MX6* | This study |
| YSP1754 | *ESM356-Elm1-GFP-Hph* Δ*gin4::3HA-trp* *Shs1-GBP-KanMX6* Δ*hsl1::natNT1* *Inn1-3xmCherry-His3MX6* | This study |
| YSP1724 | *ESM356-Elm1-GFP-Hph* Δ*gin4::3HA-trp* *Myo1-3xmCherry-His3MX6* | This study |
| YSP1726 | *ESM356-Elm1-GFP-Hph*, Δ*gin4::3HA-Trp* *Shs1-GBP-KanMX6* *Myo1-3xmCherry-His3MX6* | This study |
| YSP1722 | *ESM356-Elm1-GFP-Hph* Δ*gin4::3HA-trp*<br>Δ*hsl1::natNT1 Myo1-3xmCherry-His3MX6* | This study |
| YSP1721 | *ESM356-Elm1-GFP-Hph* Δ*gin4::3HA-trp Shs1-GBP-KanMX6* Δ*hsl1::natNT1 Myo1-3xmCherry-His3MX6* | This study |
| YSP1417 | *ESM356-Gin4-GFP-Hph, Hsl1-GBP-KanMX6* | This study |
| YSP1589 | *ESM356-Gin4-GFP-Hph Cdc3-mCherry-Ura* | This study |
| YSP1593 | *ESM356-Gin4-GFP-Hph* Δ*elm1::3HA-Trp Cdc3-mCherry-Ura* | This study |
| YSP1595 | *ESM356-Gin4-GFP-Hph* Δ*elm1::3HA-Trp Hsl1-GBP-KanMX6 Cdc3-mCherry-Ura* | This study |
| YSP1599 | *ESM356-Gin4-GFP-Hph Hsl1-GBP-KanMX6 Cdc3-mCherry-Ura* | This study |
| YSP1489 | *ESM356-Gin4-GFP-Hph Hsl1-GBP-KanMX6 Myo1-ymScarlet-I* | This study |
| YSP1491 | *ESM356-Gin4-GFP-Hph* Δ*elm1::3HA-Trp Hsl1-GBP-KanMX6 Myo1-ymScarlet-I* | This study |
| YSP1627 | *ESM356-Gin4-GFP-Hph Myo1-ymScarlet-I-Nat* | This study |
| YSP1629 | *ESM356-Gin4-GFP-Hph* Δ*elm1::3HA-Trp Myo1-ymScarlet-I-Nat* | This study |
| YSP951 | *ESM356-Cdc3-mCherry-Ura* | This study |
| YSP1033 | *ESM356-Cdc3-mCherry-Ura, Δelm1::3HA-Trp* | This study |
| YSP2374 | *ESM356-Cdc3-mCherry-Ura Δswe1::His3MX6* | This study |
| YSP2376 | *ESM356-Gin4-GFP-Hph* Δ*elm1::3HA-Trp Cdc3-mCherry-Ura Δswe1::His3MX6* | This study |
| YSP2377 | *ESM356-Gin4-GFP-Hph* Δ*elm1::3HA-Trp Hsl1-GBP-KanMX6 Cdc3-mCherry-Ura Δswe1::His3MX6* | This study |
| YSP2380 | *ESM356-Cdc3-mCherry-Ura, Δelm1::3HA-Trp Δswe1::His3MX6* | This study |
| YSP1348 | *ESM356-Gin4-Δka1-GFP-mRuby-Tub1-Ura* |  |
| YSP1590 | *ESM356-Gin4-GFP-Hph mRuby-Tub1-Ura* | This study |
| YSP1207 | *ESM356-Hof1-mNG, mRuby-Tub1, Gin4-Δka1::3HA-Trp* | This study |

**Supplemental Table 2:**
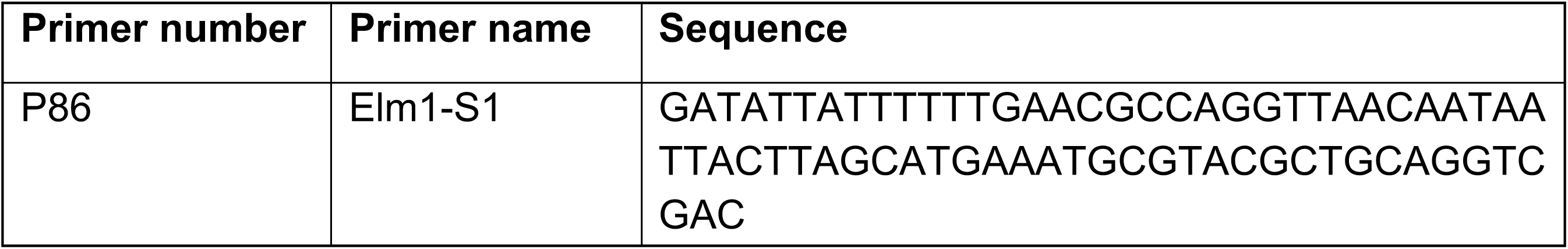
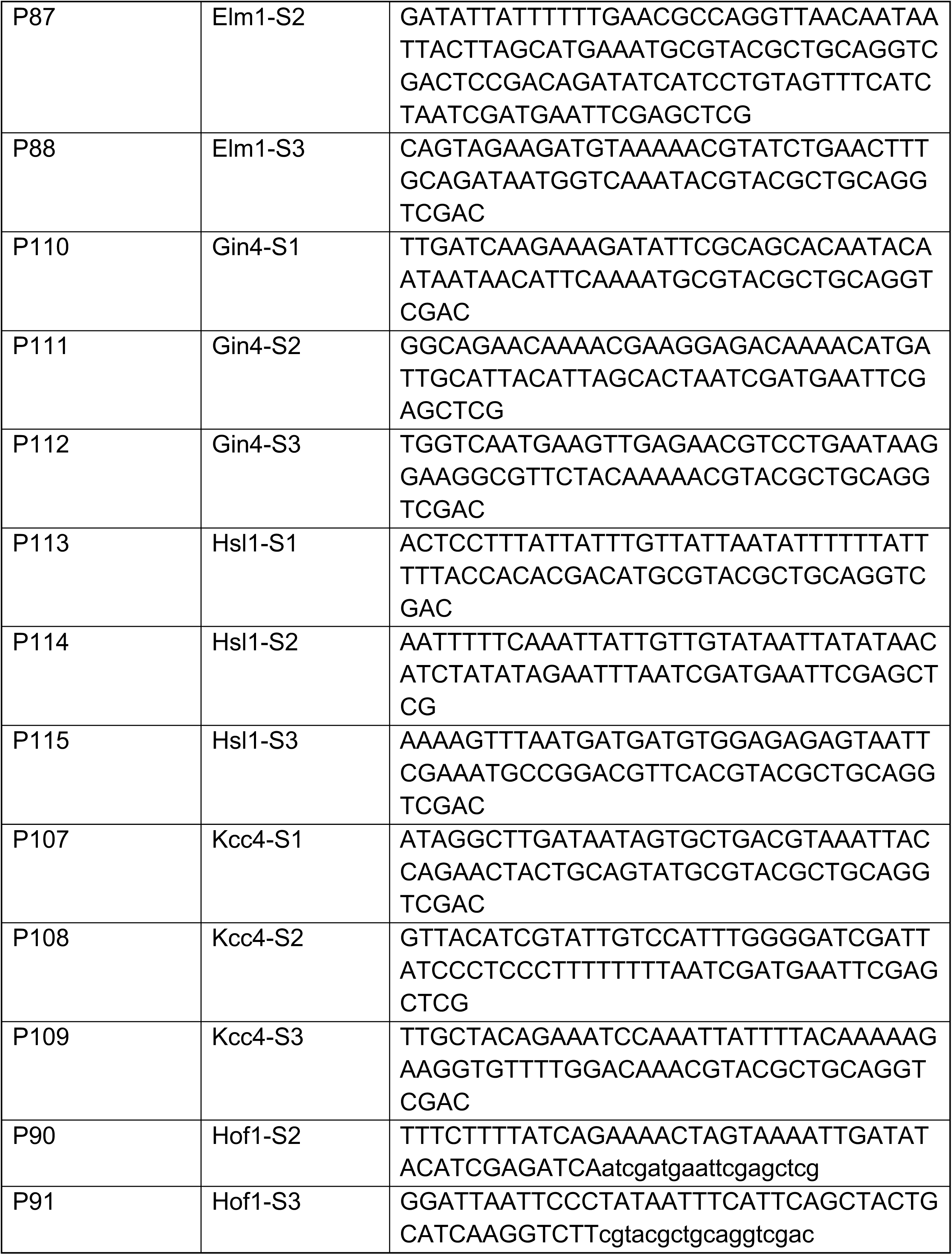
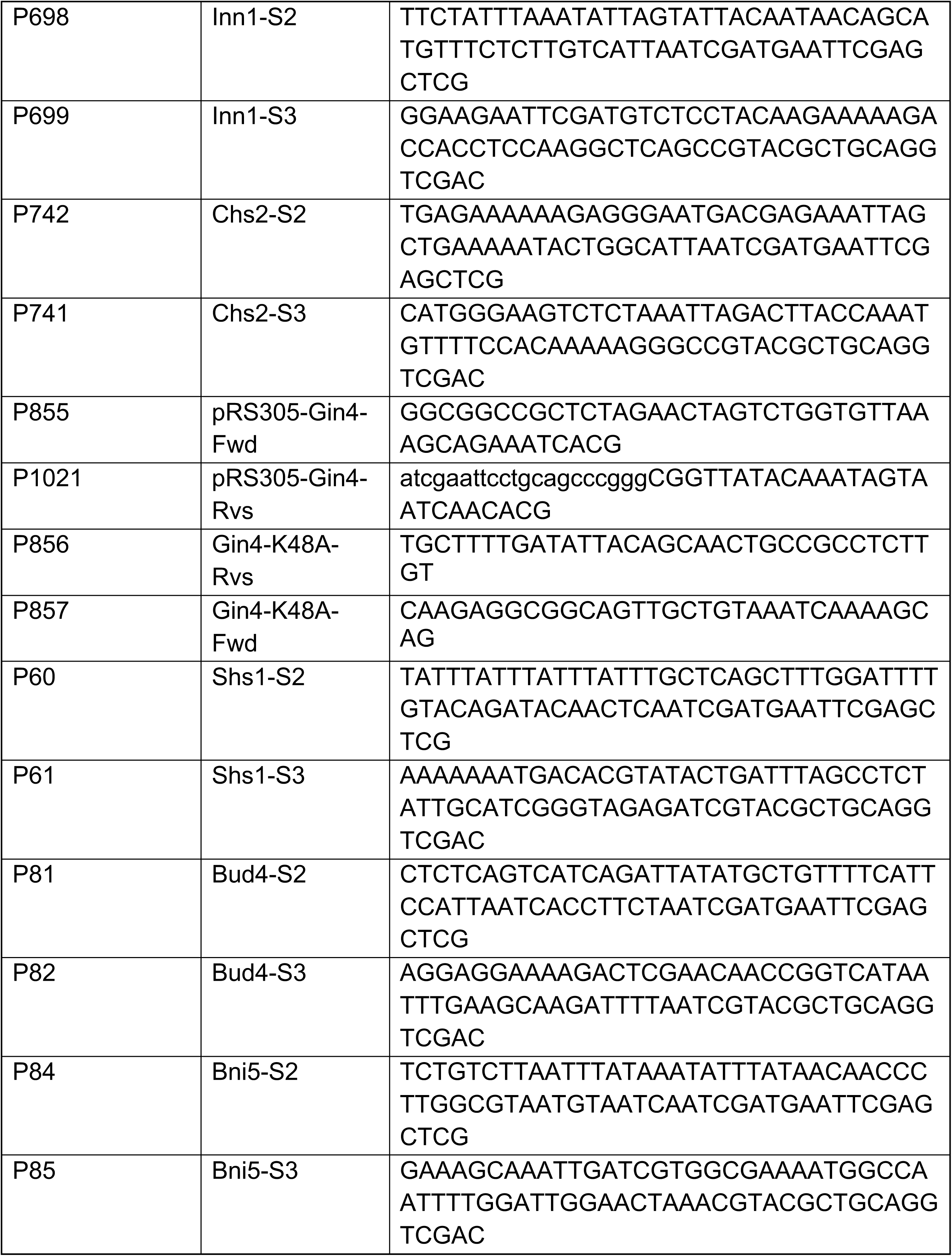
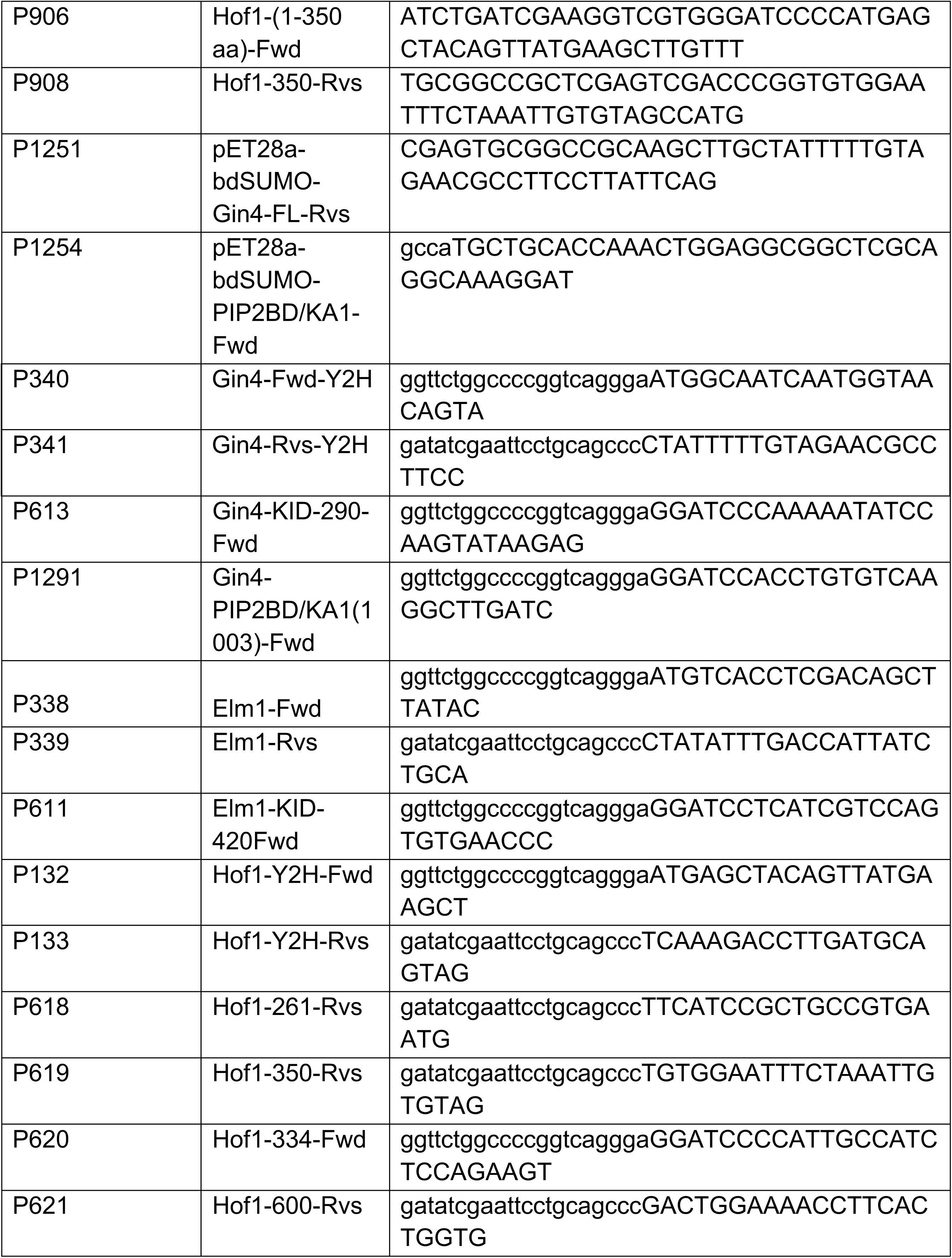
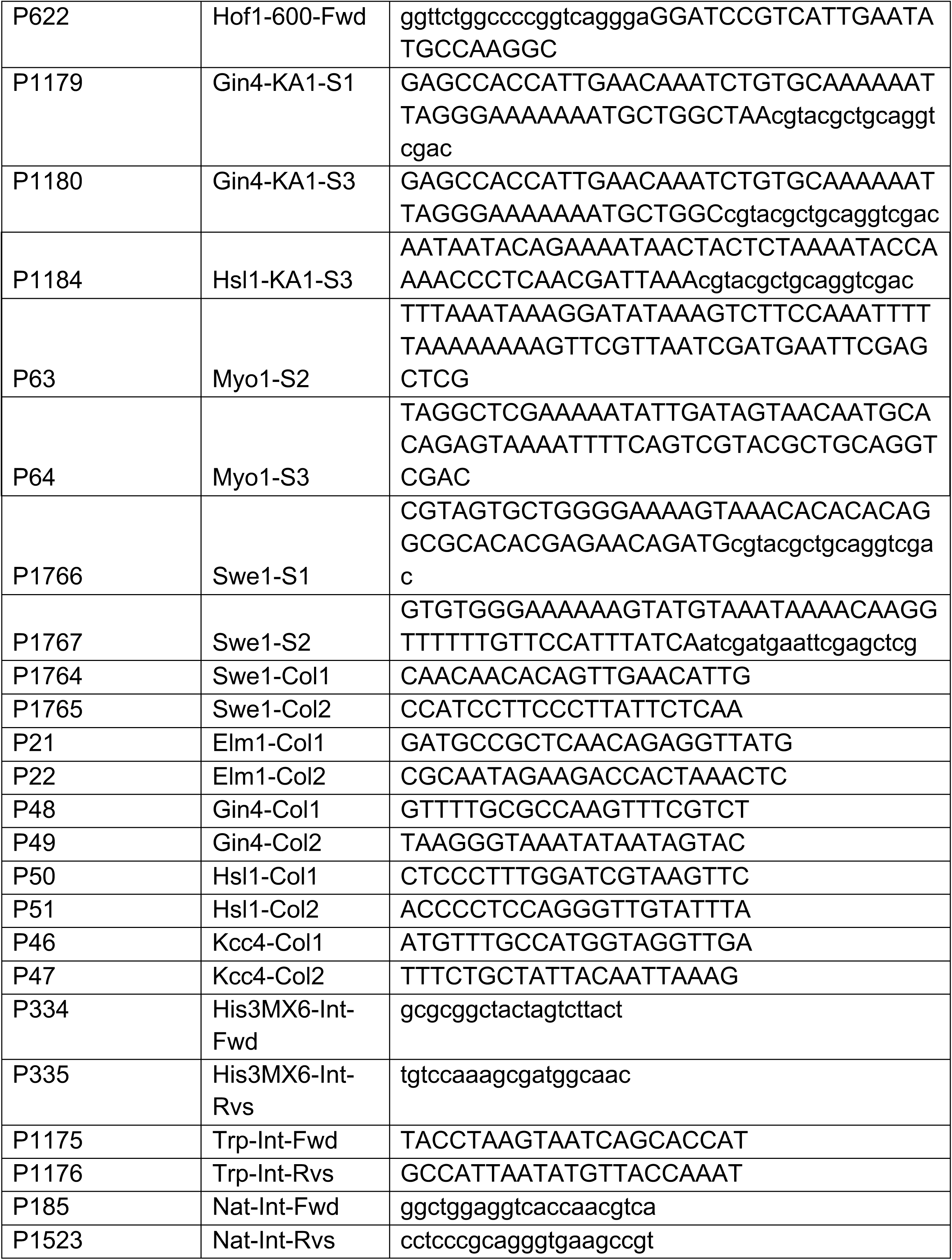
Oligonucleotides used in this study.

**Supplemental Table 3:** Plasmids used in this study.

| Plasmid number | Description | Source |
| --- | --- | --- |
| piSP23 | pFA6a-HIS3-MX6 | 96 |
| piSP14 | pYM25 (yeGFP-hphNT1) | 95 |
| piSP20 | pYM22 (3HA-klTRP1) | 95 |
| piSP185 | pYM17 (6HA-natNT2) | 96 |
| piSP659 | pDK88 (mNeongreen cassette. mNG-kanmx6) | Gift from Prof. Gislene Pereira |
| piSP1075 | pHA29-GBP-KAN | 107 |
| piSP31 | YIP211-cdc3-mCherry (URA3) | 72 |
| piSP34 | YIP204-cdc3-mCherry (TRP1) | 77 |
| piSP678 | pAFS125-GFP-TUB1 | 99 |
| piSP1224 | pHIS3p:mRuby2-Tub1+3"UTR::Ura3 | 98 |
| piSP1 | pRS305 | 100 |
| piSP1341 | pRS305-pGin4-Gin4(FL)-tGin4 | This Study<sup>a</sup> |
| piSP1739 | pRS305-pGin4-Gin4(K48A)-tGin4 | This Study<sup>a</sup> |
| piSP1614 | pRS305-pTEF-Gin4-GFP-tCYC1 | This Study<sup>b</sup> |
| piSP1704 | pRS305-pTEF-Gin4 (K48A)-GFP-tCYC1 | This Study<sup>b</sup> |
| piSP656 | pBPE39-1(3xmCherry-His3MX6 cassette) | Gift from Prof. Gislene Pereira |
| piSP112 | pMM5S-Y2H (Empty vector) | 108 |
| piSP113 | pMM6S-Y2H (Empty vector) | 108 |
| piSP635 | pMM5S-Gin4-FL (1-1142aa) | This Study |
| piSP645 | pMM6S-Gin4-FL (1-1142aa) | This Study |
| piSP632 | pMM5S-Elm1-FL (1-640aa) | This Study |
| piSP642 | pMM6S-Elm1-FL (1-640aa) | This Study |
| piSP685 | pMM5S-Gin4-KID (290-1142aa) | This Study |
| piSP691 | pMM6S-Gin4-KID (290-1142aa) | This Study |
| piSP683 | pMM5S-Elm1-KID (421-640aa) | This Study |
| piSP689 | pMM6S-Elm1-KID (421-640aa) | This Study |
| piSP568 | pMM5S-Hof1-FL (1-669aa) | This Study |
| piSP580 | pMM6S-Hof1-FL (1-669aa) | This Study |
| piSP1275 | pMM5S-Hof1(1-350aa) | This Study |
| piSP1277 | pMM6S-Hof1(1-350aa) | This Study |
| piSP1279 | pMM5S-Hof1(334-600aa) | This Study |
| piSP1281 | pMM6S-Hof1(334-600aa) | This Study |
| piSP1283 | pMM5S-Hof1(600-669aa) | This Study |
| piSP1285 | pMM6S-Hof1(600-669aa) | This Study |
| piSP1839 | pMM5S-Gin4-KA1(1003-1142aa) | This Study |
| piSP1844 | pMM6S-Gin4-KA1(1003-1142aa) | This Study |
| piSP65 | pET28a-6HIS-bdSUMO | 109 |
| piSP1094 | pGEX-5X1 | GE Healthcare, <sup>110</sup> |
| piSP1320 | pGEX-5X1-Hof1(1-350) | This Study |
| piSP1712 | pET28a-6HIS-bdSUMO-Gin4(KA1) | This Study |

<sup>a & b</sup> plasmids piSP1341, piSP1739, piSP1614 and piSP1704 were cut with KasI restriction enzyme and integrated into *Leu* locus.

