## Supplementary material for "Nim1-related kinases regulate septin organization and cytokinesis by modulating Hof1 at the cell division site": Bhojappa et al_Supplementary figures_legend

**Supplementary figure legends:**

**Figure S1: Dynamics of septin-associated kinases exhibiting simultaneous recruitment during bud emergence and ordered disassembly during the HDR transition. (A)** Representative time-lapse images and normalized fluorescence intensity graph showing the temporal kinetics of Elm1-GFP during bud-emergence ($t_0$) (n=18 cells). **(B)** Representative time-lapse images and normalized fluorescence intensity graph showing the temporal kinetics of Gin4-GFP during bud-emergence ($t_0$) (n=16 cells). **(C)** Representative time-lapse images and normalized fluorescence intensity graph showing the temporal kinetics of Hsl1-GFP during bud-emergence ($t_0$) (n=18 cells). **(D)** Representative time-lapse images and normalized fluorescence intensity graph for temporal kinetics of Kcc4-GFP during bud-emergence ($t_0$) (n=16 cells). Images in panels S1A-S1D correspond to differential intensity contrast. **(E)** Representative bud neck montages of Elm1-GFP, Gin4-GFP, Hsl1-GFP and Kcc4-GFP during HDR transition ($t_0$= Cdc3-mCherry splitting), DC*=Differential contrast. **(F)** Quantitative analysis of raw fluorescence intensity showing the dynamics of septin kinases indicated in (E) during septin remodelling. The Cdc3-mCherry profile of Kcc4 strain is plotted as a reference for septin remodeling dynamics during cytokinesis. (Elm1-GFP: n=22, Gin4-GFP: n=26, Hsl1-GFP: n=17 and Kcc4-GFP: n=22 cells). **(G)** Plot of normalized fluorescence intensity at the bud neck for the indicated strains shown in (E). The Cdc3-mCherry profile of the Kcc4 strain is plotted as a reference for septin remodelling.

**Figure S2: Deletion of septin-associated kinases differentially perturbs cellular morphology and septin organization. (A)** Brightfield images showing the cellular morphological defects in *elm1*Δ, *gin4*Δ, *hsl1*Δ and *kcc4*Δ cells at 23℃, 30℃ and 37℃ temperatures. Scale bar-5µm. **(B)** Stacked column graphs showing the percentage of cells exhibiting normal and abnormal cell shape in the indicated strains shown in (A) (N=3, n=300 cells/strain). **(C)** Growth assay of the indicated strains shown in (A) at 23℃, 30℃ and 37℃. Representative images were captured after 48 hours of incubation at indicated temperatures. **(D)** Representative montage showing the temporal kinetics of Cdc3-mCherry at one-minute time intervals during mitotic spindle break ($t_0$). Scale bar-5µm. **(E)** Images representing the normal and mislocalization of Cdc3-mCherry during bud emergence in the indicated strains shown in (1B). White arrows depict mislocalized septins in *elm1*Δ and *gin4*Δ strains. Scale bar-5µm. **(F)** Normalized fluorescence intensity graph showing the kinetics of Cdc3-mCherry during cytokinesis in the represented strain shown in (D). **(G)** Stacked column graph showing the percentage of cells exhibiting normal localization and mislocalization of Cdc3-mCherry to the bud cortex in the indicated strains of (E) (wild-type: n=45, *elm1*Δ: n=48, *gin4*Δ: n=41, *hsl1*Δ: n=56, *kcc4*Δ: n=50 cells).

**Figure S3: Gin4 regulates Hof1 organization and cell viability in a kinase-independent manner. (A)** Representative images showing the localization profile of Myo1-mNG in *elm1*Δ and *gin4*Δ cells. White arrows indicate the mislocalized Myo1 in the represented strains. Scale bar-5µm. **(B)** Stacked column graph showing the percentage of cells exhibiting normal localization and mislocalization of Myo1 in the indicated strains of (A) (wild-type: n=26, *elm1*Δ: n=29 and *gin4*Δ: n=30 cells). **(C)** Representative time-lapse images of Chs2-mNG in wild-type, *elm1*Δ and *gin4*Δ cells imaged at two-minute time intervals ($t_0$=appearance of Chs2 at the bud neck). **(D)** Quantification of the residence time of Chs2-mNG at the bud neck in indicated strains shown in (C), Kruskal-Wallis nonparametric test (**: $p<0.01$, ns: $p>0.05$). **(E)** Plot of normalized fluorescence intensity of Chs2-mNG in indicated strains of (C). A population *elm1*Δ and *gin4*Δ cells exhibiting asymmetric constriction is plotted in the graph ($t_0$=spindle breakpoint) (wild-type: n=40, *elm1*Δ: n=9, *gin4*Δ: n=23 cells). **(F)** Representative images showing rescue of Hof1-misorganization at the cell division site upon expression of $Gin4^{FL}$ and $Gin4^{KD}$ constructs cloned under the endogenous promoter in *gin4*Δ cells. Scale bar-5µm. **(G)** Graph representing the percentage of cells exhibiting normal organization and misorganization of Hof1 rings in the indicated strains shown in (F), one-way ANOVA with Tukey's multiple-comparison test (****: $p<0.0001$, ns: $p>0.05$. p-values correspond to the population of cells exhibiting disorganized Hof1 at the bud neck (N=3, wild-type: n=284, *gin4*Δ-Empty vector: n=138, *gin4*Δ-$Gin4^{FL}$: n=148, *gin4*Δ-$Gin4^{KD}$: n=113 cells). **(H)** Representative images from the Yeast Two-Hybrid assay depicting the interaction between Gin4-KA1 domain and different domains of Hof1. **(J)** Spot assay showing rescue of synthetic lethality upon expression of $Gin4^{FL}$ and $Gin4^{KD}$ constructs cloned under the endogenous promoter in *gin4*Δ *hof1*Δ cells. Plates were incubated at 23℃ and scanned after 72 hours.

**Figure S4: Gin4-*ka1*Δ-GFP exhibits reduced localization to the bud neck prior to cytokinesis and recapitulates Hof1 defects observed in *gin4*Δ cells. (A)** Bud neck montages representing the temporal kinetics of Gin4-GFP in comparison with Gin4-*ka1*Δ-GFP ($t_0$=spindle breakpoint). **(B)** Plot depicting the raw fluorescence intensity profile of the indicated strains shown in (A) (Gin4-GFP: n=29 and Gin4-*ka1*Δ-GFP: n=29 cells). **(C)** Temporal kinetics graph showing normalized fluorescence intensity for the indicated strains shown in (A). **(D)** Representative montages showing the temporal kinetics of Hof1-mNG during cytokinesis at one-minute time intervals in *gin4*Δ and Gin4-*ka1*Δ cells ($t_0$=spindle breakpoint). **(E)** Quantification of the residence time of Hof1-mNG after spindle breakpoint in the indicated strains shown in (D), Kruskal-Wallis nonparametric test (****: $p<0.0001$, ns: $p>0.05$). **(F)** Temporal kinetics graph of the normalized fluorescence intensity of Hof1-mNG during cytokinesis in the indicated strains shown in (D) (wild-type: n=30, *gin4*Δ: n=29 and Gin4-*ka1*Δ: n=30 cells).

**Figure S5: Targeted localization of Gin4-GFP to the bud neck in *elm1*Δ cells via different bud neck proteins. (A)** Representative montages showing the recruitment of Gin4-GFP during bud emergence ($t_0$) in *elm1*Δ background. Scale bar-5µm. **(B)** Representative montages of the bud neck depicting the localization of Gin4-GFP in *elm1*Δ cells during cytokinesis ($t_0$=spindle breakpoint), DC*= Differential contrast. **(C)** Graph showing the raw fluorescence intensity of Gin4-GFP in the indicated strains shown in (B) (wild-type: n=29 and *elm1*Δ: n=31 cells). **(D)** Plot of normalized fluorescence signal intensity of Gin4-GFP in the indicated strains shown in (B) ($t_0$=spindle breakpoint). **(E)** Representative images showing artificial tethering of Gin4-GFP via Shs1-GBP, Bud4-GBP and Bni5-GBP in Δ*elm1* cells. Scale bar-5µm. **(F)** Bar graph showing the percentage of cells exhibiting round morphology in the indicated strains shown in (E), one-way ANOVA with Tukey's multiple-comparison test (****: p<0.0001, ns: p>0.05) (N=3, wild-type: n=407, *elm1*Δ: n=508, *elm1*Δ-Shs1-GBP: n=488, *elm1*Δ-Bud4-GBP: n=471 and *elm1*Δ-Bni5-GBP: n=448 cells). **(G)** Quantification of the aspect ratios in the indicated strains shown in (E), Kruskal-Wallis nonparametric statistical test (*: p<0.05), (N=3, n>155 cells/strain).

**Figure S6: Absence of Swe1 rescues morphological defects observed in *elm1*Δ cells. (A)** Brightfield images representing cellular morphology in the *elm1*Δ and *elm1*Δ *swe1*Δ cells in which Gin4-GFP is artificially tethered to bud neck via Hsl1-GBP. Scale bar-5µm. **(B)** Bar graph representing the percentage of cells exhibiting round morphology in the indicated strains shown in (A), one-way ANOVA with Tukey's multiple-comparison test (**: p<0.01, ****: p<0.0001, ns: p>0.05) (N=3, wild-type: n=706, *swe1*Δ: n=540, *elm1*Δ: n=436, *elm1*Δ *swe1*Δ: n=638, *elm1*Δ-Gin4-GFP: n=450, *elm1*Δ *swe1*Δ-Gin4-GFP: n=465, *elm1*Δ-Gin4-GFP-Hsl1-GBP: n=623, *elm1*Δ *swe1*Δ-Gin4-GFP-Hsl1-GBP: n=604 cells). **(C)** Quantification of the aspect ratios in the indicated strains shown in (A), Kruskal-Wallis nonparametric statistical test (****: p<0.0001, ns: p>0.05), (N=3, n>170 cells/strain).

**Figure S7: Redirecting Kcc4-GFP localization to bud neck via Hsl1-GBP rescues defects associated with *elm1*Δ cells. (A)** Representative images showing artificial tethering of Kcc4-GFP to the bud neck via Shs1-GBP, Bud4-GBP, Bni5-GBP, Gin4-GBP and Hsl1-GBP in *elm1*Δ cells. Scale bar-5µm. **(B)** Bar graph showing the percentage of cells exhibiting round morphology in the indicated strains shown in (A), one-way ANOVA with Tukey's multiple-comparison test (*: p<0.05, **: p<0.01, ****: p<0.0001, ns: p>0.05) (N=3, wild-type: n=724, *elm1*Δ: n=491, *elm1*Δ-Shs1-GBP: n=587, *elm1*Δ-Bud4-GBP: n=663, *elm1*Δ-Bni5-GBP: n=569, *elm1*Δ-Gin4-GBP: n=483 and *elm1*Δ-Hsl1-GBP: n=708 cells). **(C)** Quantification of aspect ratios in the indicated strains shown in (A), Kruskal-Wallis nonparametric statistical test (***: p<0.001, ****: p<0.0001, ns: p>0.05), (N=3, n>155 cells/strain).

**Figure S8: Kcc4 is recruited to the bud neck in an Elm1-dependent and Gin4-independent manner. (A)** Representative montages showing the recruitment of Hsl1-GFP during bud emergence ($t_0$) in *elm1*Δ and *gin4*Δ backgrounds. DC*=Differential Contrast. Scale bar-5µm. **(B)** Representative montages showing the recruitment of Kcc4-GFP during bud emergence ($t_0$) in *elm1*Δ and *gin4*Δ backgrounds. DC*=Differential Contrast. Scale bar-5µm. **(C)** Representative montages of the bud neck depicting the localization of Kcc4-GFP in *elm1*Δ and *gin4*Δ cells during cytokinesis. **(D)** Plot of raw fluorescence intensity of Kcc4-GFP in the indicated strains shown in (C) ($t_0$=spindle breakpoint) (wild-type: n=30, *elm1*Δ: n=26 and *gin4*Δ: n=30 cells). **(E)** Plot of normalized fluorescence intensity of Kcc4-GFP in indicated strains shown in (C) ($t_0$=spindle breakpoint).

**Figure S9: Hsl1 kinase becomes essential downstream of Elm1 to regulate cytokinesis in *gin4*Δ cells.** **(A)** Representative montages showing the recruitment of Elm1-GFP during bud emergence ($t_0$) in *gin4*Δ background. Scale bar-5µm. **(B)** Representative montages of the bud neck depicting the localization of Elm1-GFP during cytokinesis in *gin4*Δ cells. DC*=Differential contrast. **(C)** Plot representing the raw fluorescence intensity of Elm1-GFP in the indicated strains shown in (B) ($t_0$=spindle breakpoint) (wild-type: n=26 and *gin4*Δ: n=32 cells). **(D)** Plot of normalized fluorescence intensity of Elm1-GFP in indicated strains of (B) ($t_0$=spindle breakpoint). **(E)** Representative images showing artificial tethering of Elm1-GFP via Shs1-GBP, Bud4-GBP and Hsl1-GBP in *gin4*Δ cells. Scale bar-5µm. **(F)** Bar graph showing the percentage of cells exhibiting round morphology in the indicated strains shown in (E), one-way ANOVA with Tukey's multiple-comparison test (*: p<0.05, **: p<0.01, ***: p<0.001, ns: p>0.05) (N=3, wild-type: n=389, *gin4*Δ: n=415, *gin4*Δ-Shs1-GBP: n=800, *gin4*Δ-Bud4-GBP: n=688 and *gin4*Δ-Hsl1-GBP: n=886 cells). **(G)** Quantification of aspect ratios in the indicated strains shown in (E), Kruskal-Wallis nonparametric statistical test (**: p<0.01, ****: p<0.0001, ns: p>0.05) (N=3, n>165 cells/strain). **(H)** Representative images showing artificial tethering of Elm1-GFP with Hsl1-Δ*ka1*-GBP in *gin4*Δ cells. Scale bar-5µm. **(I)** Bar graph showing percentage of cells exhibiting round and elongated/clumped morphologies in the indicated strains shown in (H), one-way ANOVA with Tukey's multiple-comparison test (****: p<0.0001, ns: p>0.05), p-values correspond to population of cells with round morphology, (N=3, wild-type: n=765, Hsl1-GBP: n=663, *gin4*Δ: n=602 and *gin4*Δ-Hsl1-GBP: n=609 cells). **(J)** Quantification of aspect ratios in the indicated strains shown in (H), Kruskal-Wallis nonparametric statistical test (****: p<0.0001, ns: p>0.05) (N=3, n>165 cells/strain).

**Figure S10: Absence of Hsl1 exacerbates the mislocalization of Myo1 to bud cortex in *gin4*Δ cells.** **(A)** Representative images showing mislocalization of Myo1-3xmCherry to the bud cortex in *gin4*Δ and *gin4*Δ *hsl1*Δ cells in which Elm1-GFP is artificially tethered to the bud neck via Shs1-GBP. White arrows indicate mislocalized Myo1-3xmCherry in the represented strains. Scale bar-5µm. **(B)** Stacked column graph showing the percentage of cells exhibiting normal localization and mislocalization of Myo1-3xmCherry during bud emergence in the indicated strains shown in (A) ( *gin4*Δ: n=39, *gin4*Δ-Shs1-GBP: n=36, *gin4*Δ *hsl1*Δ: n=38 and *gin4*Δ *hsl1*Δ-Shs1-GBP: n=39 cells).

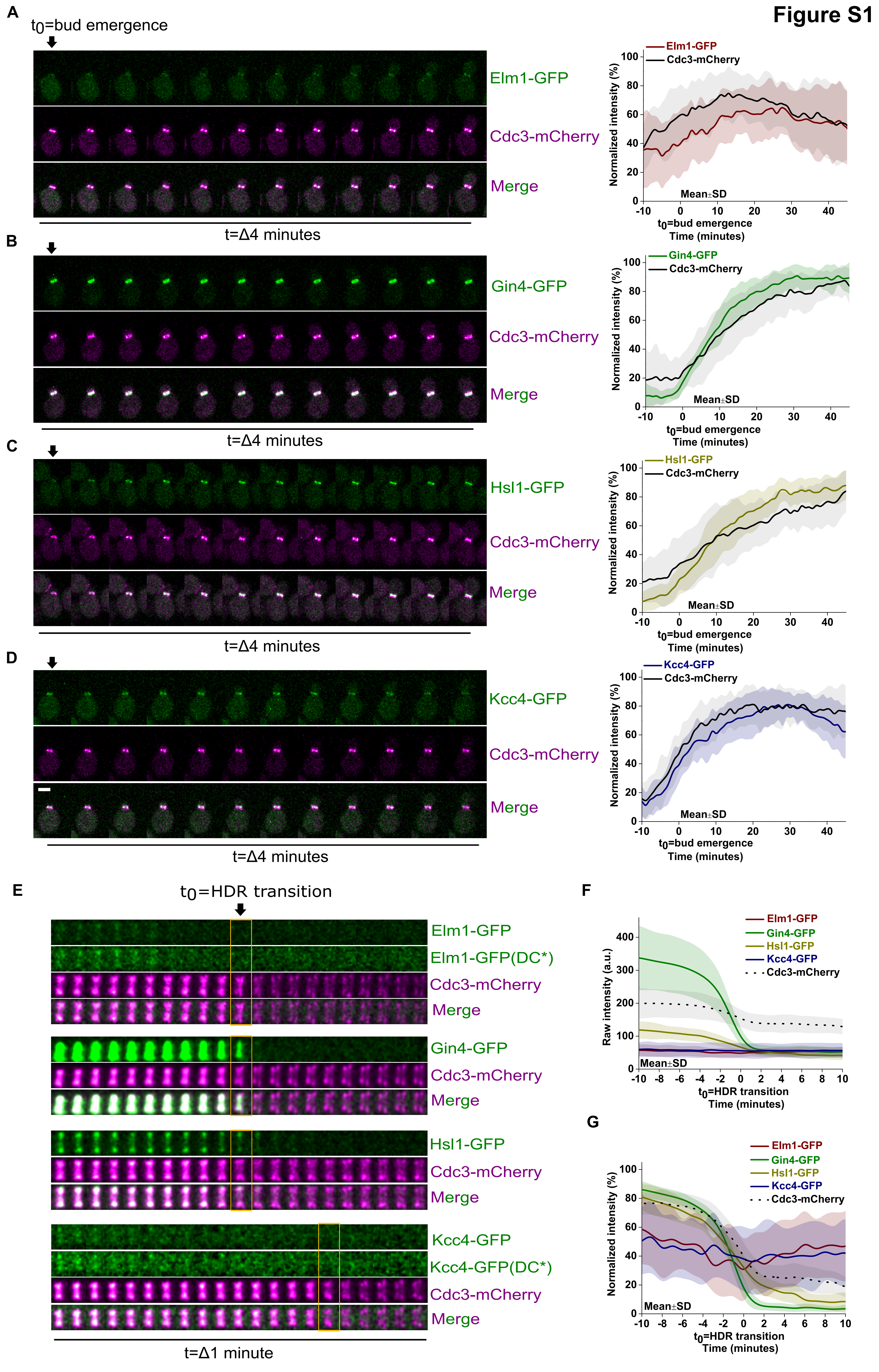
Figure S1
A
t0=bud emergence
Elm1-GFP
Cdc3-mCherry
Merge
t=Δ4 minutes
Mean±SD
Normalized intensity (%)
t0=bud emergence
Time (minutes)
B
Gin4-GFP
Cdc3-mCherry
Merge
t=Δ4 minutes
C
Hsl1-GFP
Cdc3-mCherry
Merge
t=Δ4 minutes
D
Kcc4-GFP
Cdc3-mCherry
Merge
t=Δ4 minutes
E
t0=HDR transition
Elm1-GFP
Elm1-GFP(DC*)
Cdc3-mCherry
Merge
Gin4-GFP
Hsl1-GFP
Kcc4-GFP
Kcc4-GFP(DC*)
t=Δ1 minute
F
Raw intensity (a.u.)
t0=HDR transition
Time (minutes)
G
Normalized intensity (%)

#### Figure S2

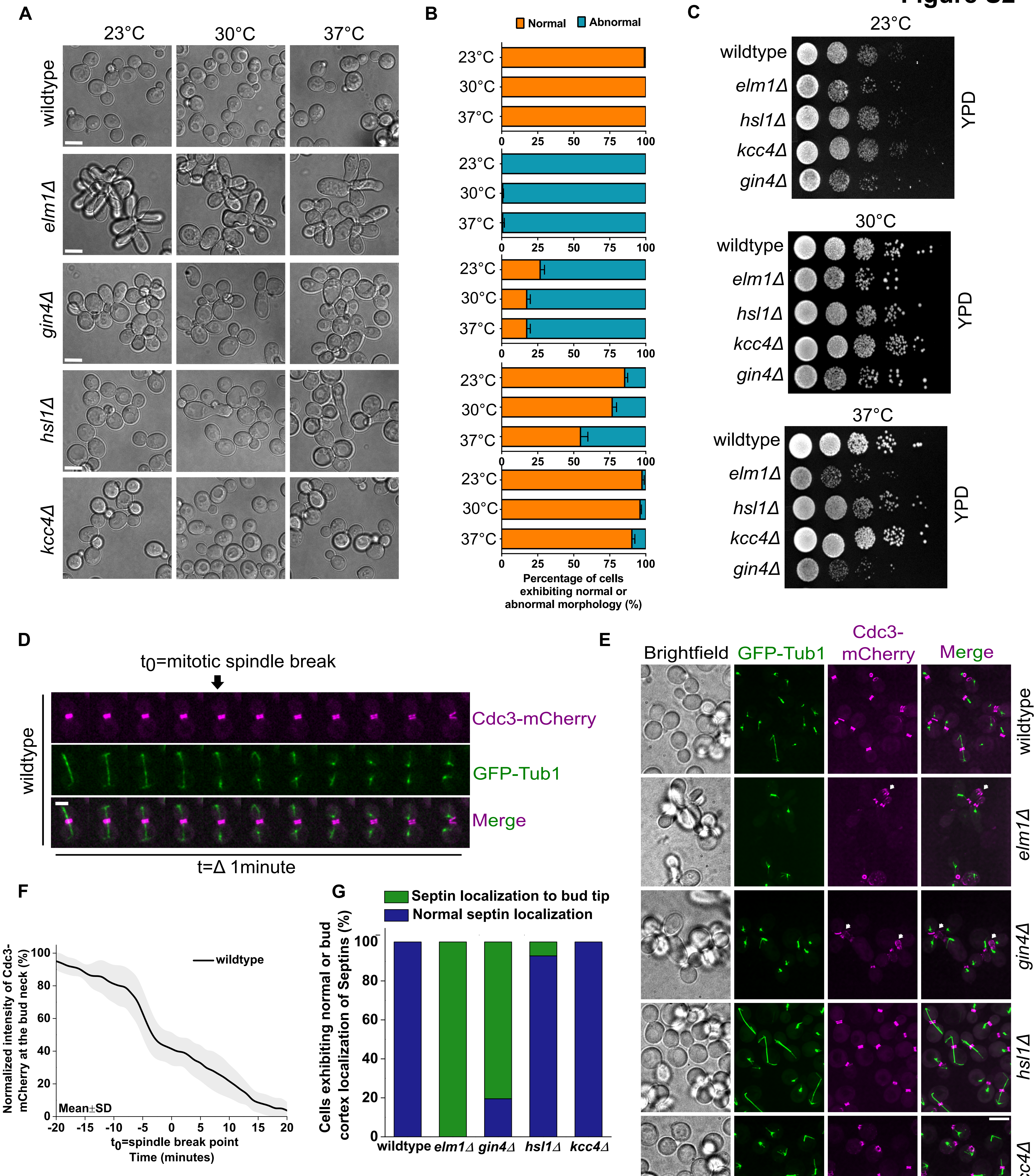

A
23°C
30°C
37°C
wildtype
elm1Δ
gin4Δ
hsl1Δ
kcc4Δ
B
Normal
Abnormal
Percentage of cells exhibiting normal or abnormal morphology (%)
C
23°C
30°C
37°C
YPD
D
t0=mitotic spindle break
Cdc3-mCherry
GFP-Tub1
Merge
t=Δ 1minute
E
Brightfield
GFP-Tub1
Cdc3-mCherry
Merge
F
wildtype
Mean±SD
Normalized intensity of Cdc3-mCherry at the bud neck (%)
t0=spindle break point
Time (minutes)
G
Septin localization to bud tip
Normal septin localization
Cells exhibiting normal or bud cortex localization of Septins (%)

**Figure S3**

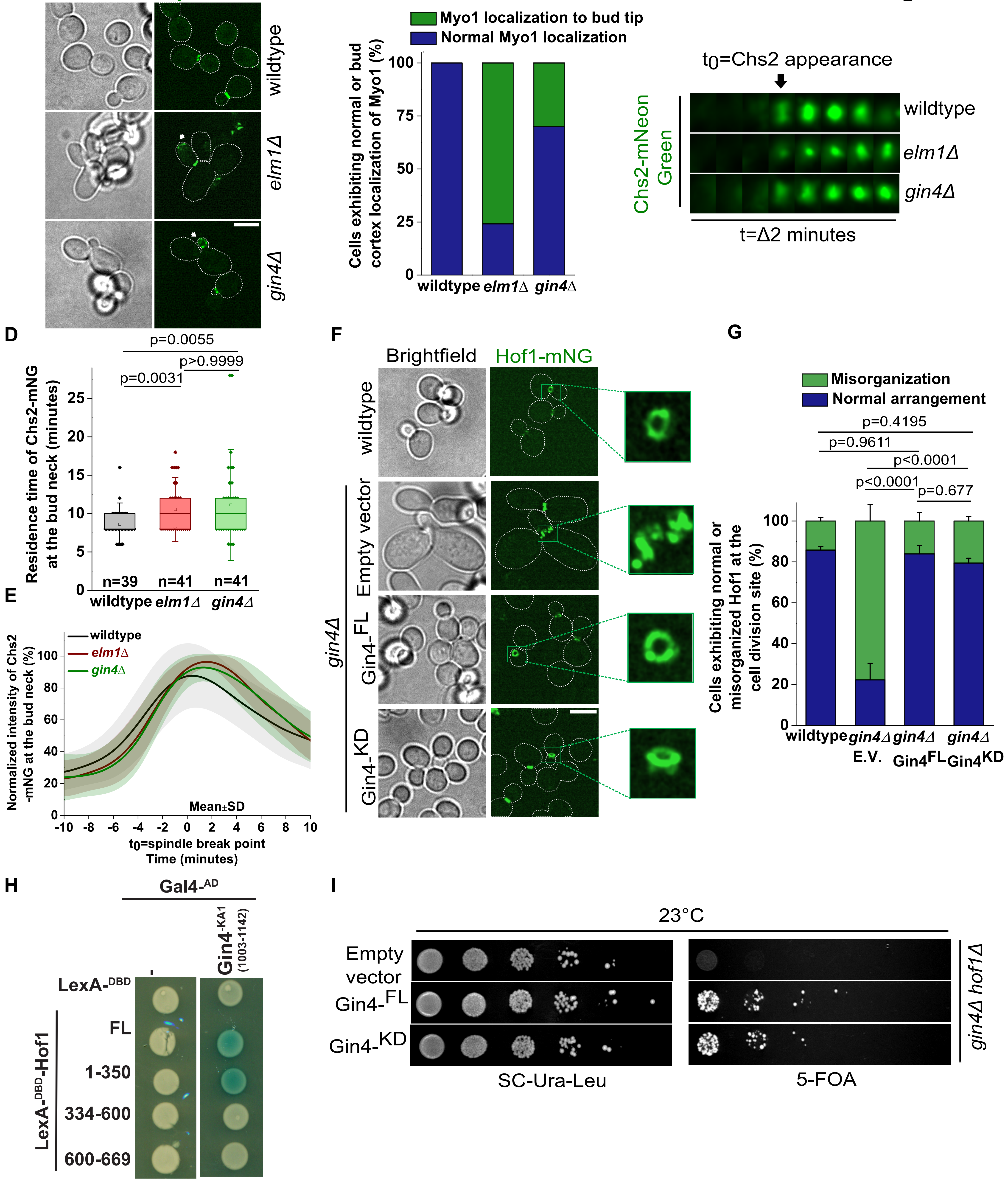
A
Brightfield
Myo1-mNG
wildtype
elm1Δ
gin4Δ
B
Myo1 localization to bud tip
Normal Myo1 localization
Cells exhibiting normal or bud cortex localization of Myo1 (%)
0
25
50
75
100
wildtype
elm1Δ
gin4Δ
C
t0=Chs2 appearance
Chs2-mNeon Green
wildtype
elm1Δ
gin4Δ
t=Δ2 minutes
D
p=0.0055
p>0.9999
p=0.0031
Residence time of Chs2-mNG at the bud neck (minutes)
n=39
n=41
n=41
wildtype
elm1Δ
gin4Δ
E
Normalized intensity of Chs2 -mNG at the bud neck (%)
wildtype
elm1Δ
gin4Δ
Mean±SD
t0=spindle break point
Time (minutes)
F
Brightfield
Hof1-mNG
wildtype
Empty vector
Gin4-FL
Gin4-KD
gin4Δ
G
Misorganization
Normal arrangement
p=0.4195
p=0.9611
p<0.0001
p<0.0001
p=0.677
Cells exhibiting normal or misorganized Hof1 at the cell division site (%)
wildtype
gin4Δ E.V.
gin4Δ Gin4FL
gin4Δ Gin4KD
H
Gal4-AD
Gin4-KA1 (1003-1142)
LexA-DBD
FL
1-350
334-600
600-669
LexA-DBD-Hof1
I
23°C
Empty vector
Gin4-FL
Gin4-KD
SC-Ura-Leu
5-FOA
gin4Δ hof1Δ

### Figure S4

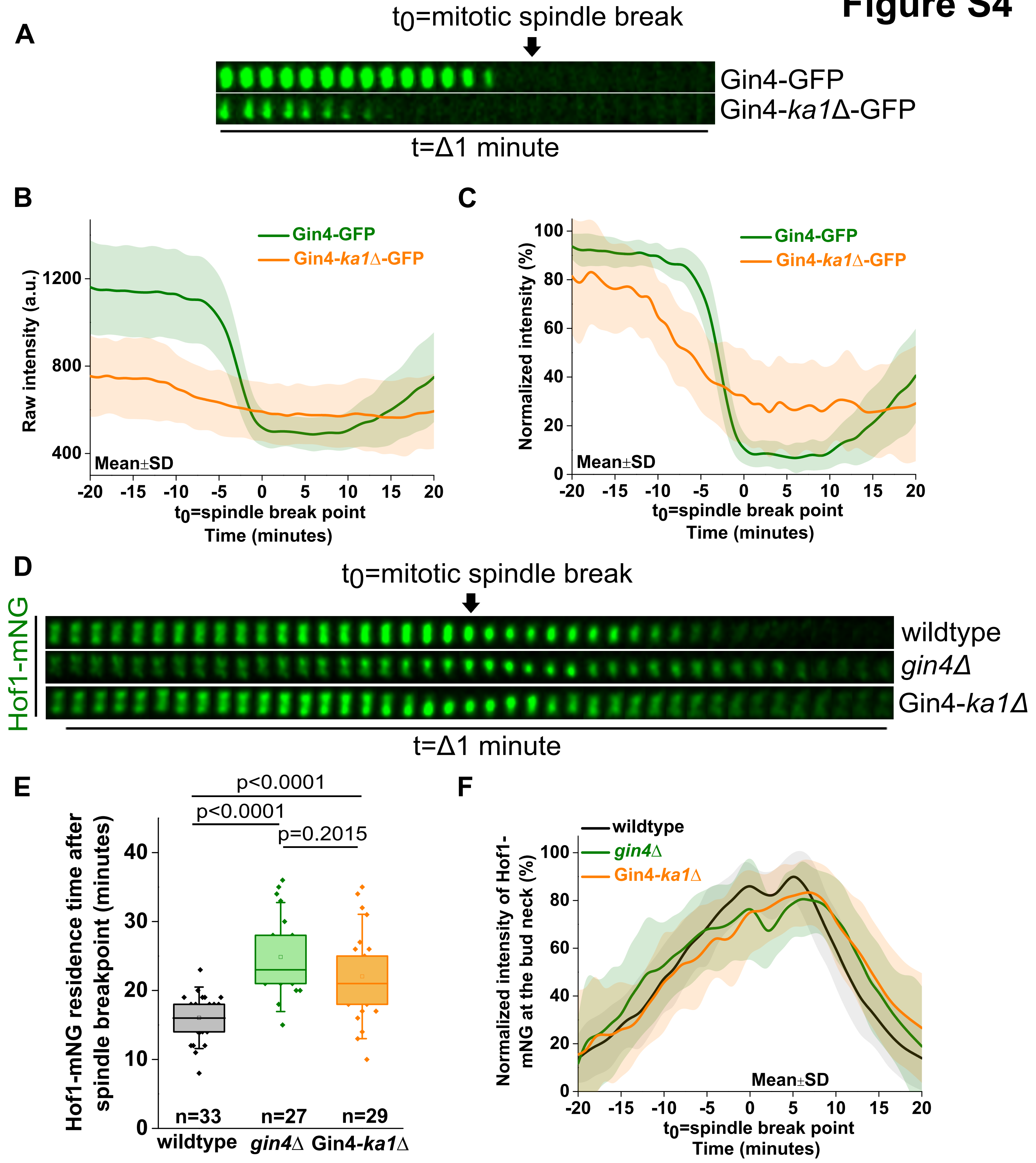
A
t0=mitotic spindle break
Gin4-GFP
Gin4-ka1Δ-GFP
t=Δ1 minute
B
Gin4-GFP
Gin4-ka1Δ-GFP
Raw intensity (a.u.)
1200
800
400
Mean±SD
-20 -15 -10 -5 0 5 10 15 20
t0=spindle break point
Time (minutes)
C
Gin4-GFP
Gin4-ka1Δ-GFP
Normalized intensity (%)
100
80
60
40
20
0
Mean±SD
-20 -15 -10 -5 0 5 10 15 20
t0=spindle break point
Time (minutes)
D
t0=mitotic spindle break
Hof1-mNG
wildtype
gin4Δ
Gin4-ka1Δ
t=Δ1 minute
E
p<0.0001
p<0.0001
p=0.2015
Hof1-mNG residence time after spindle breakpoint (minutes)
40
30
20
10
0
n=33
n=27
n=29
wildtype
gin4Δ
Gin4-ka1Δ
F
wildtype
gin4Δ
Gin4-ka1Δ
Normalized intensity of Hof1-mNG at the bud neck (%)
100
80
60
40
20
0
Mean±SD
-20 -15 -10 -5 0 5 10 15 20
t0=spindle break point
Time (minutes)

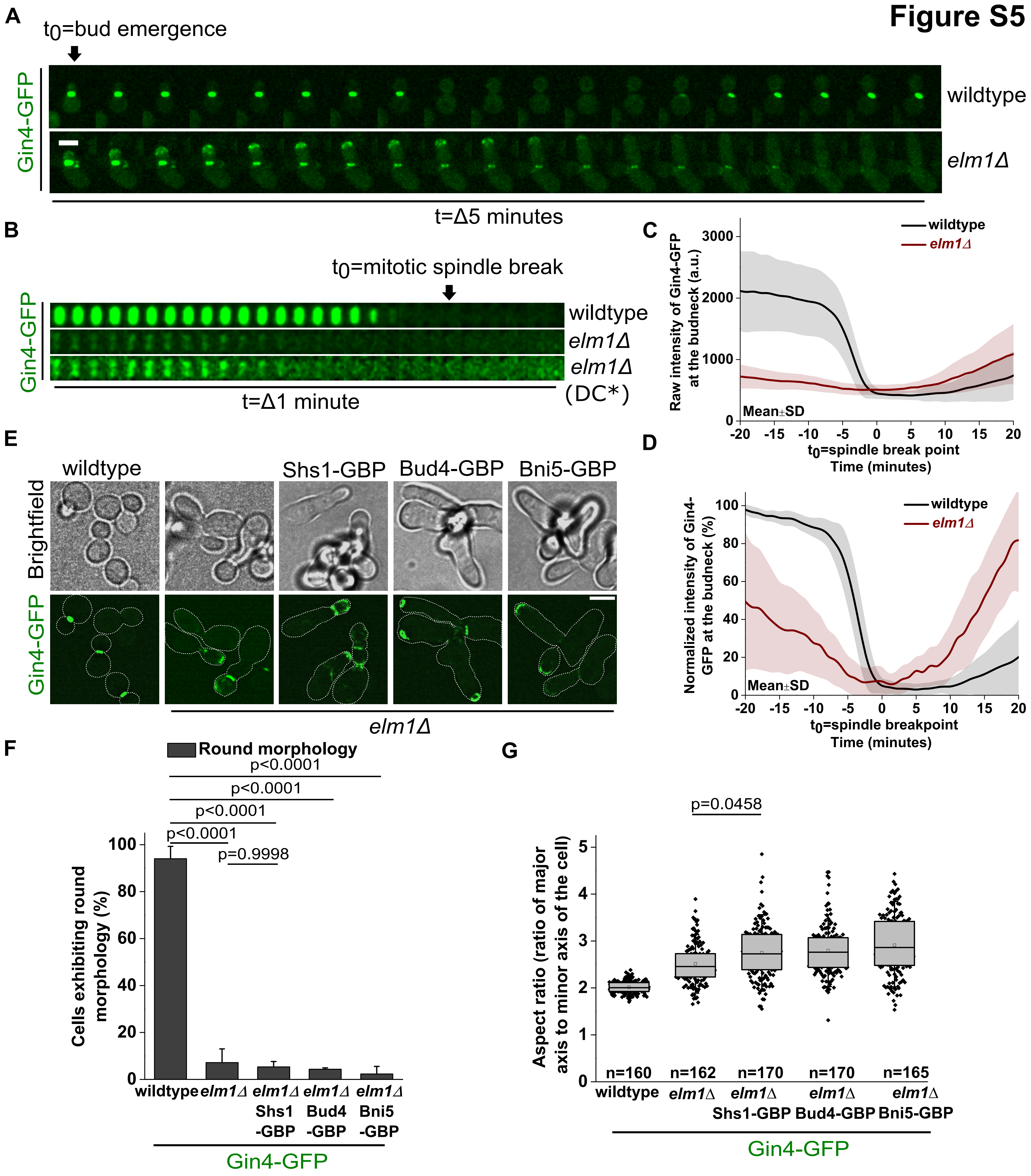

A
t0=bud emergence
Gin4-GFP
wildtype
elm1Δ
t=Δ5 minutes
B
t0=mitotic spindle break
wildtype
elm1Δ
elm1Δ
(DC*)
t=Δ1 minute
C
Raw intensity of Gin4-GFP at the budneck (a.u.)
wildtype
elm1Δ
Mean±SD
t0=spindle break point
Time (minutes)
D
Normalized intensity of Gin4-GFP at the budneck (%)
wildtype
elm1Δ
Mean±SD
t0=spindle breakpoint
Time (minutes)
E
wildtype
Shs1-GBP
Bud4-GBP
Bni5-GBP
Brightfield
Gin4-GFP
elm1Δ
F
Round morphology
p<0.0001
p<0.0001
p<0.0001
p<0.0001
p=0.9998
Cells exhibiting round morphology (%)
wildtype
elm1Δ
elm1Δ Shs1 -GBP
elm1Δ Bud4 -GBP
elm1Δ Bni5 -GBP
Gin4-GFP
G
p=0.0458
Aspect ratio (ratio of major axis to minor axis of the cell)
n=160
n=162
n=170
n=170
n=165
wildtype
elm1Δ
elm1Δ Shs1-GBP
elm1Δ Bud4-GBP
elm1Δ Bni5-GBP
Gin4-GFP


**Figure S5**

### Figure S6

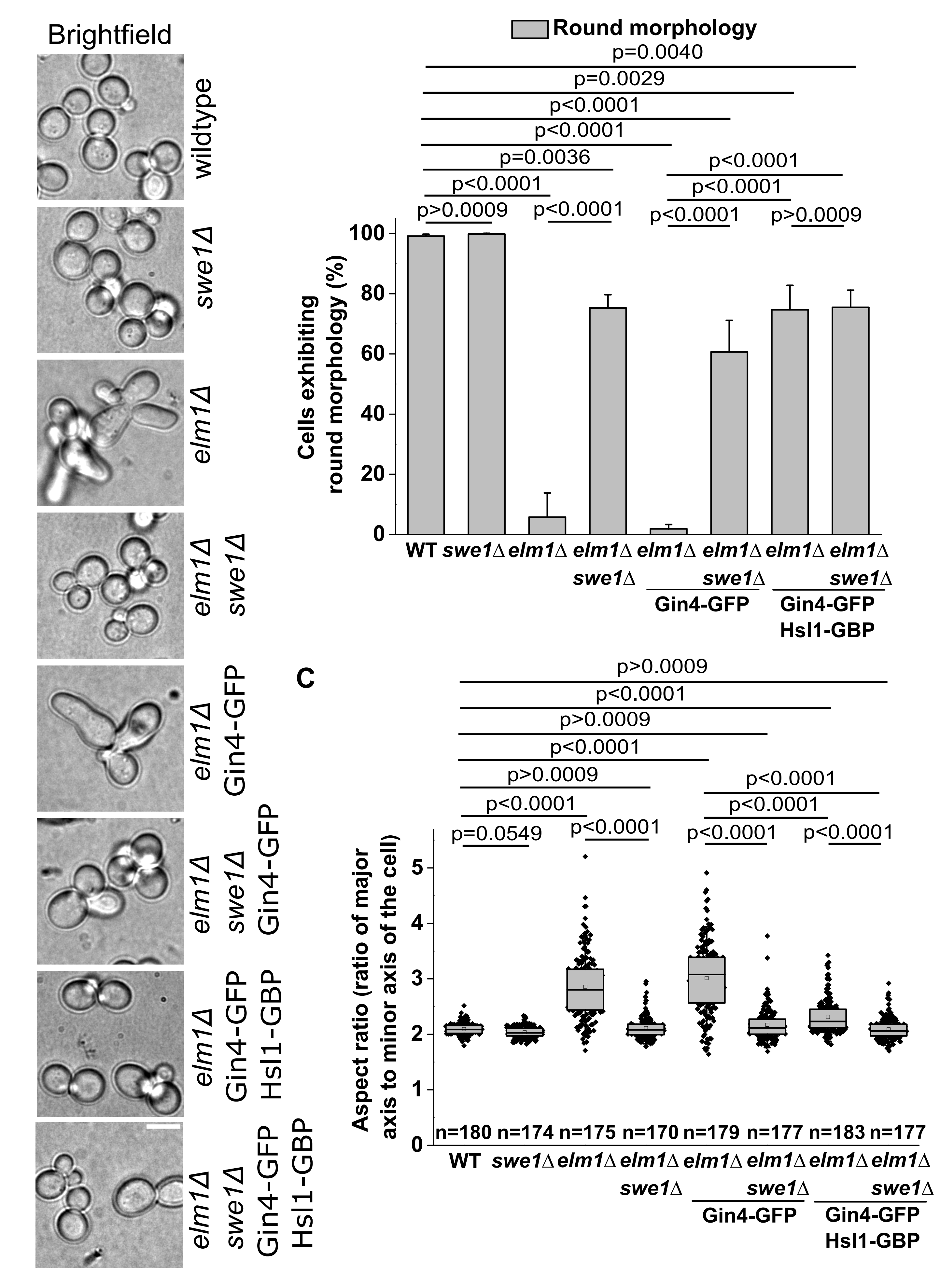

A
Brightfield
wildtype
swe1Δ
elm1Δ
elm1Δ swe1Δ
elm1Δ Gin4-GFP
elm1Δ swe1Δ Gin4-GFP
elm1Δ Gin4-GFP Hsl1-GBP
elm1Δ swe1Δ Gin4-GFP Hsl1-GBP
B
Round morphology
p=0.0040
p=0.0029
p<0.0001
p<0.0001
p=0.0036
p<0.0001
p>0.0009
p<0.0001
p<0.0001
p<0.0001
p<0.0001
p>0.0009
Cells exhibiting round morphology (%)
0
20
40
60
80
100
WT
swe1Δ
elm1Δ
elm1Δ swe1Δ
elm1Δ
elm1Δ swe1Δ
Gin4-GFP
elm1Δ
elm1Δ swe1Δ
Gin4-GFP Hsl1-GBP
C
p>0.0009
p<0.0001
p>0.0009
p<0.0001
p>0.0009
p<0.0001
p=0.0549
p<0.0001
p<0.0001
p<0.0001
p<0.0001
p<0.0001
Aspect ratio (ratio of major axis to minor axis of the cell)
0
1
2
3
4
5
n=180
n=174
n=175
n=170
n=179
n=177
n=183
n=177
WT
swe1Δ
elm1Δ
elm1Δ swe1Δ
elm1Δ
elm1Δ swe1Δ
Gin4-GFP
elm1Δ
elm1Δ swe1Δ
Gin4-GFP Hsl1-GBP

### Figure S7

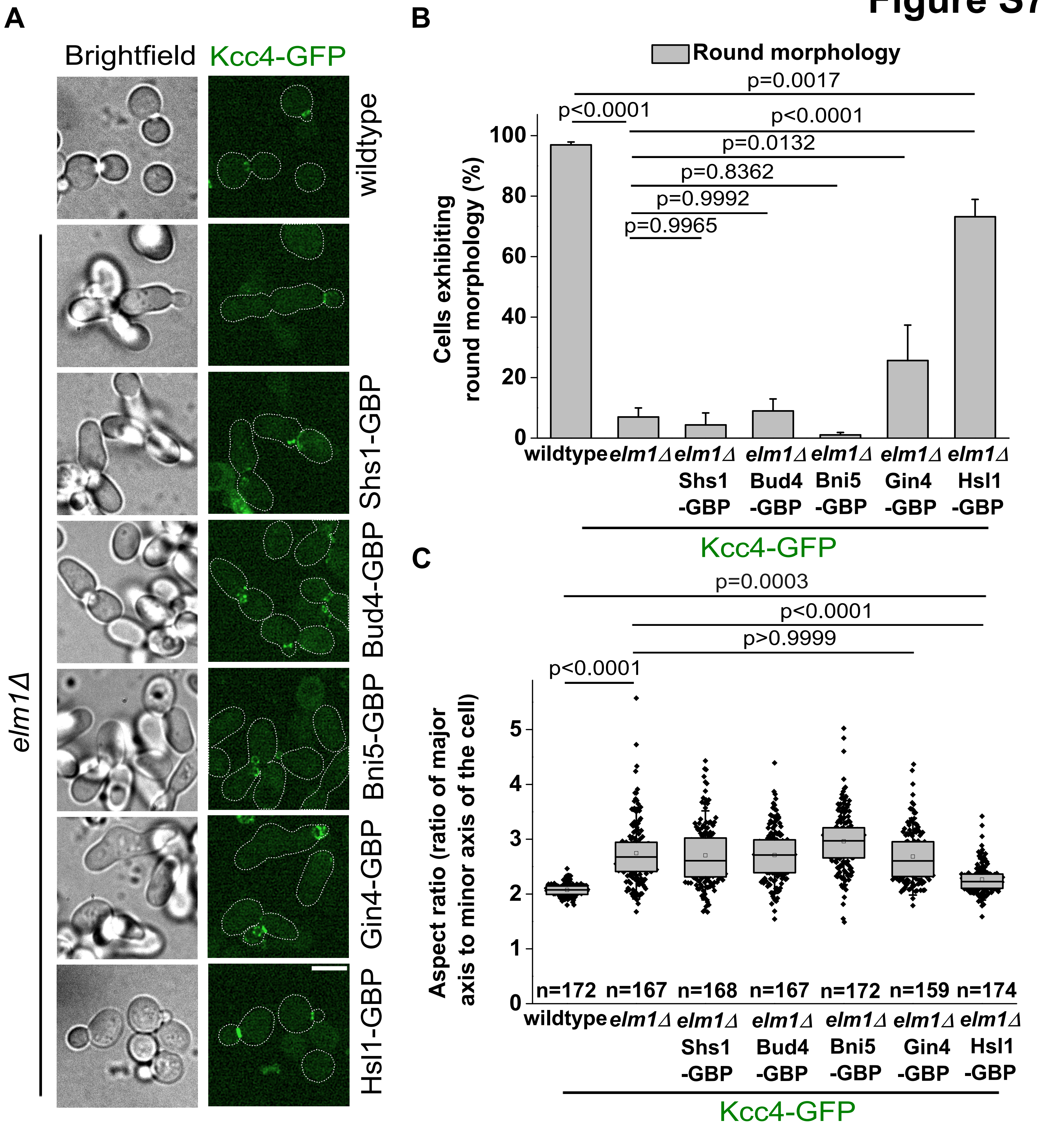
A
Brightfield
Kcc4-GFP
wildtype
Shs1-GBP
Bud4-GBP
Bni5-GBP
Gin4-GBP
Hsl1-GBP
elm1Δ
B
Round morphology
p=0.0017
p<0.0001
p<0.0001
p=0.0132
p=0.8362
p=0.9992
p=0.9965
Cells exhibiting round morphology (%)
0
20
40
60
80
100
wildtype
elm1Δ
elm1Δ Shs1 -GBP
elm1Δ Bud4 -GBP
elm1Δ Bni5 -GBP
elm1Δ Gin4 -GBP
elm1Δ Hsl1 -GBP
Kcc4-GFP
C
p=0.0003
p<0.0001
p>0.9999
p<0.0001
Aspect ratio (ratio of major axis to minor axis of the cell)
0
1
2
3
4
5
n=172
n=167
n=168
n=167
n=172
n=159
n=174
wildtype
elm1Δ
elm1Δ Shs1 -GBP
elm1Δ Bud4 -GBP
elm1Δ Bni5 -GBP
elm1Δ Gin4 -GBP
elm1Δ Hsl1 -GBP
Kcc4-GFP

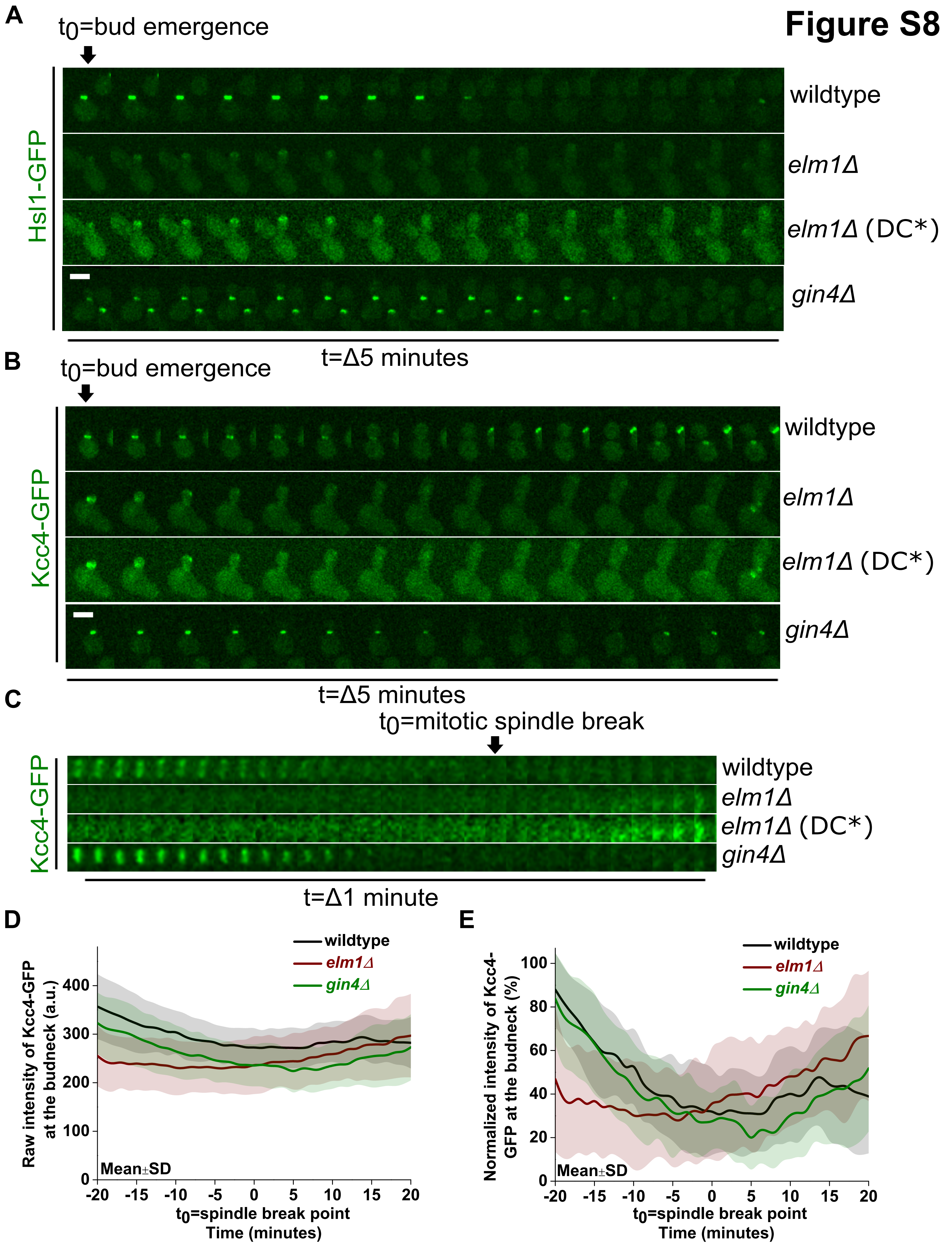
Figure S8
A
t0=bud emergence
Hsl1-GFP
wildtype
elm1Δ
elm1Δ (DC*)
gin4Δ
t=Δ5 minutes
B
t0=bud emergence
Kcc4-GFP
wildtype
elm1Δ
elm1Δ (DC*)
gin4Δ
t=Δ5 minutes
C
t0=mitotic spindle break
Kcc4-GFP
wildtype
elm1Δ
elm1Δ (DC*)
gin4Δ
t=Δ1 minute
D
wildtype
elm1Δ
gin4Δ
Raw intensity of Kcc4-GFP at the budneck (a.u.)
Mean±SD
t0=spindle break point
Time (minutes)
E
wildtype
elm1Δ
gin4Δ
Normalized intensity of Kcc4-GFP at the budneck (%)
Mean±SD
t0=spindle break point
Time (minutes)

### Figure S9

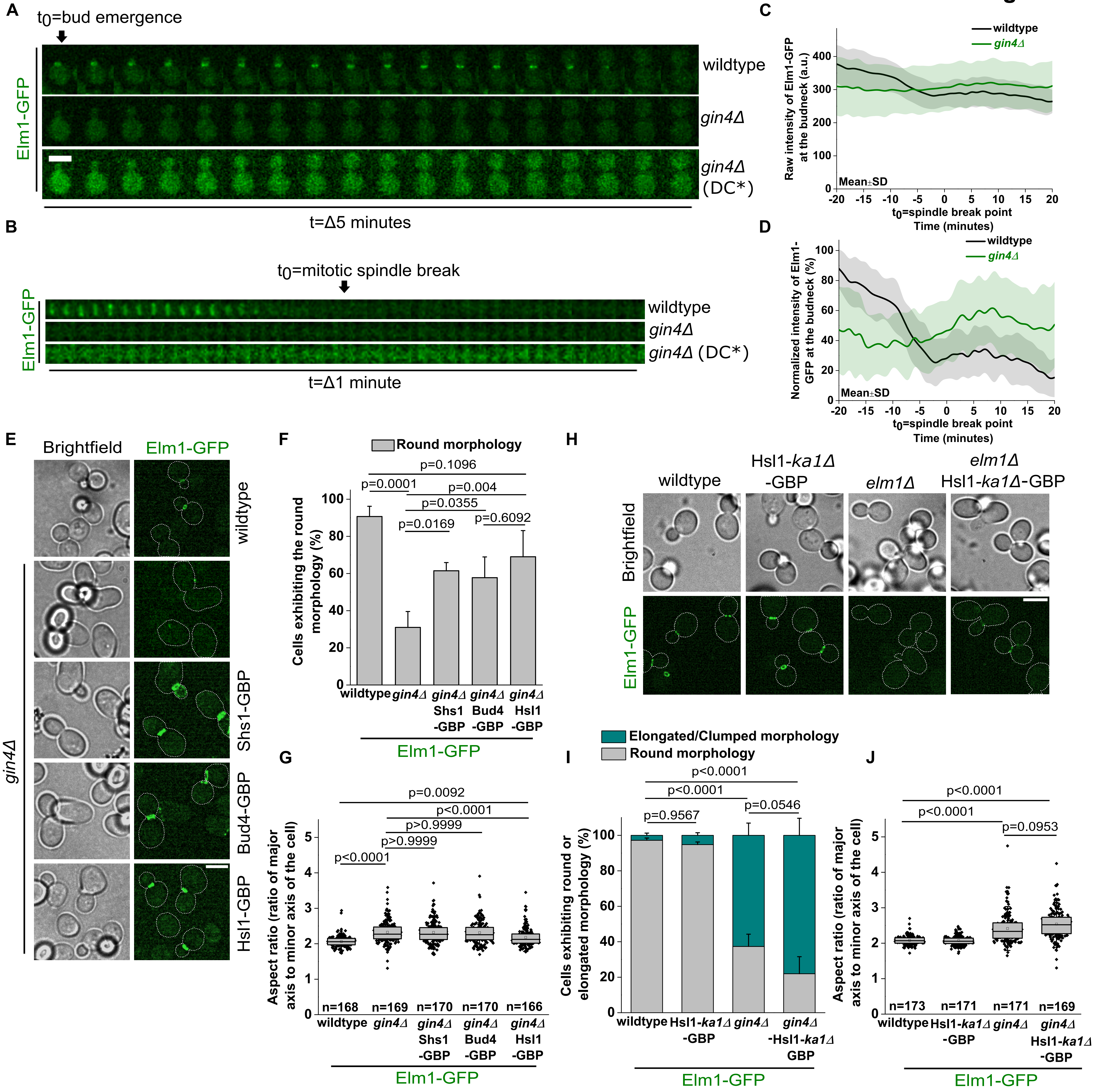

A
t0=bud emergence
Elm1-GFP
wildtype
gin4Δ
gin4Δ (DC*)
t=Δ5 minutes
B
t0=mitotic spindle break
Elm1-GFP
wildtype
gin4Δ
gin4Δ (DC*)
t=Δ1 minute
C
wildtype
gin4Δ
Raw intensity of Elm1-GFP at the budneck (a.u.)
Mean±SD
t0=spindle break point
Time (minutes)
D
wildtype
gin4Δ
Normalized intensity of Elm1-GFP at the budneck (%)
Mean±SD
t0=spindle break point
Time (minutes)
E
Brightfield
Elm1-GFP
wildtype
gin4Δ
Shs1-GBP
Bud4-GBP
Hsl1-GBP
F
Round morphology
p=0.1096
p=0.0001
p=0.004
p=0.0355
p=0.0169
p=0.6092
Cells exhibiting the round morphology (%)
wildtype
gin4Δ
gin4Δ Shs1 -GBP
gin4Δ Bud4 -GBP
gin4Δ Hsl1 -GBP
Elm1-GFP
G
p=0.0092
p<0.0001
p>0.9999
p>0.9999
p<0.0001
Aspect ratio (ratio of major axis to minor axis of the cell)
n=168 n=169 n=170 n=170 n=166
wildtype
gin4Δ
gin4Δ Shs1 -GBP
gin4Δ Bud4 -GBP
gin4Δ Hsl1 -GBP
Elm1-GFP
H
wildtype
Hsl1-ka1Δ -GBP
elm1Δ
elm1Δ Hsl1-ka1Δ-GBP
Brightfield
Elm1-GFP
I
Elongated/Clumped morphology
Round morphology
p<0.0001
p<0.0001
p=0.9567
p=0.0546
Cells exhibiting round or elongated morphology (%)
wildtype
Hsl1-ka1Δ -GBP
gin4Δ
gin4Δ -Hsl1-ka1Δ GBP
Elm1-GFP
J
p<0.0001
p<0.0001
p=0.0953
Aspect ratio (ratio of major axis to minor axis of the cell)
n=173 n=171 n=171 n=169
wildtype
Hsl1-ka1Δ -GBP
gin4Δ
gin4Δ Hsl1-ka1Δ -GBP
Elm1-GFP

### Figure S10

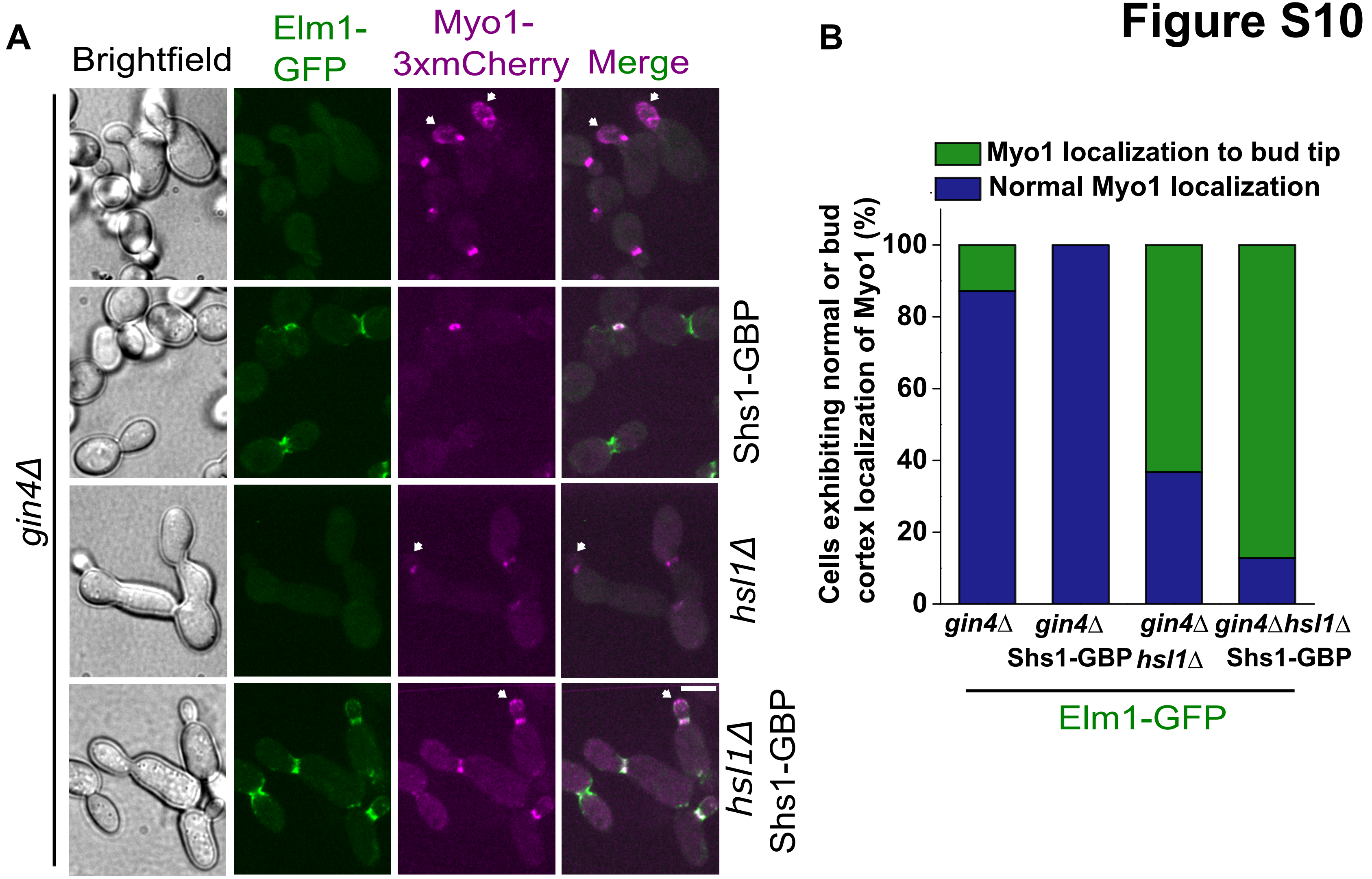
A
Brightfield
Elm1-GFP
Myo1-3xmCherry
Merge
gin4Δ
Shs1-GBP
hsl1Δ
hsl1Δ Shs1-GBP
B
Myo1 localization to bud tip
Normal Myo1 localization
Cells exhibiting normal or bud cortex localization of Myo1 (%)
0
20
40
60
80
100
gin4Δ
gin4Δ Shs1-GBP
gin4Δ hsl1Δ
gin4Δhsl1Δ Shs1-GBP
Elm1-GFP
